# Global homeostatic forces shape the composition of a hyperdiverse tropical insect fauna

**DOI:** 10.64898/2026.11.04.764147

**Authors:** Fredrik Ronquist, Ela Iwaszkiewicz-Eggebrecht, Laura J. A. van Dijk, Robert Goodsell, Bruno C. Bellini, Gavin R. Broad, Nerivania N. Godeiro, Emma Granqvist, Tilia Häggström, John Heraty, Seraina Klopfstein, Greg P. A. Lamarre, Fabian A. Lundell, John Sundh, Claudia C. Weber, Dimby Raharinjanahary, Eric T. Rajoelison, Johannes Bergsten, Brian L. Fisher, Anders F. Andersson, Piotr Łukasik, Andreia Miraldo, Tomas Roslin, Ayco J. M. Tack

## Abstract

Despite comprising the bulk of terrestrial biodiversity, we know surprisingly little about insect faunas, particularly in the tropics. Here, we characterize the insect fauna of Madagascar, a global biodiversity hotspot, based on extensive DNA metabarcoding of Malaise trap and leaf litter samples. We estimate that the fauna comprises at least 250,000 species, only 5% of which are currently recorded. The ecological and taxonomic composition is surprisingly similar to that of temperate forests. We show that this is likely caused by global homeostatic forces shaping net diversification rates after colonization of Madagascar rather than by the composition of colonizing lineages. Features differing from temperate forests include the dearth of parasitoids and richness of beetles. Understanding the forces shaping these patterns is key in sustainable management of tropical insect faunas.

---

Insects make up a major fraction of terrestrial ecosystems both in terms of species richness and biomass. Yet, we still know surprisingly little about the size and composition of the insect faunas of the planet. This is especially true for the tropics, where the impressive diversity constitutes a daunting challenge, particularly in neglected but hyperdiverse ‘dark taxa’ (*1*). A better understanding of how insect faunas are structured and what shapes them would provide essential insights into the likely responses of tropical ecosystems to environmental change. This is an urgent priority at a time of unprecedented pressures from a warming climate, changing land use, and other stressors (*2*, *3*).

An insect fauna can be decomposed along different axes, providing complementary insights into the forces shaping it (*4*). For tropical faunas where the majority of species have evolved in place, the taxonomic composition describes how different lineages have diversified into the available resource space, in interaction with other lineages and the local environment (*5*). In contrast, temperate faunas are typically assembled from species that evolved elsewhere, and have been reshuffled by recent glaciations (*6*). A different perspective is provided by the faunal composition in terms of feeding niches and microhabitats, as similar functional roles have evolved repeatedly across numerous lineages. Finally, patterns of species turnover reveal the faunal dynamics in space and across seasons.

Current estimates of global insect diversity are often based on compositional assumptions with shaky empirical support, such as the size of insect faunas being proportional to that of vascular plant floras across biomes and latitudes (*7*, *8*). It has also been common to extrapolate features of better-known temperate faunas, such as the parasitoid to host species ratio, to largely unexplored tropical faunas (*9*, *10*). A recent study supports the idea of insect faunas having similar taxonomic composition across the globe (*1*), while another emphasizes the differences by suggesting that latitudinal diversity gradients are weak or reversed in many of the hyperdiverse taxa (*11*). The Ichneumonidae, the largest family of parasitic wasps, provide a famous example of a reversed gradient (*12*), attributed to more fragmented host resources in the tropics (*13*) or better-protected (“nastier”) hosts containing more toxic compounds (*14*). Currently, it seems unclear whether the reversed gradient applies to all parasitoids (*9*, *15–19*) and, if it does, why (*11*, *20*).

Previous attempts to analyze the composition of tropical insect faunas have focused on small areas and a limited set of taxonomic groups (*8*, *21*). In recent years, DNA metabarcoding has opened up the possibility to survey entire insect faunas at scale (*22–24*). Here, we take advantage of these methods in characterizing the insect fauna of Madagascar, one of the foremost global biodiversity hotspots.

The fourth-largest island in the world, Madagascar is well known for its high level of endemic diversity (*25–28*). Madagascar separated from Africa about 170 million years ago (Ma), and from India about 88 Ma (*29*, *30*). The long period of relative isolation, in combination with a complex geography and significant climatic changes (*31–33*), has produced a unique fauna and flora shaped by a mix of historical contingencies, local radiations, phylogenetic inertia and filtering by environmental, climatic and biotic factors. Thus, the Madagascan fauna provides the ultimate test of the existence of global patterns in the composition of insect faunas, and a key to the causal factors shaping them.

Our study is based on metabarcoding of 1,919 Malaise trap samples and 185 leaf litter samples collected over 12 months at 50 forest sites across Madagascar in the Insect Biome Atlas (IBA) project (*24*) (fig. S1). The sites include 25 wet forest sites at 14 locations in the east, 2 montane forest sites at different central locations, and 23 dry forest sites at 17 locations spread through the rest of the island. We collectively refer to the wet forest and montane forest as “rainforest”.

We metabarcoded a 418 bp stretch of cytochrome c oxidase subunit I. The operational taxonomic units (OTUs)—hypothetical species—resulting from noise filtering and clustering of the data were placed in a reference tree developed for this study, with expanded coverage of dark taxa and comprising 48,313 insect species representing 854 insect families. Ecological composition of the fauna was studied by using evolutionarily conserved life-history traits coded at the subfamily level (the hyperdiverse Cecidomyiidae, Braconidae and Ichneumonidae) or family level (other insects).

We assessed the accuracy of our results through extensive validation tests using the similarly designed IBA survey of the well-studied insect fauna of Sweden (*24*). We also use the Swedish IBA data from forest sites, spanning the nemoral to boreal transition, as a suitable reference for comparing the insect fauna of Madagascan forests to that of temperate forests. We note that the countries are of similar size, and that Sweden represents an extreme contrast to Madagascar in that its insect communities were assembled very recently, after the last glacial maximum (*34*). Given their completely different origins, similarities between the faunas would provide strong evidence for global stabilizing forces affecting the composition. The phylogenetic placement of sampled OTUs, in combination with the long isolation of Madagascar, offer additional possibilities to untangle the role of source faunas and subsequent endemic radiation in shaping the composition of the Madagascan insect fauna.

## Results

### Species richness

Our Madagascan dataset comprises 55,478 OTUs, far exceeding the currently described insect fauna. The OTUs belong to 548 different family or subfamily clades, and most of them are undescribed. Only 4.5% of them were identified to species level when matching sequences to the BOLD reference library with SINTAX (the corresponding fraction for the Swedish data was 46%).

Species accumulation curves for the Madagascan data increase almost linearly, indicating that the total species pool is considerably larger than the number of sampled species (Fig. 1A). The Chao2 estimate—based on the frequency of singleton and doubleton occurrences—increases more rapidly than the number of species regardless of sampling effort, showing that even the last sites increase the proportion of rare species. This contrasts with the temperate forest reference data, for which a similar effort results in accumulation curves and Chao2 estimates that clearly approach an asymptote (Fig. 1B).

**Fig. 1.**
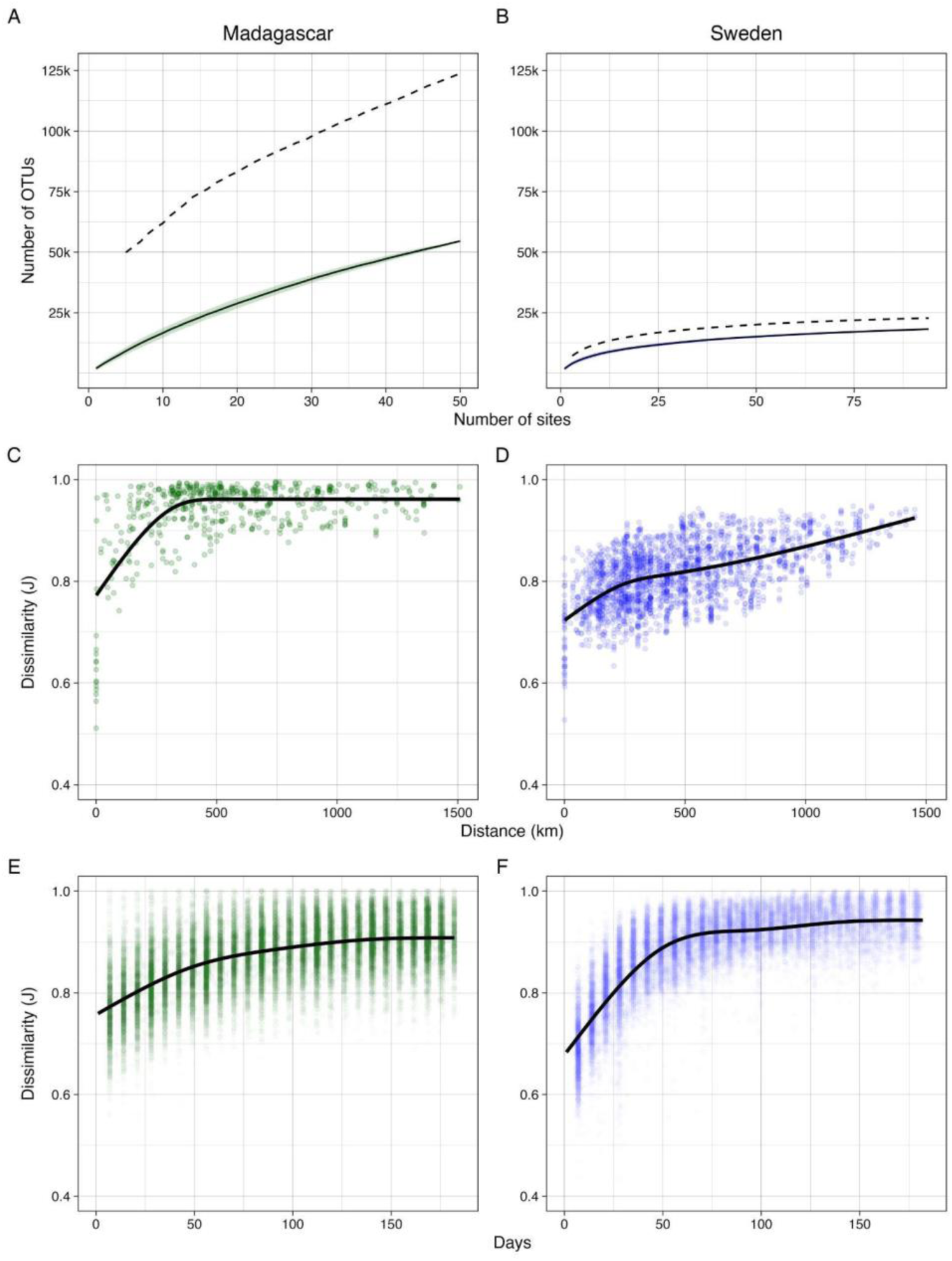
Species richness and turnover of the insect fauna of Madagascan forests, compared to that of temperate reference forests (Sweden). **(A–B)** Species accumulation with sampling effort (trap sites). The solid lines illustrate richness as a function of sampling effort and are based on the mean of 100 permutations; the shaded regions around them mark values within one standard deviation from the mean. The dashed line represents average Chao2 estimates of total richness, computed from randomly permuted data. The estimate is based on the frequency of singleton and doubleton occurrences; it increases more rapidly than the observed number of species as long as the proportion of rare species in the sample increases. **(C–F)** Turnover (the species replacement component of beta diversity). Turnover is expressed as Jaccard dissimilarity between sample pairs separated in space or time. The fitted line is generated from a monotonic generative additive model with five knots. **(C–D)** Spatial turnover. **(E–F)** Temporal turnover within trap sites.

The Chao2 estimator is known to provide a lower bound on total species richness (*35*, *36*). Benchmarks suggest that at least half the species need to be sampled before the Chao2 estimates are likely to be within a factor 2 of the true value (*37*). This indicates that the Madagascan species pool is well above 250,000, double the Chao2 estimate (Fig. 1A).

The species richness of Madagascan forests is linked to higher spatial turnover (species replacement over space) than in the temperate reference forests (Fig. 1C, D). In contrast, temporal turnover (species replacement through time) is lower in Madagascar than in temperate forests (Fig. 1E, F).

### Origins and spatial distribution

In terms of overall patterns, species richness and diversity tend to be higher in the rainforest than in the dry forest (Fig. 2A, B), although there is considerable variation among sites in both forest types. As might be expected from the extreme spatial turnover, few OTUs are shared across sites: average Bray-Curtis distances to other sites range from 0.95 to 1.00 (Fig. 2C). The spatial turnover is slightly higher and the temporal turnover slightly lower in the rainforest than in the dry forest (fig. S2).

**Fig. 2.**
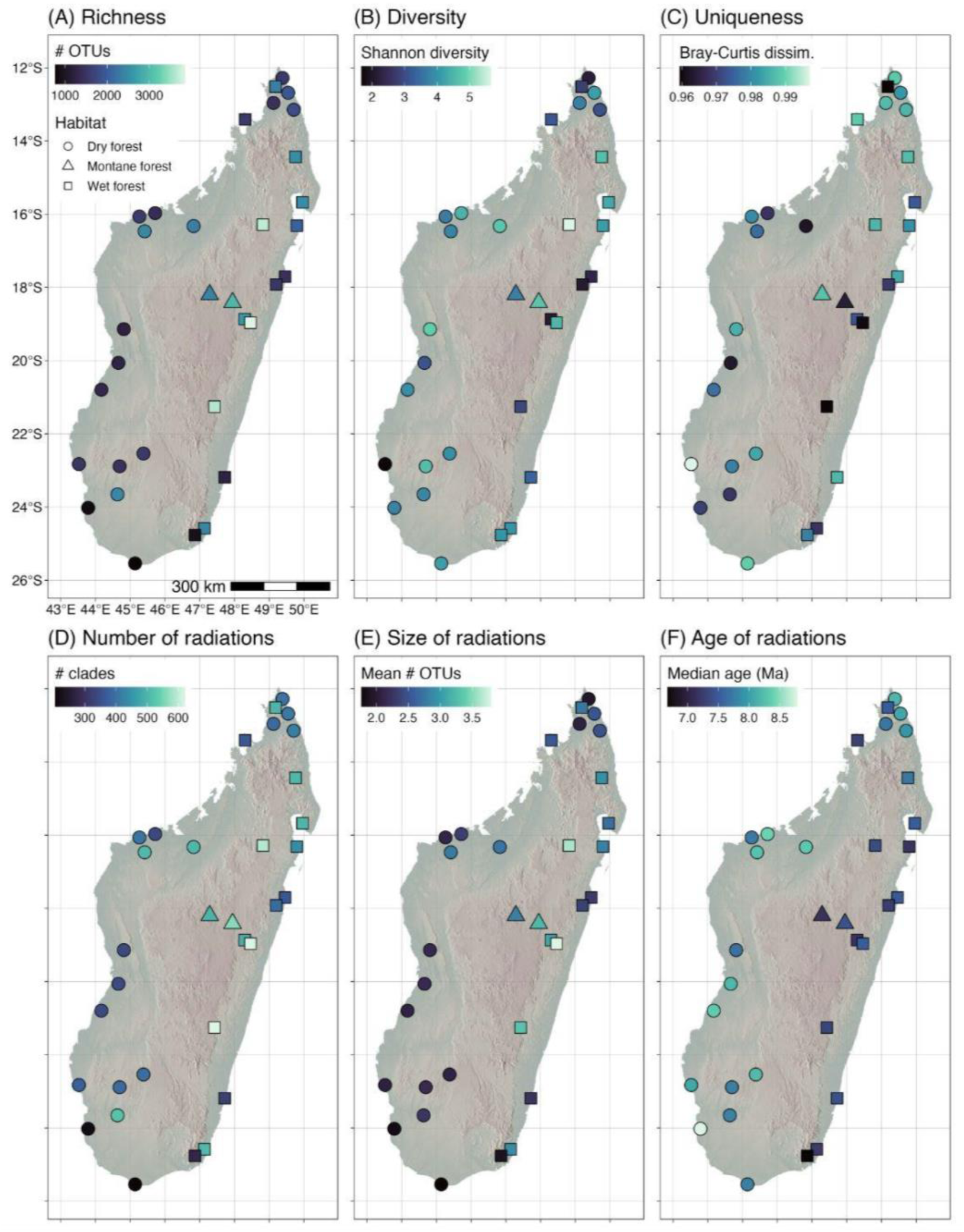
General characteristics of the Madagascan insect fauna as documented by metabarcoding of Malaise trap and litter samples. (**A**) Species richness. (**B**) Diversity (Shannon index). (**C**) Uniqueness (average Bray-Curtis distance to other sites). (**D**) Number of endemic Madagascan radiations (each consisting of ≥ 5 sampled species) represented at each site. (**E**) Mean size of Madagascan radiations (number of OTUs). (**F**) Median age of Madagascan radiations (Ma). For multitrap locations, we show the median value across traps. In analyses by forest type (Fig. 3), we grouped montane forest and wet forest as “rainforest”.

Of the 55,478 OTUs, 40,375 could be placed unambiguously on one of the branches of the reference tree. They document a minimum of 6,476 independent colonizations of Madagascar; we refer to the 1,325 of these that resulted in at least five Madagascan species detected in our study as “endemic radiations”. The total number and average size of endemic radiations tend to be higher in the rainforest, although some dry forest sites approach similar characteristics (Fig. 2D, E). The median estimated age of the endemic radiations is higher in the western and northern parts of the country (Fig. 2F), reflecting a larger proportion of old radiations. However, in terms of absolute numbers, the rainforest sites in the east harbor larger numbers of both older (“paleoendemic”) and younger (“neoendemic”) radiations, regardless of whether the boundary between old and young is set at the end of the Oligocene (23 Ma) or Eocene (34 Ma) (fig. S3).

### Ecological and taxonomic composition

Despite the high degree of spatial turnover and associated uncertainty in the estimate of total species richness, our sample gives stable estimates of the ecological and taxonomic composition of the fauna, as indicated by both spatial and temporal accumulation curves (figs S4–S9). Overall, the composition of the Madagascan fauna is surprisingly similar to that of the temperate forest reference fauna (Fig. 3). Note that the accumulation of data across samples (yellow diamonds in Fig. 3; see also figs S4–S9) often results in additional convergence among regions and forest types, suggesting that the true faunas are even more similar than indicated by the distribution of 12-month site samples (box plots in Fig. 3).

**Fig. 3.**
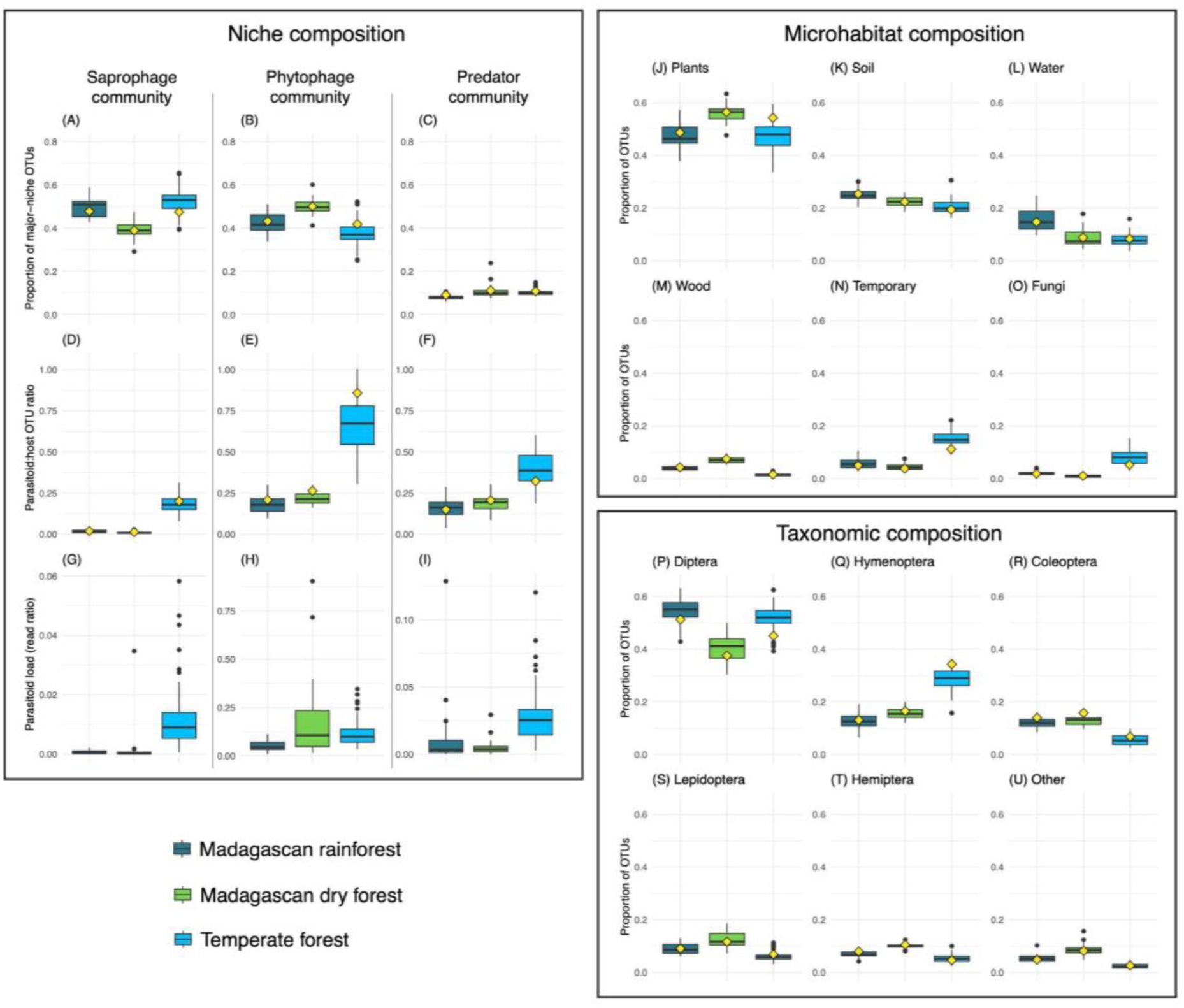
Ecological and taxonomic composition of the Madagascan insect fauna. The Madagascan data are shown by forest type: rainforest (montane forest and wet forest) and dry forest. The temperate reference forest data are based on the Swedish IBA survey. Boxplots summarize the distribution of values across 12-month samples from each trap site. The diamonds indicate the values to which the accumulation curves converge (Figs S4–S9), and the distance between each diamond and the mean of the corresponding box plot indicates the rate at which the value changes with additional spatial sampling. (**A–I**) Niche composition. (A–C) Proportion of OTUs belonging to the three major feeding niches. (D–F) Parasitoid to host OTU ratios for each of the three communities. (G–I) Parasitoid load for each of the three communities, estimated as the ratio between total calibrated read numbers for parasitoids and hosts in each community. (**J– O**) Microhabitat composition. “Plants” refer to species developing in or on plants; “soil” to species in soil, litter or on the ground; “water” to species in water; “wood” to species in wood; “temporary habitats” to species in carrion, dung, etc; and “fungi to species in fungi. (**P–U**) Taxonomic composition.

Focusing on the three major feeding niches (saprophages, phytophages and predators), insect communities are dominated by the two former, while predators (excluding parasitoids) account for approximately 10% of the OTUs (Fig. 3A–C). The ratio of phytophages to saprophages is slightly higher in the dry forest than in the rainforest but in both cases similar to the temperate forest. The parasitoid to host species ratio is higher in the phytophage community than in the saprophage and predator communities in both tropical and temperate forests (Fig. 3D–F). However, the relative contribution of parasitoids to the total species pool is strikingly small in Madagascar.

Using calibrated read numbers as a proxy for biomass, our data indicate that parasitoid load—the ratio of parasitoid to host biomass—in the phytophage community is higher than the parasitoid to host species ratio would suggest, with the dry forest being comparable to that of the temperate reference forest (Fig. 3H). This suggests that generalist parasitoids are more common in Madagascar. The spatial turnover of phytophage-parasitoids is similar to that of their hosts (fig. S10), indicating that Madagascan parasitoids attack a wider range of local hosts rather than tracking related host species that replace each other across space. The parasitoid load does not appear to be elevated compared to the parasitoid to host species ratio in the saprophage and predator communities, but the numbers are small, making conclusions uncertain (Fig. 3G, I).

In terms of main feeding microhabitats (Fig. 3J–O), most Madagascan insects develop in or on plants, followed by soil, water, wood, temporary habitats and fungi. This is quite similar to the temperate reference forest, with the most notable difference being a higher proportion of temperate species developing in or on temporary substrates (carrion, dung, etc.) or in fungi. The variations between rainforest and dry forest are small.

Taxonomically, the sampled Madagascan fauna is dominated by Diptera, followed by Hymenoptera, Coleoptera, Lepidoptera, and Hemiptera (Fig. 3P–U). Both the temporal (fig. S9) and spatial (fig. S6; see also diamonds in Fig. 3P–U) accumulation of species suggests that the proportions of Coleoptera and Hymenoptera are underestimated in the sample, while the proportion of Diptera is overestimated, although these effects are minor. The most striking difference to the temperate reference data is that the Hymenoptera constitute a considerably smaller portion of the species pool. This is clearly linked to the lower diversity in Madagascan forests of parasitoids, most of which are hymenopterans. The Diptera comprise a similar fraction of both faunas, while the Coleoptera are substantially more diverse in Madagascar. The taxonomic composition is similar in the two Madagascan forest types, with the most notable difference being a higher proportion of Diptera OTUs in the rainforest than in the dry forest (Fig. 3P).

Focusing on the more fine-grained family level reveals similar mixtures in the rainforest and dry forest (Fig. 4C, D) despite limited overlap at the OTU level (Fig. 4A). Most endemic radiations are shared, however (Fig. 4B). Among the top 20 families in terms of number of OTUs, 18 are shared between forest types. The gall midges (Diptera: Cecidomyiidae) dominate hugely in both forest types. A notable entry is the Mordellidae (tumbling flower beetles; 772 OTUs, none of which can be assigned to named species), a poorly studied but apparently diverse taxon in the canopy of tropical forests (*38*).

**Fig. 4.**
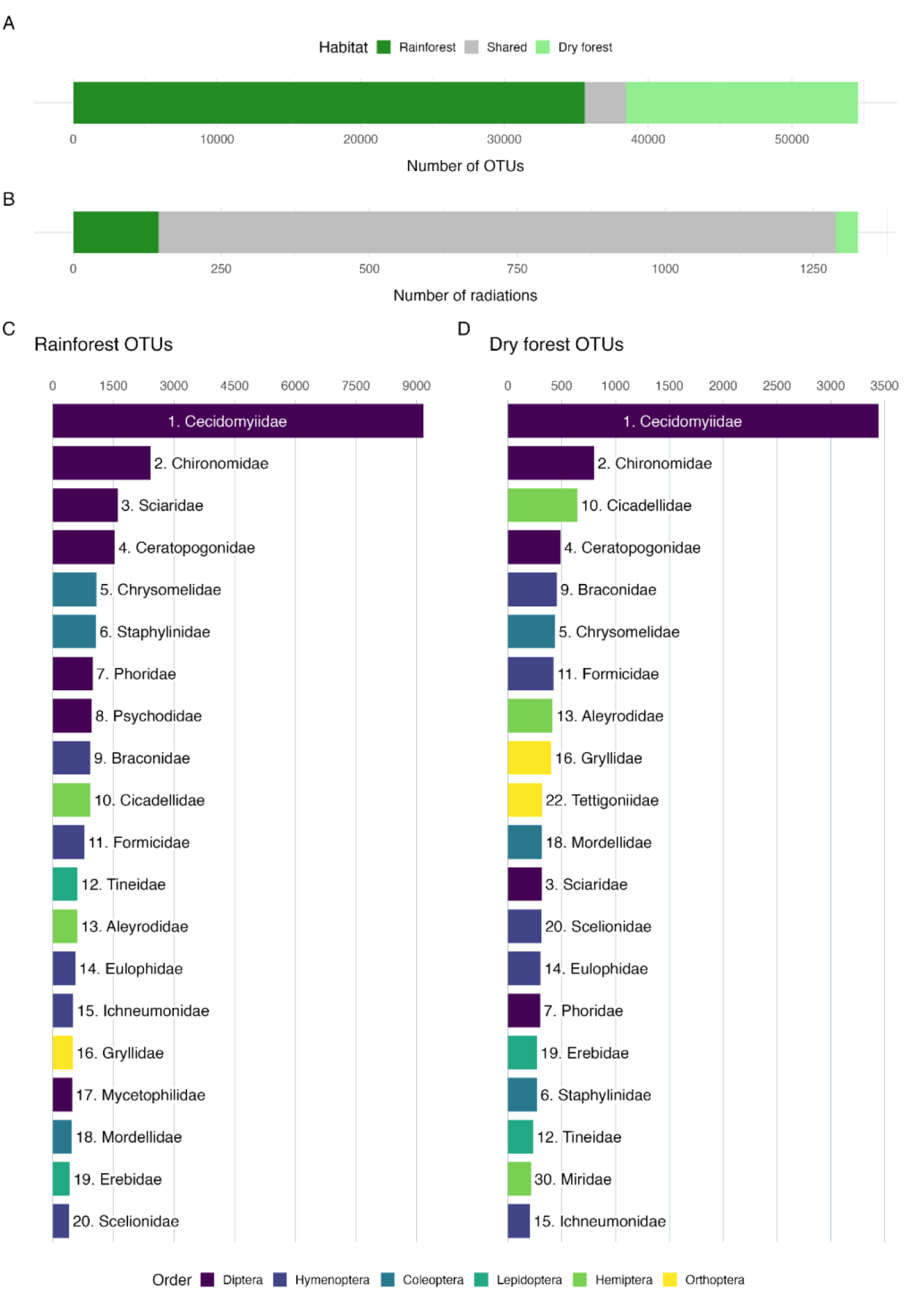
Dominant insect families in the Madagascan insect fauna. (**A**) Decomposition of the retrieved OTUs into those occurring only in rainforest or dry forest, and those shared between the habitats. (**B**) Similar decomposition of the endemic Madagascan radiations (≥ 5 sampled species). (**C**) Top 20 dominant families in the rainforest, ranked by the number of OTUs placed in them. (**D**) Ditto for the dry forest. Numbers in C–D refer to the rank of each family in rainforest sites.

When the data are broken into subfamilies, four separate gall-midge taxa still make it to the top 20 in Madagascan forests (fig. S11). Despite the similarities in ecological composition between the Madagascan and temperate forests, the differences in taxonomic composition are substantial at the family or family clade (Cecidomyiidae, Braconidae and Ichneumonidae broken into subfamilies) level (figs S11–S12). Only 12 of the top 20 families in the temperate reference forest are shared with Madagascan forests, and gall midges are much less dominant, even though they remain the most species-rich family (fig. S12).

### Origin of compositional patterns

Using neutral (lineage-agnostic) diversification as a null model, we analyzed the relative influence of colonizer composition and subsequent radiation on the Madagascan faunal composition (Fig. 5). These tests suggest that forces acting on the radiation of lineages after colonization have driven several features to be more similar to those of temperate forests than they were originally. In particular, this is true for the relative diversity of saprophages, phytophages and predators, and the taxonomic composition at the order and family levels (Fig. 5A). Microhabitat use has moved in the opposite direction relative to what would have been expected from neutral diversification, indicating active environmental filtering in response to unique Madagascan features during the subsequent radiation. The proportion of parasitoids is slightly but significantly lower than expected (p=0.0027), while the proportion of beetles matches expectations (Fig. 5B). In both cases, the expected composition under neutral diversification is quite different from temperate faunas, potentially reflecting the composition of source faunas.

**Fig. 5.**
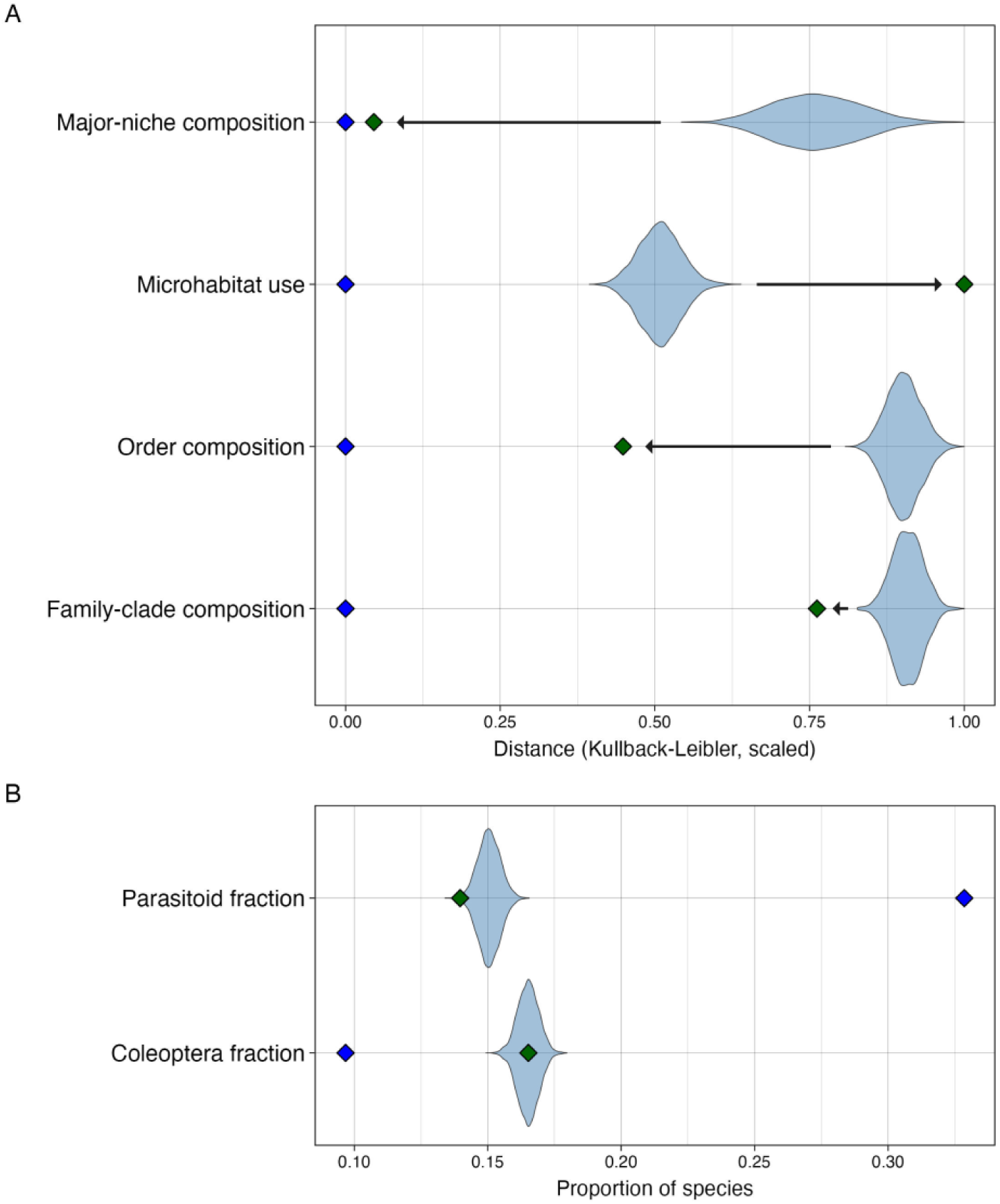
Diversification analysis of compositional features. We fitted a common time-dependent diversification model to all Madagascan radiations, conditioning on the age and the number of sampled OTUs of each radiation, and compensating for the sampling bias against beetles (Coleoptera) documented in the validation tests. By simulating data from the fitted model, we tested whether observed compositional features (green diamonds) deviated from the expectations under neutral diversification of colonizing lineages and, if so, whether the composition evolved (arrows) to become more similar or more different to that of temperate forests (blue diamonds). The simulations (violin plots) are likely to reflect compositional features of source faunas filtered through partly stochastic and partly deterministic selection of successful colonizers. (**A**) Compositional features measured in terms of relative Kullback-Leibler distance to temperate forests. (**B**) Fraction of parasitoids and fraction of beetles among non-parasitoids. Results were similar without the correction for Coleoptera sampling bias, except for the analysis of the beetle fraction (fig. S13).

## Discussion

The spectacular diversity and spatial turnover of the Madagascan insect fauna present unique surveying challenges. The Madagascan Malaise trap material analyzed here is estimated to contain 1.7 million insect specimens (*24*). Such an effort would be sufficient to establish an upper bound on the size of the species pool of a comparable temperate region, as shown by our reference data, but the nearly linear species accumulation curve and Chao2 estimates indicate that our inventory is not nearly enough to reliably estimate the total size of the Madagascan fauna. Until more robust estimates can be obtained, we suggest that a quarter of a million species may be taken as a very conservative lower bound.

The large number of endemic insect radiations identified in our study highlights the numerous lineages that have successively colonized Madagascar. The large proportion of old radiations in the north and west is consistent with the island being colonized mainly from the African mainland (*39*). As these communities became saturated, new lineages were presumably forced to disperse to the south and east. At the same time, the large absolute number of old insect radiations in the east suggests that these habitats are old, contradicting the narrative of a uniformly arid early Cenozoic Madagascar (*32*) but matching paleoendemic plant diversity patterns (*40*). Mesic forests similar to some of today’s humid and subhumid highland forests were likely present already at the K/P boundary (*33*) and could have served as refugia and centers of radiation for insect lineages adapted to such habitats (*41*). As conditions arose later for the modern orographic, vertically stratified, rainforests of the east (*32*, *33*), they provided exceptional potential for further radiation as documented by the numerous young endemic clades and the rich species diversity encountered there.

Our results demonstrate a number of striking similarities between the insect faunas of Madagascan and temperate forests. They have almost identical fractions of phytophagous, saprophagous and predatory species, and there are clear similarities in their taxonomic composition at the order and family levels. The neutral diversification tests suggest that this is not a chance occurrence. Instead, the similarities appear to be the result of stabilizing homeostatic forces that have shaped the radiation of insect lineages on Madagascar, pushing these aspects of faunal composition towards a stable global pattern. The underlying mechanism may be environmental and biotic similarities among forests across the planet, combined with phylogenetic constraints on how different insect lineages can divide up the available resources as they radiate.

Against this backdrop, we document several deviations. The most striking feature is the dearth of parasitoids, which appears to affect all parasitoid groups and host communities, in line with the resource fragmentation hypothesis (*13*). The presumably tropical source faunas appear similarly devoid of parasitoids, but post-colonization processes have lowered the proportion further, potentially due to the extreme microendemism of Madagascan hosts. The nasty-host hypothesis (*14*) predicts larger effects on parasitoids of toxic hosts but there is scant support in the literature for the supposed variation in toxicity, and it matches parasitoid diversity poorly (see Supplementary Text for details) as observed also in other analyses (*11*). Another notable difference compared to temperate faunas is the richness of beetles among the non-parasitoid insects. This appears to be a characteristic retained from the source faunas and could be associated with features shared among tropical forests.

Single Malaise trap catches are known to produce biased samples of insect diversity (*42*, *43*). Nevertheless, the validation tests (see Materials and Methods for details) suggest that our methodology generates compositional estimates that are largely representative of the true fauna, with a few notable exceptions such as beetles being underrepresented. The accuracy appears to be due more to the depth of temporal and spatial sampling than to the addition of leaf litter samples. A potential concern is the lack of samples from the canopy fauna, the composition of which is poorly known (*44–46*). The validation tests were based on temperate data for which the canopy fauna is small in comparison. However, we note that metabarcoding detects larvae as well as trace DNA of insect hosts and prey, potentially picking up many canopy species indirectly and correcting for other sampling biases observed when only adult specimens are identified.

Our study clearly necessitates a reassessment of the assumptions underlying many current estimates of global insect diversity. It is obvious that a better understanding of the diversity of Diptera and saprophages is more important than previously realized, and that arguments based on global host-parasitoid species ratios lack support. We refrain from further speculation here, but agree with recent analyses (*8*, *47*) that the consensus estimate of 5.5 million insect species (*7*) needs to be revised upwards.

The patterns that emerge from this and other recent studies represent the outcome of processes that have shaped the structure of insect communities over millions of years. The fact that the patterns appear to be global and predictable suggests that insect communities have a substantial degree of resilience at this temporal scale and compositional granularity. It remains to be seen to what extent this translates to resilience at the shorter time scales and in the detailed structure of insect communities relevant in the context of current environmental changes.

The forests where we sampled represent increasingly isolated islands of habitats that originally covered much of Madagascar. Even though we now have a better understanding of how the insect communities in these islands are structured at a coarse level, it is an urgent priority to extend these types of studies to more fine-grained analyses (*48*) and to the remaining parts of the country. The goal must be to understand how the insect communities and the ecosystem services they deliver are affected by current stressors, and to mitigate and possibly reverse the current rapid biodiversity loss. We now have the means to do so and effectively monitor change over time, even though the spectacular diversity and long tail of rare species in these communities mean that we need to approach conservation decisions in new ways (*49*).

## Acknowledgments

Computational resources were provided by the National Academic Infrastructure for Supercomputing in Sweden (NAISS), funded by the Swedish Research Council.

## Funding

Knut & Alice Wallenberg Foundation grants 2017.0003 and 2020.0239

SciLifeLab & Wallenberg National Program for Data-Driven Life Science

National Bioinformatics Infrastructure Sweden (NBIS) at SciLifeLab

Swedish Research Council grant 2021-04830 (FR)

Swedish Research Council grant 2021-05563 (AFA)

Swedish Research Council grant 2021-03784 (AJMT)

Polish National Science Centre grant 2018/31/B/NZ8/01158 (PŁ)

Mistra, the Swedish Foundation for Strategic Environmental Research grant DIA 2020/10 (EG)

Sistema Nacional de Investigación, SENACYT, Panamá (GPAL)

National Natural Science Foundation of China, research fund for international young scientists grant 32350410418 (NNG)

Brazilian National Council for Scientific and Technological Development grant 309114/2021-7 (BCB)

## Author contributions

Conceptualization: FR, AFA, AJMT, TR, JB, BLF, PŁ, AM, EIE, RG, LJAD Methodology: FR, AFA, AJMT, TR, BLF, JB, PŁ, AM, EIE, RG, LJAD, DR, ETR, JS, CCW, EG Investigation: FR, NNG, BCB, GB, SK, FAL, JH, TH, GPAL, JS Visualization: FR, RG, EG Funding acquisition: FR, AFA, AJMT, TR, BLF, AM Project administration: AM, EIE Writing – original draft: FR Writing – review & editing: All authors

## Data, code, and materials availability

All source data and scripts used for the analyses and for generating the plots are available at https://github.com/insect-biome-atlas/paper-mg-insects (v 1.0) and in an archived version at Zenodo (DOI: 10.5281/zenodo.23262459); an overview of how the repository is organized and how it can be used is provided in the Supplementary Text. The processed metabarcoding data (version 4) and sample metadata (version 6) used here and how to access them are described in (*24*).

## Supplementary Materials

## Materials and Methods

### Field Sampling

Field sampling was part of the Insect Biome Atlas (IBA) project (https://insectbiomeatlas.org) and is described in detail elsewhere (*24*). In brief, we deployed 50 Malaise traps across 33 forest sites (parks) in Madagascar: 25 Malaise traps in rainforests (14 sites), 2 Malaise traps in montane rainforests (2 sites) and 23 Malaise traps in dry forests (17 sites) (fig. S1A). At 8 of the 33 sites, we deployed 3 to 4 traps within a single habitat. Distances between Malaise traps at these multitrap sites ranged from 100 to 4700 meters. At the remaining sites, a single trap was used. Malaise trap samples were collected weekly from August 2019 to July 2020, 12 months in total. Every fourth sample from each trap location was left at the Madagascar Biodiversity Center (https://www.madagascarbio.org/) to help build a natural history collection of Malagasy insects; our analysis is based on the remaining 1,919 samples.

At each Malaise trap location, four leaf litter samples were collected, one in each direction of the Malaise trap, five meters away from it. Each sample was concentrated to 2 L of leaf litter from a 1 m^2^ square using a Winkler sifter. Before sifting, the leaf litter was minced with a machete to dislodge any insects hiding in the twigs or decayed logs. Within 12 hours of sampling, living arthropods were extracted at ambient temperature overnight into a 100 mL bottle filled with 95% ethanol using a mini-Winkler extractor for a total of 12 hours. Samples were collected once per trap site in the period from January to November 2020.

### Metabarcoding

DNA extraction and sequencing of the Malaise trap and leaf litter samples is described in detail elsewhere (*24*). In brief, Malaise trap samples were processed using the FAVIS mild lysis protocol (*50*), targeting an approximately 418 bp long piece of the CO1 gene matching the tail end of the standard CO1 barcode. Five biological spike-ins were added to each sample for calibration, to obtain more reliable abundance data (*51*). Leaf litter samples were processed with the same protocol, except that biological spike-ins were not added.

After sequencing on an Illumina NovaSeq 6000 instrument, reads were denoised in the nf-core/ampliseq Nextflow workflow v2.4.0 (*52*), which uses the DADA2 algorithm (*53*) to infer amplicon sequence variants (ASVs). The original ASVs (*24*) were then reprocessed with the HAPP pipeline (*54*). In HAPP, we chose OTU clustering with swarm, chimera filtering using the samplewise algorithm, and noise filtering using the NEEAT algorithm as in (*24*). However, taxonomic assignments with the *k*-mer based SINTAX method (*55*), as implemented in vsearch v2.29.1 (*56*), and the alignment-based VSEARCH method implemented in QIIME2 v2024.10 (*57*) were recomputed based on version 5 of the BOLD CO1 database described in (*54*), updated in February 2026, and using the original BOLD taxonomic annotations of the reference sequences rather than the GBIF-processed annotations as in (*24*). As the taxonomic annotations are used in parallelizing some HAPP steps, we reran the entire pipeline, which caused a few subtle changes in the generated OTU data. The new data are available as version 4 of the processed ASV data from the IBA project (*58*). The taxonomic assignment of ASVs for downstream analyses was based on phylogenetic placement on a custom reference tree (see below) using the dyn-heur algorithm in EPA-NG v0.3.8 (*59*) through GAPPA v0.9.0 (*60*).

### Taxonomic annotation strategy

Placing the mostly undescribed Madagascan OTUs in the correct family is challenging due to the gaps in available barcode reference libraries. Standard methods, such as *k*-mer matching or alignment to reference sequences, struggle to place OTUs correctly in higher taxa unless there are close relatives in the library (*54*), which is unlikely to be the case for the majority of Madagascan OTUs. In such cases, phylogenetic placement in a reference tree is a more reliable method (*54*, *61*), and this is the approach we took here.

### Reference tree

To obtain a suitable reference tree for taxonomic annotation of Madagascan hexapods, we started with the Chesters tree (*62*), comprising approximately 50,000 insect species, with the quality improvements, expanded Entognatha representation, and extended set of outgroups introduced in (*54*).

We then applied additional improvements as follows. First, we removed erroneous sequences. We ran a BLAST search of all sequences in the tree against the core_nt database, and manually checked all sequences with close hits to sequences in other orders than the one to which they were supposed to belong. This identified around 40 sequences that represented various types of errors (human sequences, bacterial symbiont sequences, and sequences of other insects than the taxa the entries were supposed to represent). We also removed a Diptera sequence (*Lilaea fuliginosa*), which was incorrectly placed in the Orthoptera part of the Chesters tree. Second, we made sure that the tree followed the current understanding of order-level relationships by extracting all order-level clades and reinserting them on the dated order-level backbone from the phylogenomic analysis in (*63*). Third, we expanded the coverage of species-poor and phylogenetically isolated order lineages—Zoraptera, Embioptera, Phasmatodea and Notoptera (Mantophasmatodea + Grylloblattodea)—to reduce the risk that these long branches inappropriately attracted random authentic hexapod clusters or noise clusters. We also expanded and improved the coverage of the Entognatha + outgroups, and four taxa where the original tree would have been problematic in the ecological composition analysis: Cecidomyiidae, Braconidae, Ichneumonidae and Chalcidoidea.

#### Entognatha and outgroups

This part of the tree was added in (*54*), largely based on (*64*). During the error checking for this analysis, we discovered that the sequence attributed to *Fujientomon* (Protura) is likely a yeast sequence; it was replaced with the CO1 sequence from the mitochondrial genome of *Sinentomon erythranum*, which belongs to the same order as *Fujientomon*. We also discovered that the *Neocondeellum brachytarsum* sequence is likely to be a human sequence. The other *Neocondeellum* sequence does not have any close BLAST hits in GenBank beyond itself, so we refrained from using this sequence and simply deleted *Necondeellum* from the analysis. A sequence of *Wolbachia* (a common insect symbiont in the alphaproteobacteria, from which mitochondria originated) was added to help in the capture of bacterial contaminants. Finally, a plant sequence was added (from the mitochondrial genome of *Oryza*), to improve the detection of DNA from grass pollen, which is often found in Malaise trap samples. For the phylogenetic analysis, we generated partial constraints for the clades in (*64*) and hard constraints for families and for well-established relationships among outgroup taxa, as in (*54*).

#### Isolated order lineages

We expanded taxon coverage with additional CO1 sequences retrieved from GenBank for the isolated order lineages. In the phylogenetic analyses of these taxa, we used hard constraints for the orders and for the higher clades—Embioptera + Phasmatodea and Notoptera (Mantophasmatodea + Grylloblattodea)—where relevant, with lower-level relationships resolved by the CO1 data.

#### Cecidomyiidae

The expansion was based on the multigene analysis in (*65*), from which we extracted the CO1 sequences. We then added CO1 sequences from GenBank for some taxa that were still poorly represented, and the sequences in the original tree. In the phylogenetic analysis, we constrained all clades with posterior probability > 95% in the Bayesian analysis of (*65*) (their fig. 1C) using partial constraints, and added hard constraints for the families included in the analysis, leaving the three taxa that were treated as *incertae sedis* in (*65*) unconstrained.

#### Braconidae

We based the improvement on the analysis of ultraconserved elements (UCEs) of (*66*) (their figs 3-8). First, to the braconid sequences in the Chesters tree, we added the available CO1 sequences of the missing subfamilies, and of the genera included in the UCE analysis but not present in the Chesters tree. The subfamily classification of the braconid sequences in the Chesters tree was then retrieved from the NCBI taxonomy and updated using information from the UCE analysis. For a few Doryctinae in the Chesters tree, placement into the Doryctinae (s. str.) or Doryctinae (South America) clades of the UCE analysis was inferred from a preliminary phylogenetic analysis using MrBayes, where they were allowed to float in the UCE tree together with the other non-UCE taxa to find their affinities based on the CO1 data. Finally, we manually retrieved full CO1 sequence data from mitochondrial genomes of two ichneumonid representatives suitable as outgroups. For the phylogenetic analysis, we used all higher clades (subfamily or above) with more than 95% bootstrap support in the UCE analysis as partial constraints, and the families as hard constraints.

#### Chalcidoidea

For Chalcidoidea, we started with the combined UCE and anchored hybrid enrichment (AHE) tree in (*67*), including the outgroup Mymarommatidae. We first collected available CO1 sequences from GenBank matching the tips in this analysis. We then updated the taxonomic placement of the original chalcidoid sequences in the Chesters tree, including sequences of the Mymarommatidae, to reflect the revised family-level classification in (*67*). Phylogenetic analyses were then run with the clades in ChP1 represented as partial constraints, and with the ingroup-outgroup separation and the family-level clades as hard constraints.

#### Phylogenetic analyses and final tree assembly

After expanding the taxon coverage for the isolated order lineages and the critical clades as described above, we converted the new set of CO1 sequences to amino acid sequences, aligned the amino acid sequence with MAFFT v7.526 (*68*), and converted the resulting alignment back to a DNA sequence alignment using pal2nal v7.526 (*69*). Sites with 90% or more gaps were then trimmed away, and topological constraints formulated where relevant. We then ran MrBayes v3.2.7a (*70*) analyses under a strict clock model with a codon-partitioned substitution model, including a discrete gamma model of rate variation across sites and sampling of the substitution model in the GTR space (‘nst = mixed’). We ran the analyses for 10 to 100 M generations using 16 runs of 4 chains each. We aimed to reach an average standard deviation of split frequencies (ASDSF) below 0.05, which we obtained for all datasets except Braconidae (ASDSF 0.06). The variance between runs was almost entirely restricted to more fine-grained taxonomic levels than the family-subfamily-clade level targeted in the ecological analysis. We picked one arbitrary, fully resolved tree from the sample of the posterior (the last tree in the first run), and used it to replace the corresponding clade in the original tree. The final, dated reference tree includes 48,313 insect species, including representatives of 854 insect families.

#### Phylogenetic placement

Before annotation using GAPPA, the taxonomy of the reference tree was made to conform to the NCBI taxonomy, and augmented with a “clade” level. The latter corresponds to subfamily in Cecidomyiidae, Braconidae and Ichneumonidae, to well supported clades at approximately the family level in Chalcidoidea (*67*), and to NCBI families in all other groups. The clade level is consistently referred to as “family clades” in the main text. When we use “family”, we refer to actual families in the NCBI taxonomy, that is, we treat Cecidomyiidae, Ichneumonidae and Braconidae as single groups. In the Chalcidoidea, where family classification has recently been revised, we use the family clades recognized here in both cases.

For phylogenetic placement, we first reoptimized branch lengths and substitution model parameters, and then placed sequences on the reference tree using the same model and parameters with the dyn-heur algorithm in EPA-NG. We only retained placements with accumulated likelihood weight ratio (aLWR) > 0.5. To compensate for the poor representation of outgroups in the reference tree, we used additional filtering based on SINTAX and VSEARCH matches to the BOLD reference library to remove OTUs that likely represent noise. Specifically, if SINTAX placed the OTU in a BOLD BIN, and the classification was different from that of GAPPA at the order level, or VSEARCH placed the OTU in a phylum other than Arthropoda, the OTU was removed. Before the compositional analyses, we removed all OTUs not placed in a family clade.

### Ecological composition

For the ecological composition analysis, we followed (*37*) and focused on two traits that are conservative enough that they can be reasonably assumed, in most cases, to be conserved at the family clade level: the main feeding niche and main feeding microhabitat. This is the niche and microhabitat occupied by the immature stages (the main feeding stages), and may or may not be the same as the niche and microhabitat of the typically more short-lived adult stages.

Following (*37*), main feeding niches are defined as follows:

- Parasite. This includes bloodsuckers, endoparasites (botflies) and exoparasites (lice).
- Phytophage (plant feeder). This includes both chewers and sap suckers, as well as stem borers, leaf miners, root feeders and gall inducers.
- Phytophage-parasitoid. This is restricted to all primary parasitoids of plant feeders.
- Predator. This is restricted to free-living predators; it does not include parasitoids.
- Predator-parasitoid. This includes both primary parasitoids of predators and all hyperparasitoids (parasitoids of parasitoids).
- Saprophage (decomposer). This includes scavengers, decomposers, fungivores and microflora/bacterial grazers.
- Saprop*hage-parasitoid*. This is restricted to all primary parasitoids of decomposers.

The microhabitats of the main feeding stages (larvae, nymphs) are defined as follows:

- Fungi. In or on fruiting bodies of macrofungi.
- Homeothermic animals. On homeothermic animals (birds, mammals). The habitat of parasites (bloodsuckers, endoparasites and exoparasites).
- Plants. In or on aerial parts of living plants, excluding wood.
- Soil. In soil, including ground litter as well as living roots.
- Temporary habitats. In or on ephemeral resources of rich nutrients, such as bird nests, carrion, dung and fermenting sap. Also includes indoor habitats.
- Water. In or on water.
- W*ood*. In wood.

When there is known variation within the group, it is coded for the dominant trait, that is, the trait considered likely to characterize the majority of species. Parasites, most of which live on homeothermic animals, constitute a small fraction of the total hexapod fauna and are rarely detected in Malaise traps or litter samples. Therefore, we omitted them from the subsequent compositional analyses.

Data for Sweden were taken from (*37*), with minor adjustments to fit the NCBI taxonomy. Data for Madagascar were coded separately based on standard references complemented with primary literature, as noted in the provided source data files. The taxonomy and trait data were complemented with additional information for taxa with extended representation in the reference tree, as noted in the source data files.

### Madagascan radiations

To analyze Madagascan insect radiations, we focused on the 40,375 OTUs placed on a single branch in the tree with a Likelihood Weight Ratio of at least 0.5. To provide a conservative lower-bound on the number of radiations, we grouped the OTUs by the branch to which they attached, referring to such a group as an “endemic radiation” if it included at least five OTUs in our data. We did not attempt to resolve the phylogeny of each OTU group; instead, we assumed that it was monophyletic, and computed its age as the mean age of the attachment points of the individual OTUs to the reference tree.

Before analyzing the Madagascan radiations, we identified all species in the reference tree recorded from Madagascar in GBIF (*71*). If the species or clade in the reference tree to which an OTU attached consisted exclusively of Madagascan species, we used the stem attachment point of the Madagascan clade in the reference tree instead of the individual OTU attachment point.

We recognize that this analysis provides only a rough first approximation of the number and age of Madagascan radiations. As the coverage of appropriate outgroups in the reference tree increases, and the phylogeny of each OTU group is reliably resolved, the number of radiations is likely to increase and their estimated ages are likely to decrease in many cases.

### Diversification analyses

To test whether the composition of the fauna could be due simply to the set of insect lineages that successfully colonized Madagascar and their subsequent diversification, we developed a null model based on a joint time-dependent birth-death (TDBD) model with a constant turnover and an exponential decline over time in net diversification, capturing saturation effects. Specifically, the model assumes

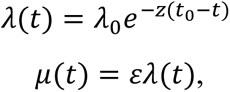

where *λ(t)* and *μ(t)* are the speciation and birth rates at time *t*, respectively, *λ_0_* is the speciation rate at *t_0_*, the time of colonization, *z* is the rate of decline in net diversification, and *ε* is the turnover rate. We conditioned the model on the age (*t_0_* values) and number of descendants of the 6,476 radiations we detected in Madagascar, defined as described above for Madagascan radiations but without any constraints on the number of OTUs. In the main analysis, we accommodated the sampling bias against Coleoptera documented in the validation tests (see below), estimating the bias using the ratio of Coleoptera to Lepidoptera OTUs in the Swedish IBA data (from all habitats) compared to the ratio of Coleoptera to Lepidoptera species in the known fauna (very few species remain to be discovered in either of these orders in Sweden). In an alternative analysis, we did not correct for any sampling bias. Since the sampling fraction is not an identifiable parameter in this model, we arbitrarily set the unbiased sampling probability to 1.0.

The posterior was estimated using MCMC with drift kernels in TreePPL v0.4 (*72*), associating *λ_0_* and *z* with Exponential(1) priors, and *ε* with a Uniform(0,1) prior. We ran four independent MCMC samplers and ensured that they converged to the same estimates of the posterior distribution after discarding the first 20% of samples as burn-in. We then simulated 10,000 datasets from the posterior, using the observed age of each lineage and recording the predicted number of descendants and their taxonomic assignment and life history traits. This simulated distribution was used as a null model to understand in what direction (if any) post-colonization diversification processes have pushed compositional features of the Madagascan insect fauna.

### Temperate reference data

For validation purposes and as a reference, we used data for the Swedish insect fauna, collected and processed using the same methods as for Madagascar (*24*). The two countries occupy a similar area, facilitating comparison. Briefly, 203 Malaise traps were deployed across Sweden for a full year (2019), or as much thereof as feasible, within six major habitats across 100 sites: 102 traps in forests, 33 in croplands, 26 in wetlands, 16 in grasslands, 14 in alpine, and 12 in urban areas. At 21 of the 100 sites, we deployed up to six traps in a hierarchical design.

Distances between Malaise traps at these multitrap sites ranged from 90 to 1200 meters, sampling two different habitats when possible (three Malaise traps per habitat). The traps were emptied weekly during the growing season and bi-weekly or monthly otherwise. Leaf litter arthropod communities were sampled by collecting a total volume of 1 L of leaf litter from five different sites around each Malaise trap, 2.5 meters away from the nearest extremity of the trap. Within 48 hours after collection, arthropods were extracted from the samples using a Berlese funnel for 96 hours. Litter was sampled once per site during the growing season. Metabarcoding proceeded as for the Madagascan samples, except that six rather than five different biological spike-in species were used for calibration.

When we present reference values for temperate forests, they are based on analyses of only the Swedish forest data, spanning the nemoral to boreal transition of the northern temperate region. Validation tests (see below) used the Swedish data from all habitats. In both cases, we restricted the analysis to data generated after mild lysis of the Swedish Malaise trap samples. One fourth of the Swedish Malaise trap samples was metabarcoded again after homogenization to yield complementary information, but homogenate data were not available for Madagascar and these data were not used in the present study.

### Species richness and composition analyses

For analyses of the species pool, we used the R package vegan version 2.7-3 (*73*). For the spatial turnover analyses, we partitioned beta diversity into turnover and nestedness using the betapart package version 1.6.1 (*74*), and focused on the turnover component. Singletons were excluded from these analyses; if included, they shift Jaccard dissimilarity upwards without changing overall patterns. We used the R packages mgcv version 1.9.4 (*75*) to fit monotonic generalized additive models to turnover plots.

### Mapping the Madagascan insect fauna

We used sf version 1.1.1 (*76*, *77*) to generate maps of Madagascar with spatial features.

### Validation analyses

To validate our methodology, we analyzed the Swedish data from all habitats and compared the results to what is known about the Swedish insect fauna (*37*). Specifically, we tested the noise filtering and taxonomic annotation steps, and potential biases in the sampling methodology.

#### Noise filtering and taxonomic annotation

DNA metabarcoding data are noisy: nuclear sequences of mitochondrial origin (NUMTs), chimeras and other erroneous sequences may account for more than 70% of OTUs resulting from clustering of data from standard pipelines (*54*). This is a particular concern when analyzing poorly studied faunas, as one cannot rely on reference libraries of authentic barcode sequences in filtering out such noise, and there is little data from other studies against which to assess the quality of the results. Thus, one needs to largely rely on patterns within the dataset to accomplish this, as in the NEEAT algorithm used here. Similarly, taxonomic annotation is challenging for poorly studied faunas because of the gaps in the available reference libraries, justifying the use of phylogenetic placement in this step rather than matching to a reference library. Both steps could potentially introduce noise and biases in the data. However, when applied to the Swedish data for which the reference libraries are significantly more complete, the phylogenetic placement performs on par with methods that rely on matching to the BOLD reference library (fig. S14), suggesting that this is not a serious issue even though there are some taxa, mostly in minor orders, for which spurious OTUs remain a problem (figs S15, S16). When applied to Madagascan data, the pipeline has similar performance in both the noise filtering and annotation steps as when it is applied to Swedish data (fig. S17).

#### Sampling biases

Because the Swedish temperate reference fauna is well studied, the Swedish IBA data can be used to assess the extent to which our methodology is likely to yield reliable estimates of the composition of the true fauna. For most families in the Swedish fauna, the true number of species is essentially known. However, there are still some families for which this is not the case; they almost exclusively belong to the orders Diptera and Hymenoptera. To compensate for the existing knowledge gaps, we used a recent estimate of the number of species in Swedish insect families (*37*) as the basis for the assessment rather than the recorded fauna.

Comparing species accumulation curves for the Swedish IBA data with the published estimate reveals that the proportion of unknown species in the Swedish fauna must be higher for Diptera and Hymenoptera than previously assumed (fig. S18). A recent analysis showed that the IBA data include approximately half of the Swedish fauna of most lepidopteran families (*78*). A reasonable assumption is that the same holds true for Diptera and Hymenoptera: the bulk of the species diversity in both orders, like that in Lepidoptera, consist of small to medium-sized flying insects that are well represented in Malaise traps. We generated a revised estimate based on this assumption, keeping the relative percentage increase in different Diptera and Hymenoptera families the same as in the original estimate, but adjusting the total diversity of these two orders to twice that observed in the inventory. No other adjustments were made to the original estimate, as the remaining errors are likely to be small in comparison.

Using this revised estimate as ground truth suggests that our methodology largely gives a correct picture of the true fauna (figs S19–S21). The most notable discrepancies include an underrepresentation of Coleoptera and of predator parasitoids (fig. S19F). The extent of the sampling bias against Coleoptera can be estimated by comparing the ratio of Coleoptera to Lepidoptera species in the IBA sample to the ratio of known species in these orders, as the Swedish fauna of both is considered to be almost completely known (*37*). This suggests a sampling bias of 0.665 against Coleoptera, that is, a Coleoptera species is only 66.5% as likely to be sampled as a species belonging to other insect orders. The potential sampling bias against predator parasitoids is difficult to analyze further given the current state of knowledge.

The apparent underestimate of the proportion of species developing in temporary habitats (fig. S20E) is likely due in large part to an exaggerated estimate of the proportion of unknown scuttle flies (Phoridae) in (*37*). Molecular analyses indicate that the preliminary taxonomic work on scuttle flies on which the estimate was based is characterized by oversplitting of species (*79*, *80*). In addition, the few scuttle-fly life histories that are known are quite varied, making the inference of the dominant traits unusually challenging for this family.

The OTUs retrieved in the Swedish leaf litter samples largely overlap with those in the Malaise traps (fig. S22). Thus, their contribution to an accurate representation of the fauna is smaller than one might have expected. The OTU subset in the litter samples paints a slightly different picture of the fauna than the Malaise traps: it is richer in saprophages, predators, and beetles, while it is poorer in phytophages (figs S23–S25). However, in several cases the accumulation trends converge with those of the Malaise traps, suggesting that deeper sampling would reduce the discrepancy between the sampling protocols. In the Madagascan data, there is more complementarity between the litter samples and the Malaise traps (fig. S22), which could in part be due to the fact that the Madagascan data cover a smaller fraction of the complete fauna than the Swedish data. The differences in composition of the litter and Malaise trap samples are similar to those observed for the Swedish data (figs S26–S28).

### Visualization

Unless stated otherwise above, plots were generated with ggplot2 version 4.0.2 (*81*), patchwork version 1.3.2 (*82*) and cowplot version 1.2.0 (*83*)

## Supplementary Text

### Predictions of the nasty host hypothesis

According to the nasty host hypothesis, tropical plants are more toxic than temperate plants, resulting in phytophagous insects being less palatable in the tropics, which affects the richness of parasitoids with phytophagous hosts (*14*). Although not directly discussed in the original paper, the hypothesis implies that the effect on the diversity of parasitoids of predators or saprophagous hosts is indirect and small in comparison, which conflicts with our data as discussed in the main text.

A number of more detailed predictions have also been derived from the nasty host hypothesis (*14*). The hypothesis suggests that, among the parasitoids of phytophagous hosts, it is primarily the species that develop for a prolonged period of time in leaf-eating host larvae (so-called koinobionts) that are affected. This implies that parasitoids of eggs or pupae should be less affected, since these life stages supposedly are not as rich in toxic compounds. For similar reasons, the effect should be less severe on parasitoids of wood-living hosts and idiobionts—the parasitoids that consume their host rapidly.

In our data, we observe some patterns that agree with these predictions and others that do not. The overall ratio of Madagascan to Swedish forest species in our data is 3.0; parasitoid groups that are unaffected should have a similar ratio. We find that phytophage parasitoids among species living in wood are actually more diverse in Madagascar than in Sweden (ratio 4.6), probably partly explained by wood-living species being more prevalent in the Madagascan fauna but nevertheless clearly suggesting that this prediction of the nasty host hypothesis holds. We also observe patterns in line with the nasty host predictions for some egg parasitoids (Scelionidae, ratio 4.0) but less clearly for others (Mymaridae, ratio 2.3). The idiobiont pupal parasitoids in the ichneumonid subfamilies Cryptinae, Pimplinae (excluding the *Polysphincta* group) and Phygadeuontinae (including the *Chirotica* group) do seem to violate the nasty host predictions (ratios 0.3, 0.3, and 0.6, respectively), a result that is consistent with a recent analysis of global Malaise trap data (*11*).

### Accessing and Using Project Data and Scripts

The data and scripts used in the study are provided in an archived copy at Zenodo (<u>DOI:</u> <u>10.5281/zenodo.23262459</u>). There is a live version of the git repository at https://github.com/insect-biome-atlas/paper-mg-insects.git.

In a set of README files, the repository contains detailed instructions on how to download the IBA data and other big datasets used in the study from the public repositories where they reside, and how to regenerate the analyses and figures once this has been completed. Where feasible, we provide intermediate results and script outputs to facilitate the rerunning of small subsets of the analyses included in the paper, and to check outputs against those of the original analyses.

The source data for the expansion of the reference tree and for the trait coding are available in the ‘source_data’ directory in the repository in both legacy formats (Microsoft Excel and Microsoft Word) and open formats (semicolon-separated cvs and rtf formats). Key source data files and their contents are listed below (add suitable file format endings to find the corresponding files):

1. Life history trait data for Madagascan insects (MG_Hexapoda_Family_traits).
2. Life history trait data for Swedish insects (Ronquist_2020_SE_traits).
3. Entognatha taxonomy and traits data (Entognatha_taxonomy_traits).
4. Cecidomyiidae taxonomy and trait data (Cecidomyiidae_taxonomy_traits).
5. Ichneumonidae taxonomy data (Expanded_Ichneumonidae_taxonomy).
6. Ichneumonidae trait data (Ichneumonidae_traits).
7. Braconidae taxonomy data (Chesters_Braconidae_taxonomy_updated).
8. Braconidae trait data (Braconidae_traits).
9. Chalcidoidea taxonomy data (Chesters_Chalcidoidea_taxonomy_updated).
10. Chalcidoidea trait data (Chalcidoidea_life_history_traits).
11. References for taxonomy and trait data (Taxonomy_and_trait_references).

## Supplementary Figures

**Fig. S1.**
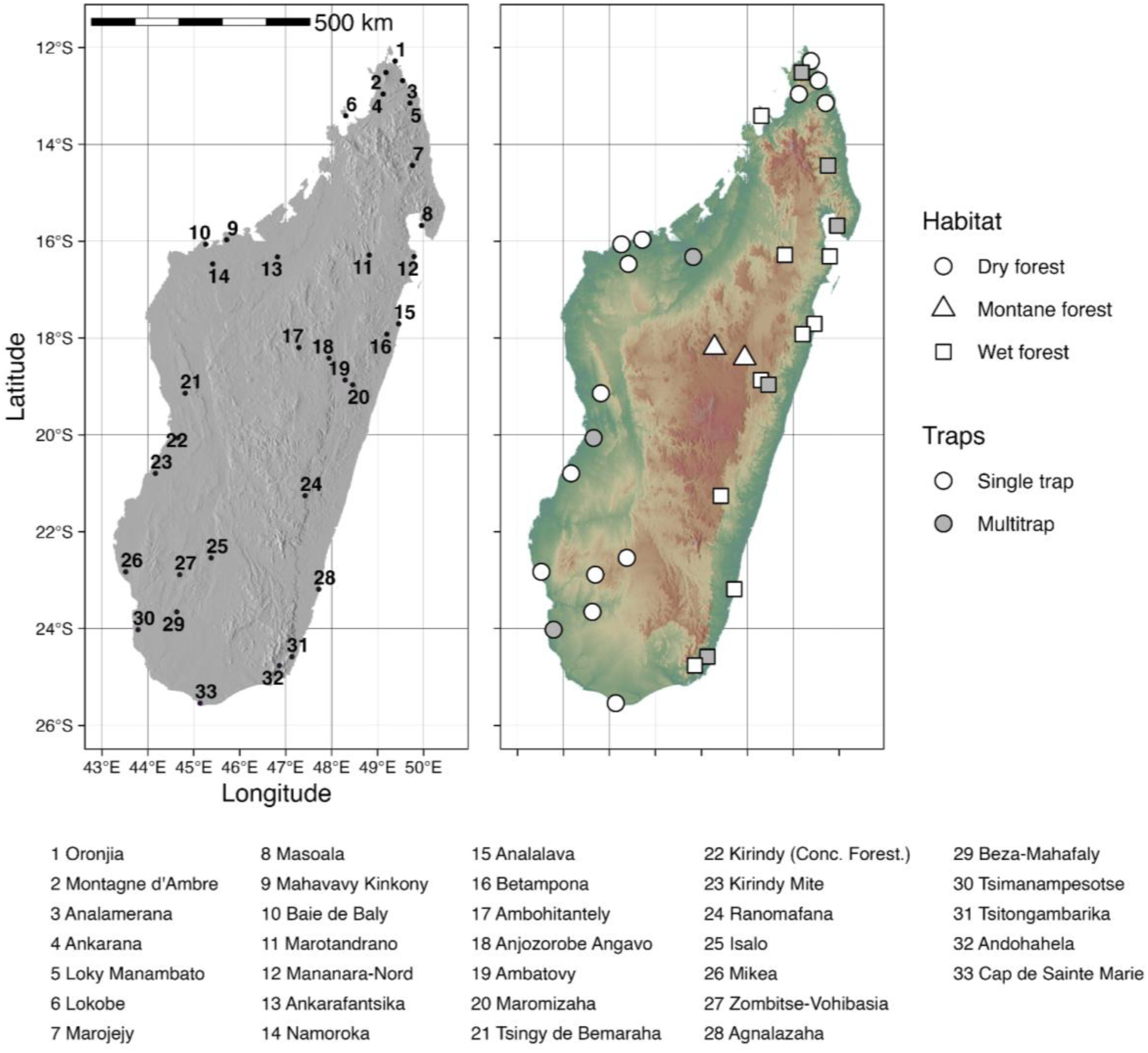
Sampling localities and forest types. Malaise-trap sampling localities across Madagascar. Left: sampling localities. Right: the same localities, classified by forest habitat (dry forest, montane forest or wet forest) with point fill indicating trap type (single trap or multitrap, the latter sampled with 3–4 Malaise traps within a locality). All localities were situated in forested habitats within protected areas, managed by Madagascar National Parks or by other conservation bodies (e.g. Missouri Botanical Garden, Fanamby, Asity, GERP). In the forest-type analyses, montane forest (“montane rainforest”) and wet forest (“rainforest”) localities were grouped together as “rainforest” and contrasted with dry forest.

**Fig. S2.**
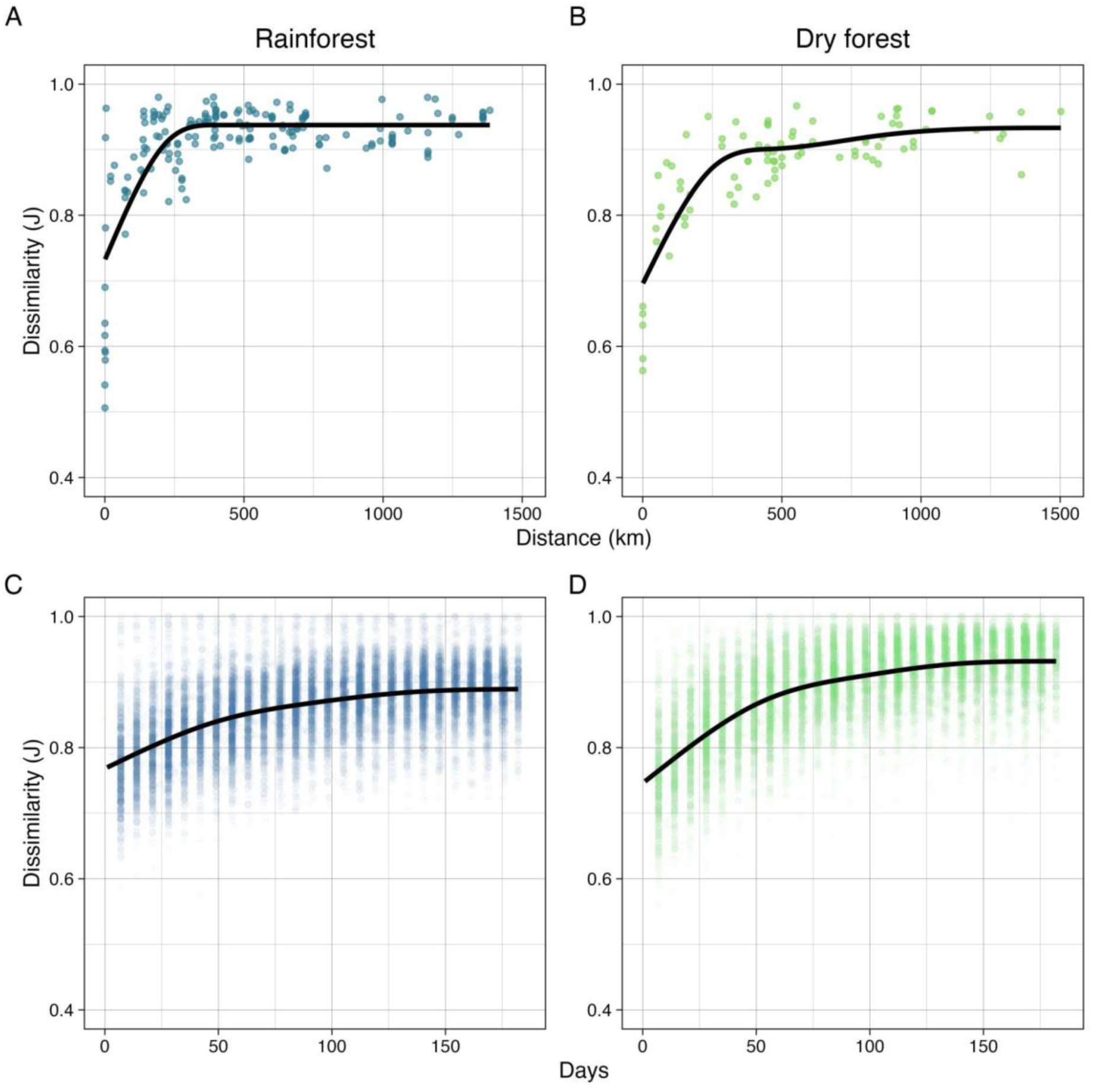
Spatial and temporal turnover by Madagascan forest type. Turnover is expressed as Jaccard dissimilarity between sample pairs separated in space or time. The fitted line is generated from a monotonic generative additive model with five knots. (**A–B**) Spatial turnover across rainforest and dry forest sites, respectively. (**C–D**) Temporal turnover within sites for rainforest and dry forest sites, respectively.

**Fig. S3.**
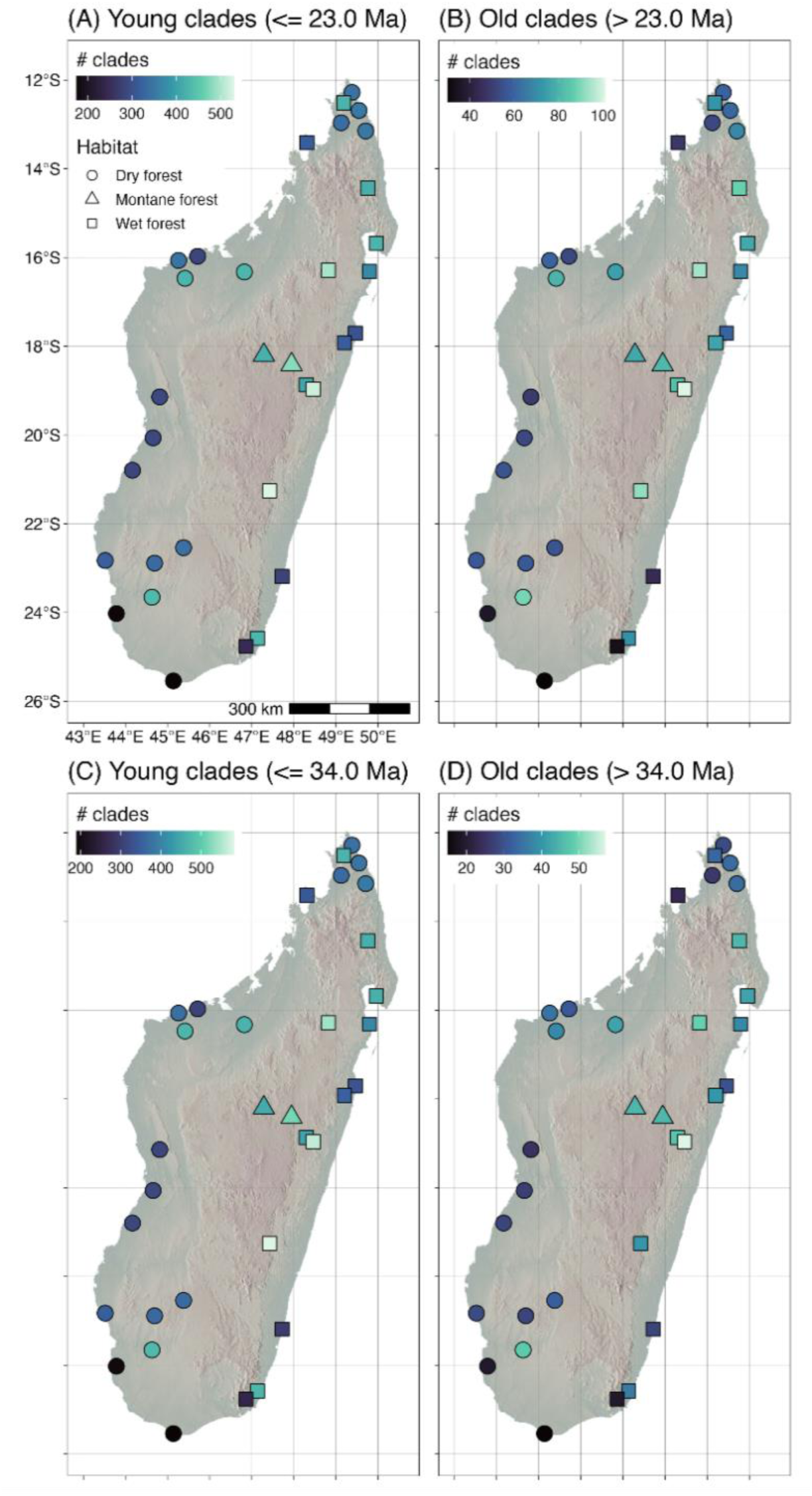
Distribution of old and young Madagascar lineages. The number of Madagascar radiations (≥ 5 sampled species) represented at each site, subdivided by age. (A) Radiations dating from the Miocene or later (age ≤ 23 Ma). (B) Radiations predating the end of the Oligocene (age > 23 Ma). (C) Radiations dating from the Oligocene or later (age ≤ 34 Ma). (D) Radiations predating the end of the Eocene (age > 34 Ma). For multitrap locations, we show the median value across traps.

**Fig. S4.**
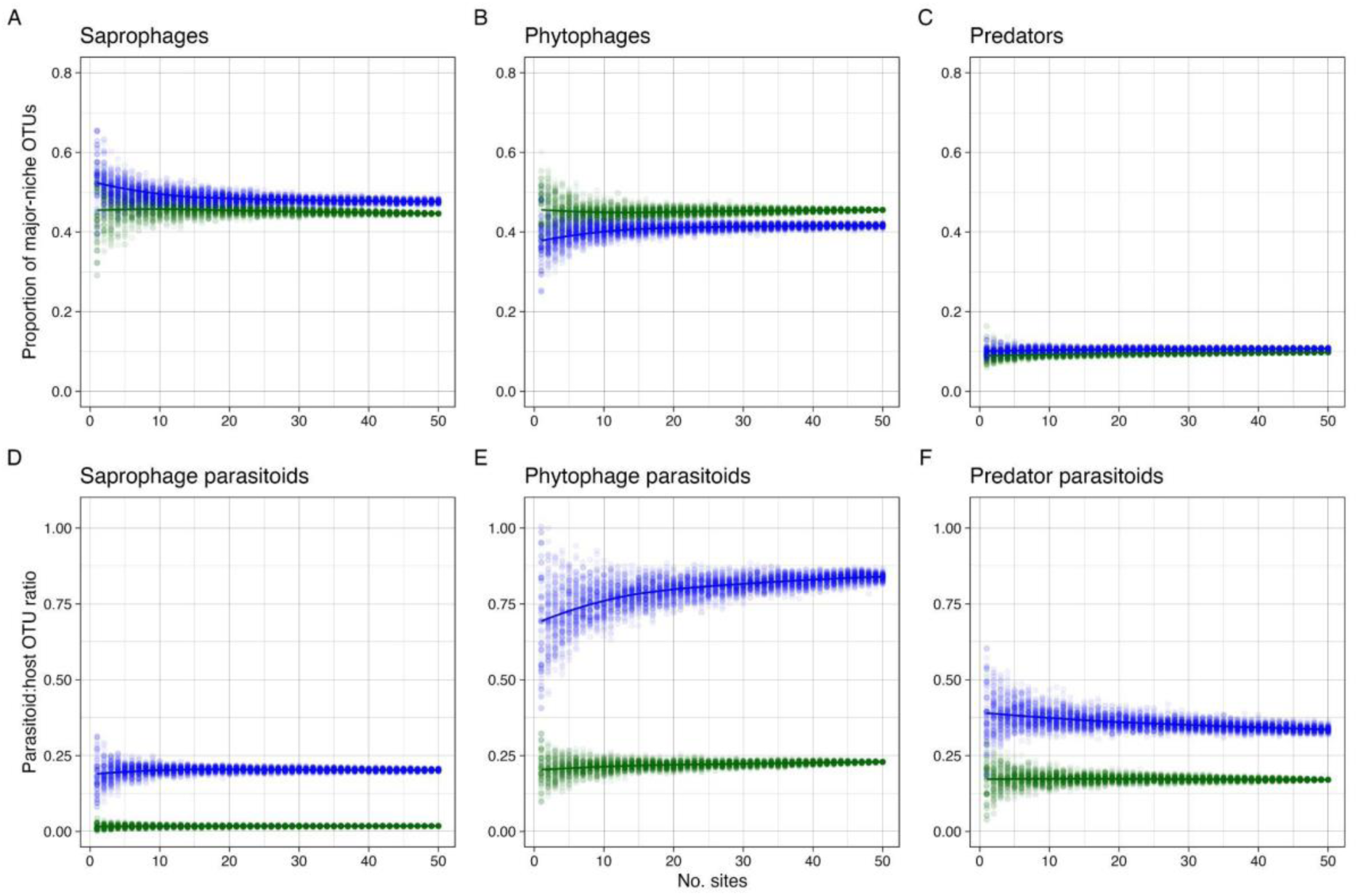
Spatial accumulation across sites of niche composition. The points represent 100 random samples without replacement for each number of sites. Madagascan data in green, temperate forest reference data in blue. (A–C) Proportion of OTUs belonging to the three major feeding niches. (A) Saprophages. (B) Phytophages. (C) Predators. (D–F) Parasitoid:host OTU ratios for saprophage parasitoids (D), phytophage parasitoids (E), and predator parasitoids (F).

**Fig. S5.**
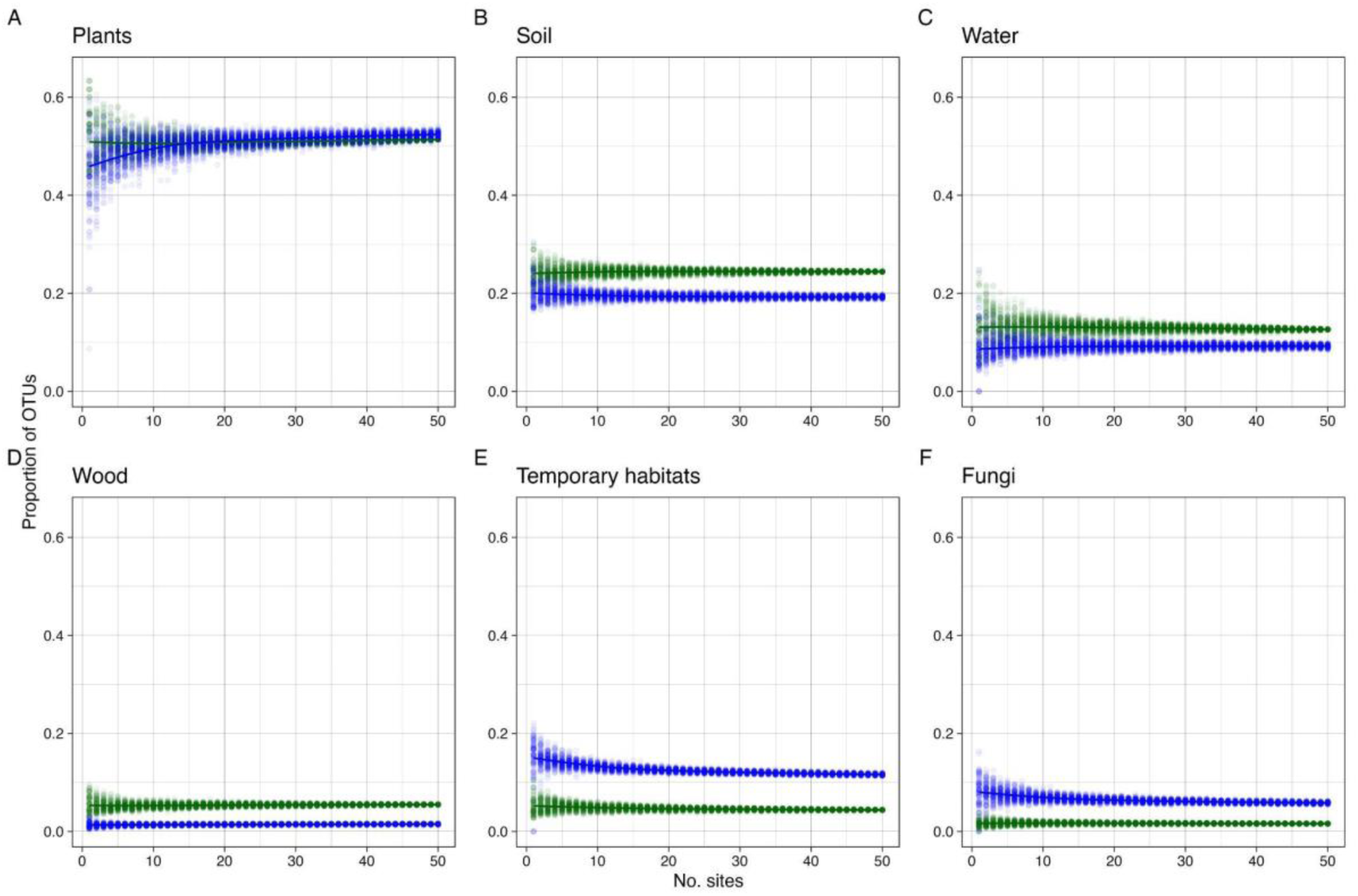
Spatial accumulation of microhabitat composition across sites. The points represent 100 random samples without replacement for each number of sites. Madagascan data in green, temperate forest reference data in blue. (A) Species developing in or on plants. (B) Species developing in soil, litter or on the ground. (C) Species developing in water. (D) Species developing in wood. (E) Species developing in temporary microhabitats, such as carrion or dung. (F) Species developing in or on fungi.

**Fig. S6.**
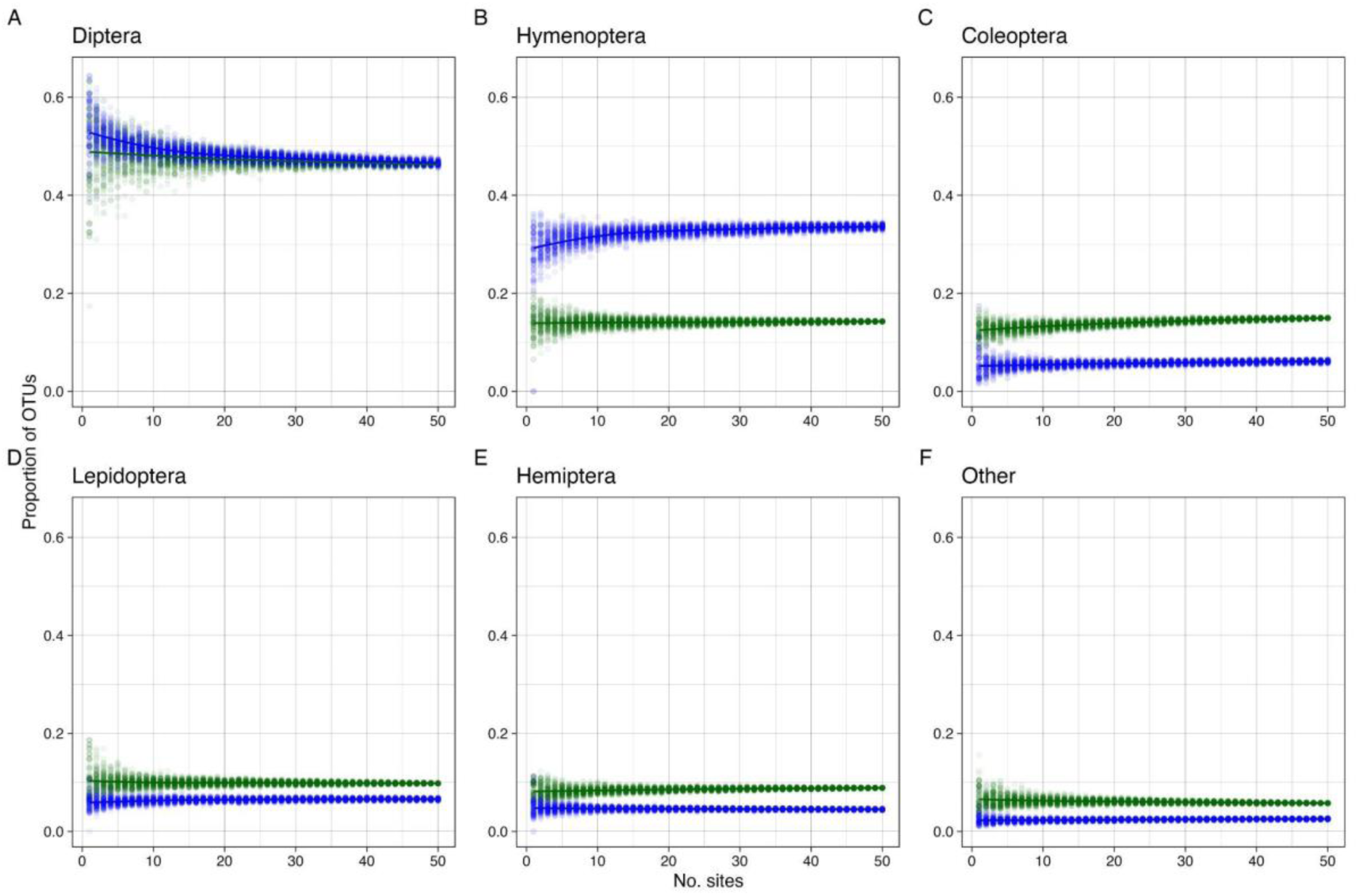
Spatial accumulation of taxonomic composition across sites. The points represent 100 random samples without replacement for each number of sites. Madagascan data in green, temperate forest reference data in blue. (A) Diptera. (B) Hymenoptera. (C) Coleoptera. (D) Lepidoptera. (E) Hemiptera. (F) Remaining hexapod orders. Note that the y-scale is different in plots D–F to display the differences among the less diverse taxonomic groups more clearly.

**Fig. S7.**
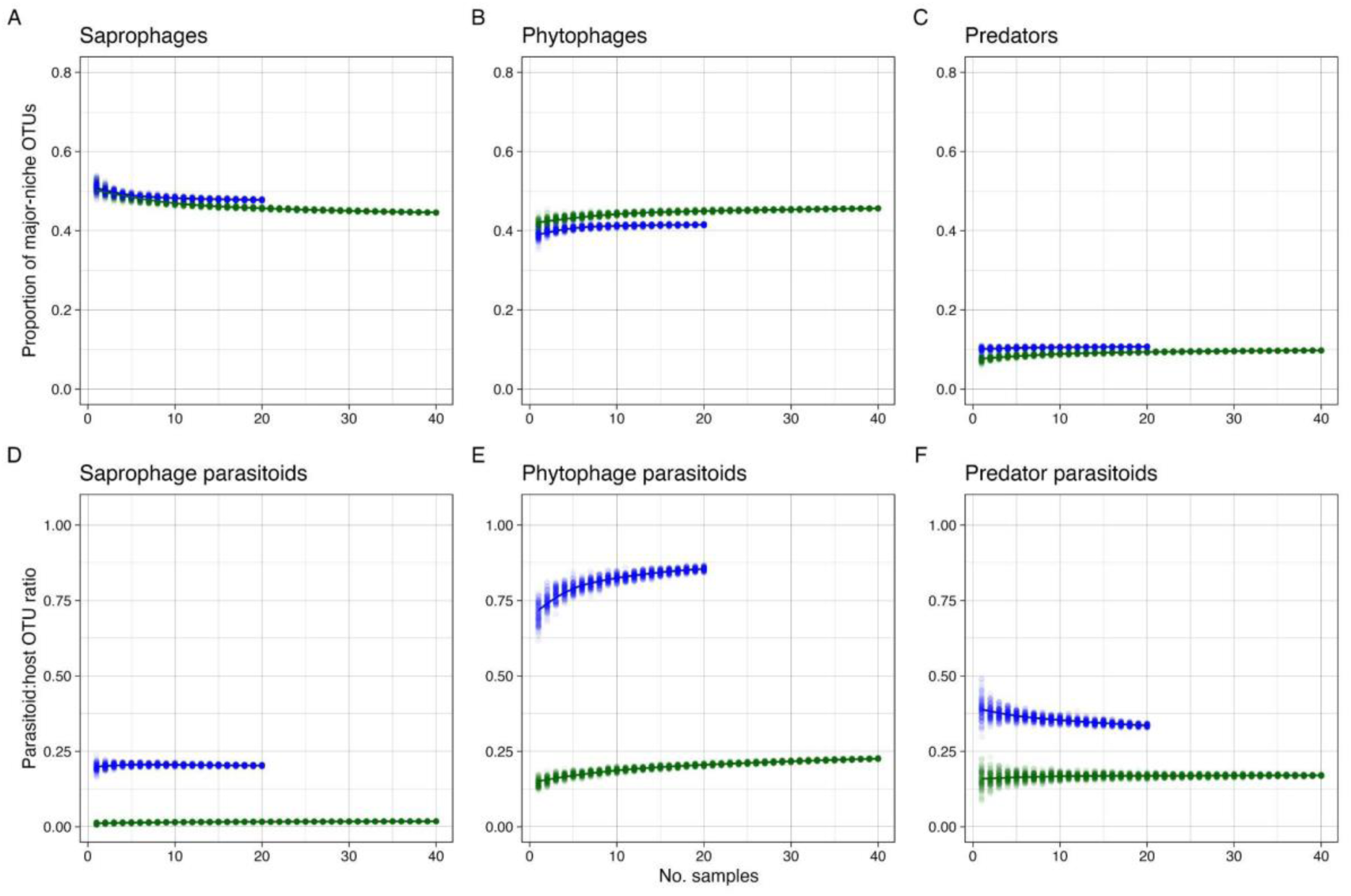
Temporal accumulation of niche composition across sites. The points represent 100 random samples without replacement for each number of temporal samples (weekly samples in Madagascar, weekly to monthly samples in Sweden depending on season) from the studied trap sites. Madagascan data in green, temperate forest reference data in blue. (A–C) Proportion of OTUs belonging to the three major feeding niches. (A) Saprophages. (B) Phytophages. (C) Predators. (D–F) Parasitoid:host OTU ratios for each of the three main communities. (D) Saprophage parasitoids. (E) Phytophage parasitoids. (F) Predator parasitoids.

**Fig. S8.**
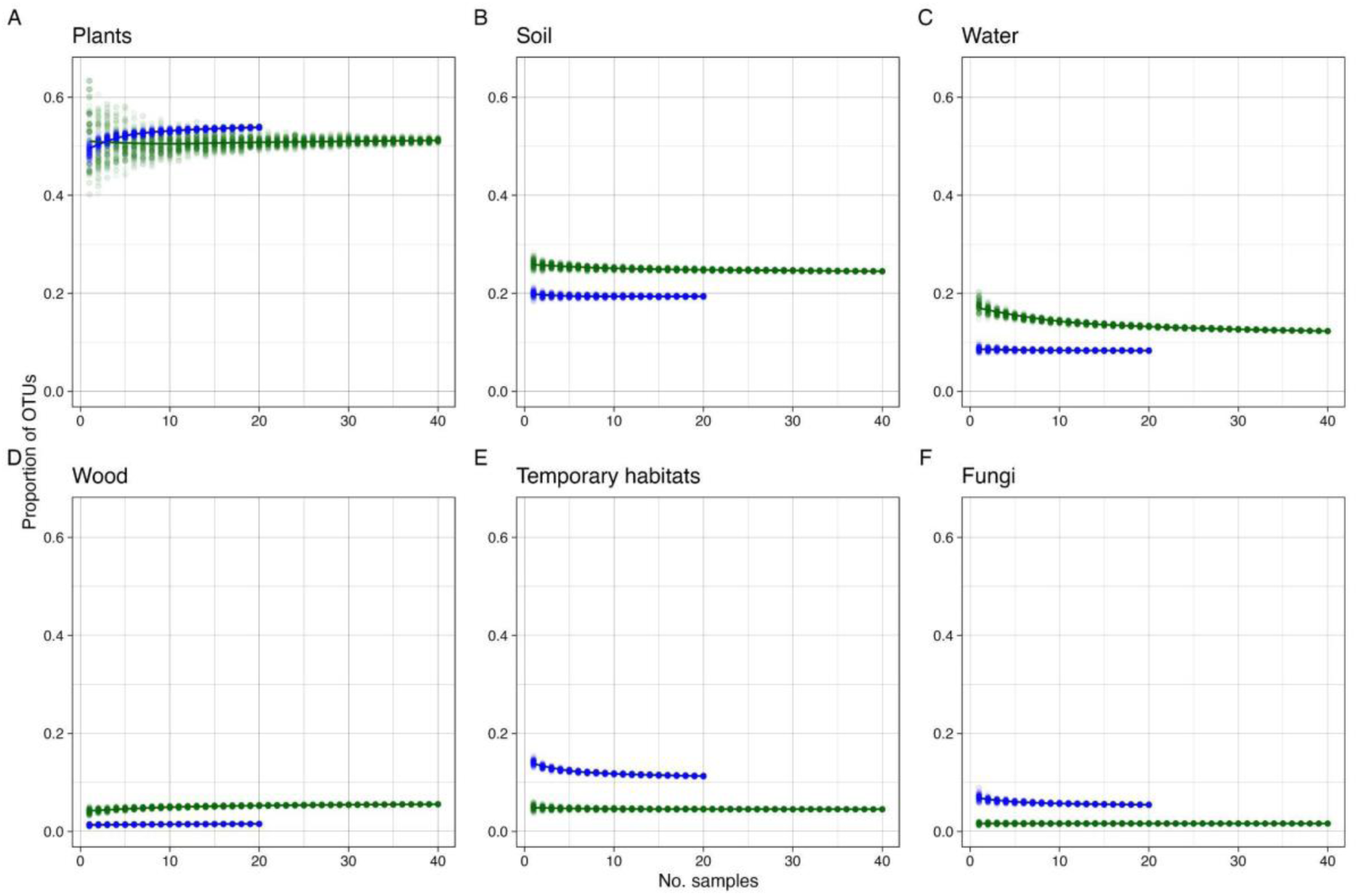
Temporal accumulation of microhabitat composition across sites. The points represent 100 random samples without replacement for each number of temporal samples (weekly samples in Madagascar, weekly to monthly samples in Sweden depending on season) from the studied trap sites. Madagascan data in green, temperate forest reference data in blue. (A) Species developing in or on plants. (B) Species developing in soil, litter or on the ground. (C) Species developing in water. (D) Species developing in wood. (E) Species developing in temporary microhabitats, such as carrion or dung. (F) Species developing in or on fungi.

**Fig. S9.**
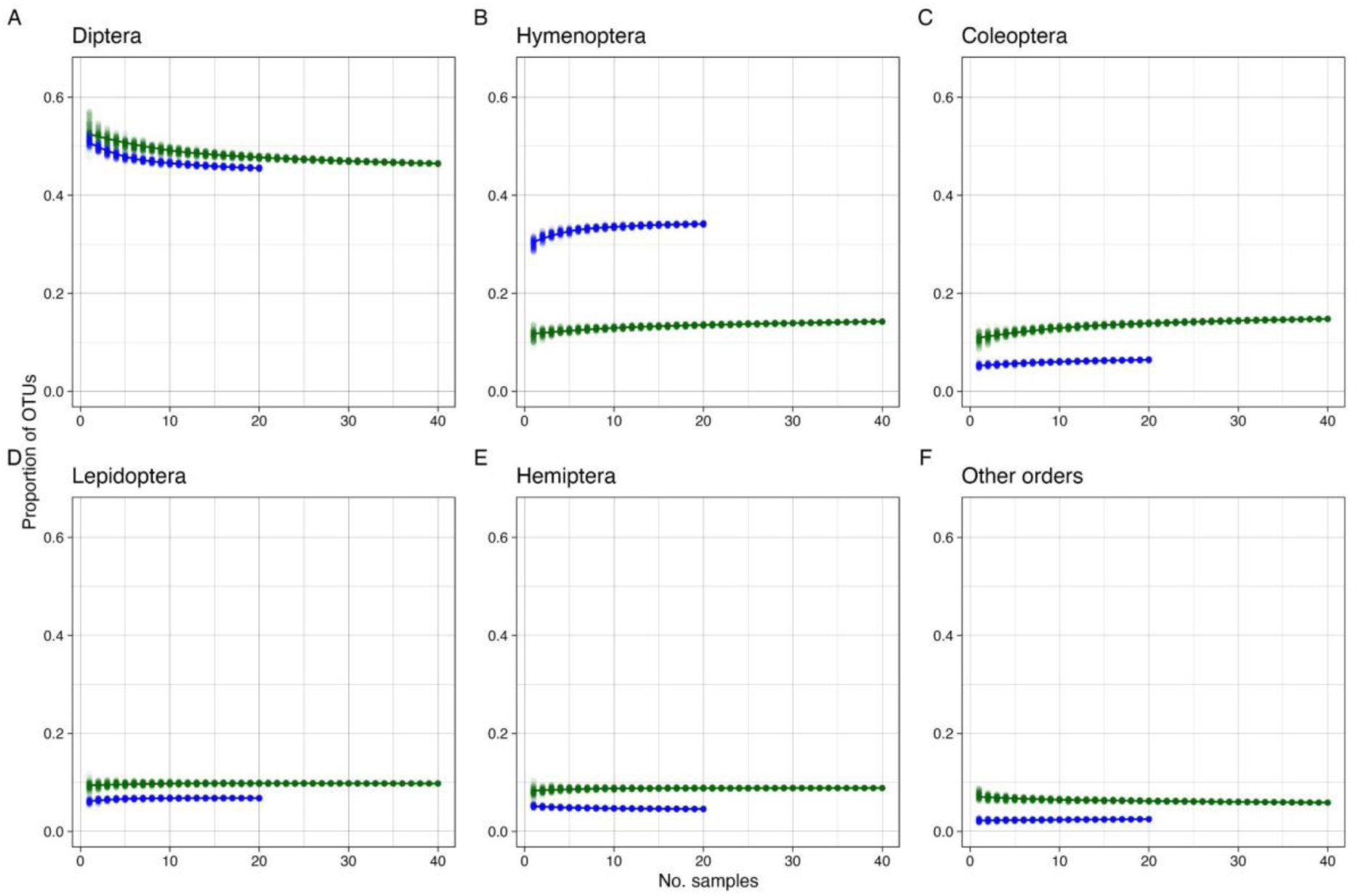
Temporal accumulation of taxonomic composition across sites. The points represent 100 random samples without replacement for each number of temporal samples (weekly samples in Madagascar, weekly to monthly samples in Sweden depending on season) from the studied trap sites. Madagascan data in green, temperate forest reference data in blue. (A) Diptera. (B) Hymenoptera. (C) Coleoptera. (D) Lepidoptera. (E) Hemiptera. (F) Remaining hexapod orders.

**Fig. S10.**
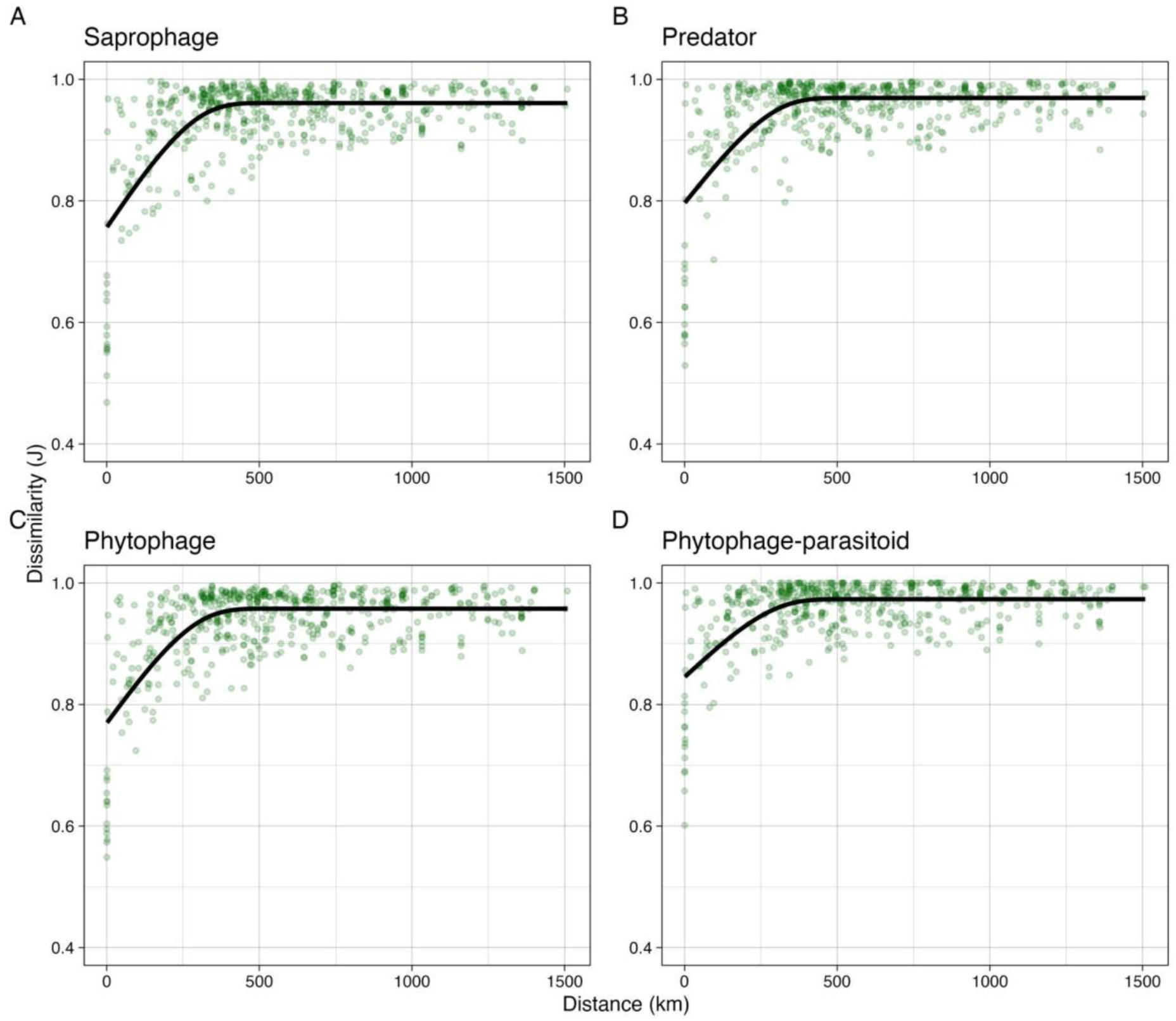
Spatial turnover of the three main feeding niches and of phytophage parasitoids in the Madagascan sample. Turnover is expressed as Jaccard dissimilarity between pairs of trap sites. The fitted line is generated from a monotonic generalized additive model with five knots. **Comment**. There are too few data points to meaningfully plot the spatial turnover for saprophage parasitoids and predator parasitoids. Note that the spatial turnover of phytophage parasitoids (**C**) is similar to that of their phytophage hosts (**D**). Compared to the temperate reference forest, the Madagascan forests are characterized by lower species diversity of parasitoids but similar parasitoid loads in the phytophage community. This suggests that Madagascan parasitoids have wider host ranges. An alternative explanation that we regard as less likely is that parasitoid loads are more unevenly distributed across host species in Madagascar. The wider host ranges in Madagascar could be achieved by parasitoids expanding their distribution ranges so that they can attack a larger set of closely related hosts; closely related host species are likely to vicariate with each other over a larger geographical range. It could also be achieved by parasitoids attacking a wider range of local hosts that are likely less closely related on average. The similarity in spatial turnover between hosts and parasitoids suggests that the latter is the more likely explanation.

**Fig. S11.**
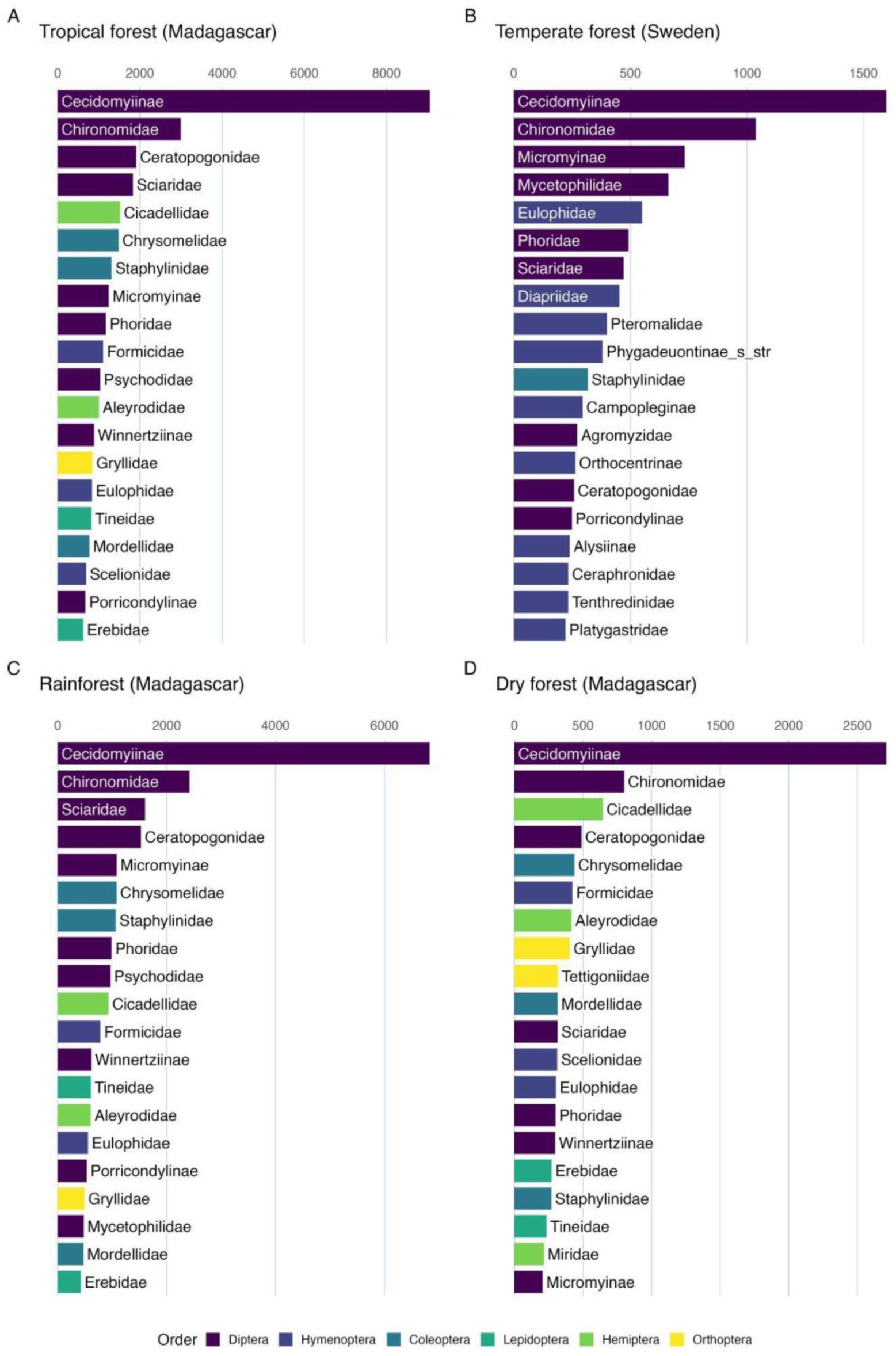
Most diverse family clades in Madagascan forests and the temperate reference forest. Top 20 family clades (corresponding to subfamilies in Cecidomyiidae, Ichneumonidae and Braconidae, and to families in other taxa) are displayed for each forest type, ranked by the number of OTUs placed in them.

**Fig. S12.**
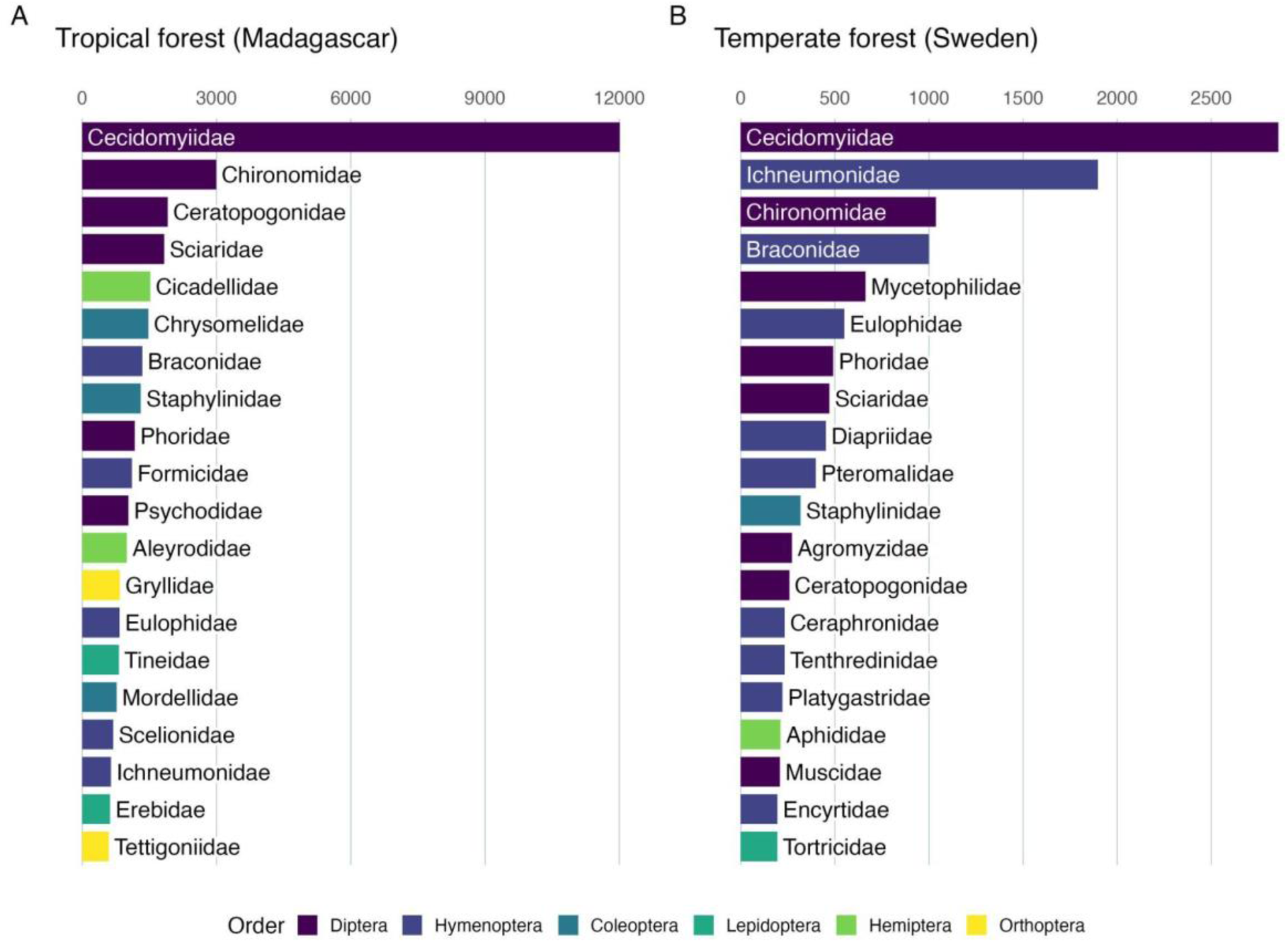
Most diverse families in the Madagascan forests and the temperate reference forest. The top 20 families are displayed for each dataset, ranked by the number of OTUs placed in them.

**Fig. S13.**
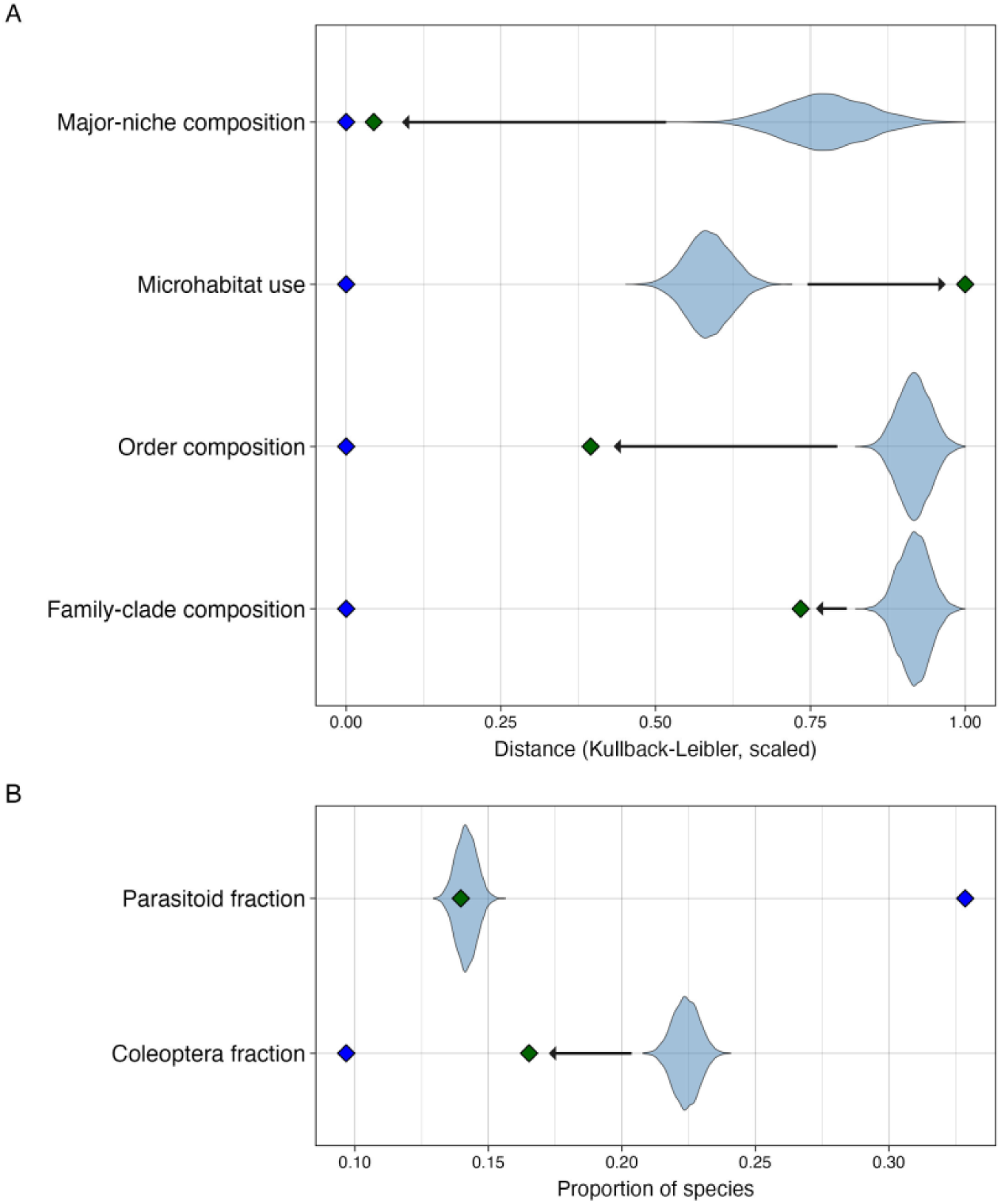
Diversification analysis of compositional features with no sampling bias correction. We fitted a common time-dependent diversification model to all Madagascan radiations, conditioning on the age and the number of sampled OTUs of each radiation. Unlike Fig. 5 in the main text, we did not correct for any sampling bias among insect groups here. By simulating data from the fitted model, we tested whether observed compositional features (green diamonds) deviated from the expectations under neutral diversification of colonizing lineages and, if so, whether the composition evolved (arrows) to become more similar or more different to that of temperate forests (blue diamonds). The simulations (violin plots) are likely to represent compositional features of source faunas filtered through partly stochastic and partly deterministic selection of successful colonizers. (**A**) Compositional features measured in terms of relative Kullback-Leibler distance to temperate forests. (**B**) Fraction of parasitoids and beetles (Coleoptera). **Comment**: Beetle lineages are diverse in terms of feeding niches and microhabitat use, explaining why the sampling bias against them has limited effect on most compositional features. However, the bias does affect the sampling of beetle species more than the sampling of beetle lineages colonizing Madagascar, changing the results of the analysis.

**Fig. S14.**
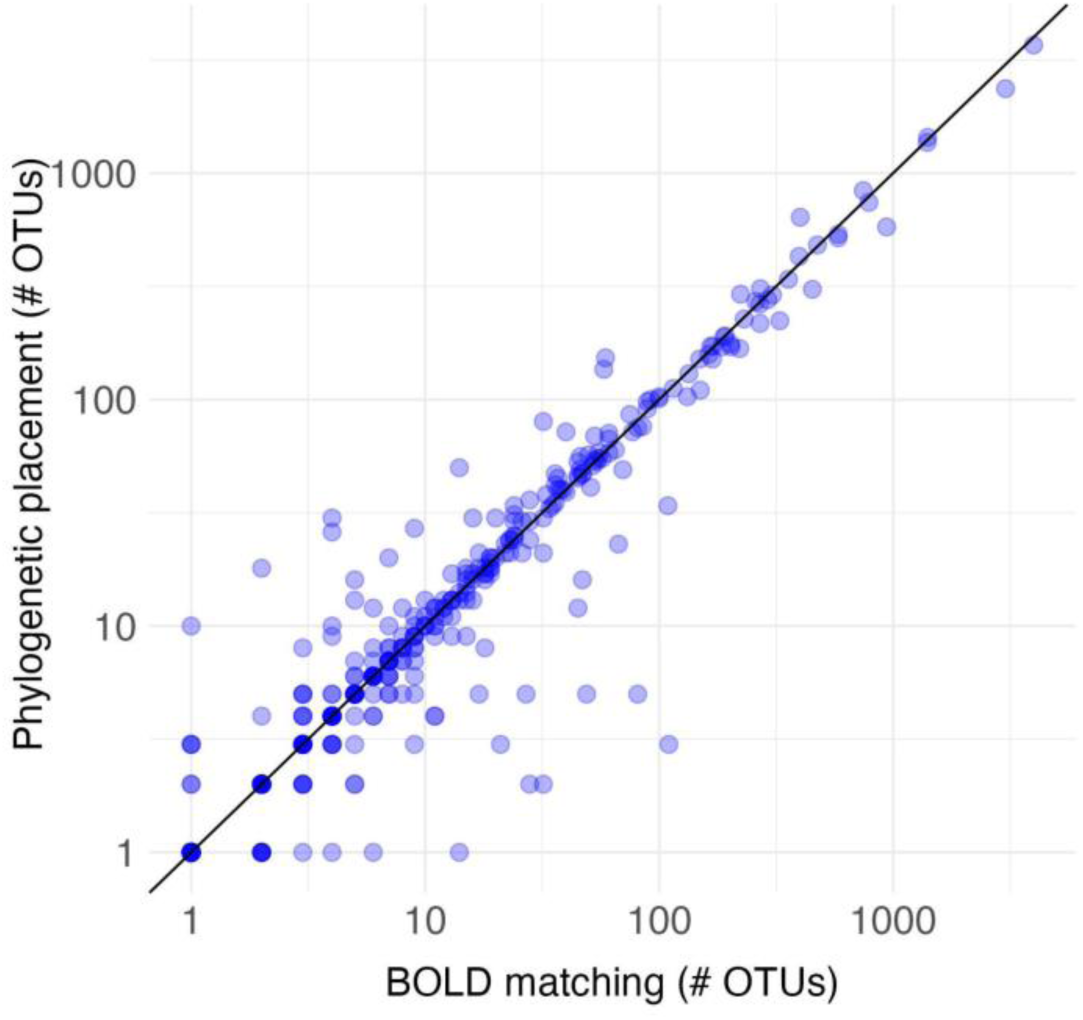
Validation of the phylogenetic placement methodology using Swedish data. We compare the results of annotating OTUs to family by phylogenetic placement (with EPA-NG and GAPPA) on a reference tree with the results of annotation to family by matching to the BOLD reference library (with SINTAX). **Comment**. For Swedish insects, the BOLD reference library has good coverage of the majority of families, resulting in good performance of annotation to higher taxa by matching sequences to the reference library. This is not the case for Madagascan insects, making phylogenetic placement in a reference tree more attractive. Here, we check that the phylogenetic placement methodology results in similar annotations to higher taxa for the Swedish data. Discrepancies include cases where there is disagreement in circumscription of families, or where families are not monophyletic, among other things. The discrepancies seem to primarily affect species-poor families.

**Fig. S15.**
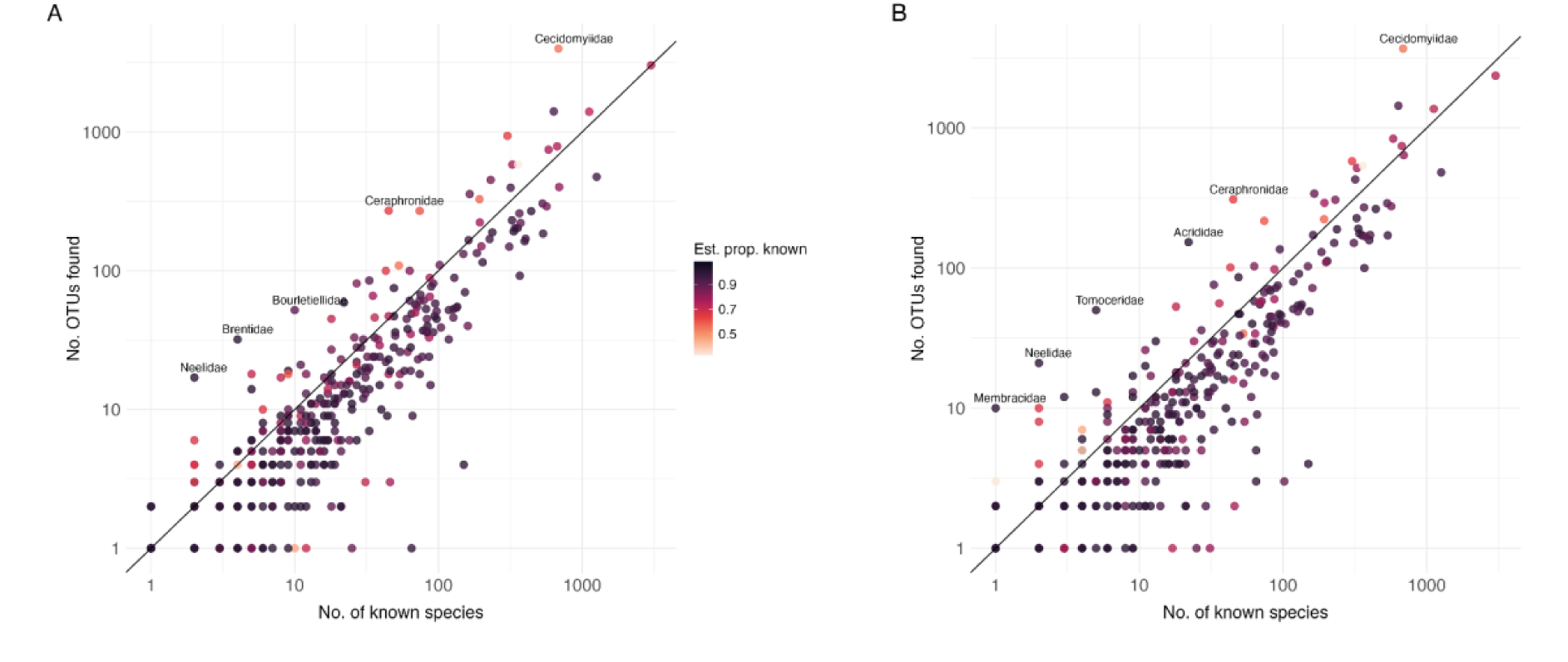
Validation of data quality using the Swedish insect fauna. Correlation between the number of sampled OTUs (Malaise samples (lysates) and litter samples) per family and the number of known species in the Swedish insect fauna (Ronquist et al. 2020). (A) Data annotated to family by matching to the BOLD reference library using sintax. (B) Data annotated to family by placing OTUs in the reference tree developed for this study. The color reflects the estimated proportion of unknown species remaining to be discovered in each family (Ronquist et al. 2020). The line corresponds to the number of OTUs expected if all the known Swedish species were included in the sample; families are expected to fall below this line if the estimated proportion of known species is high. **Comment**. Taxonomic annotation with the phylogenetic placement methodology used here (B), which is less sensitive to gaps in the reference data and thus more robust when analyzing poorly charted faunas, results in only a slight increase in false positives compared to matching to the BOLD reference library (A). Large numbers of ghost OTUs are still observed in some families that are well known (dark dots above the diagonal line), indicating that the noise filtering is ineffective for those taxa.

**Fig. S16.**
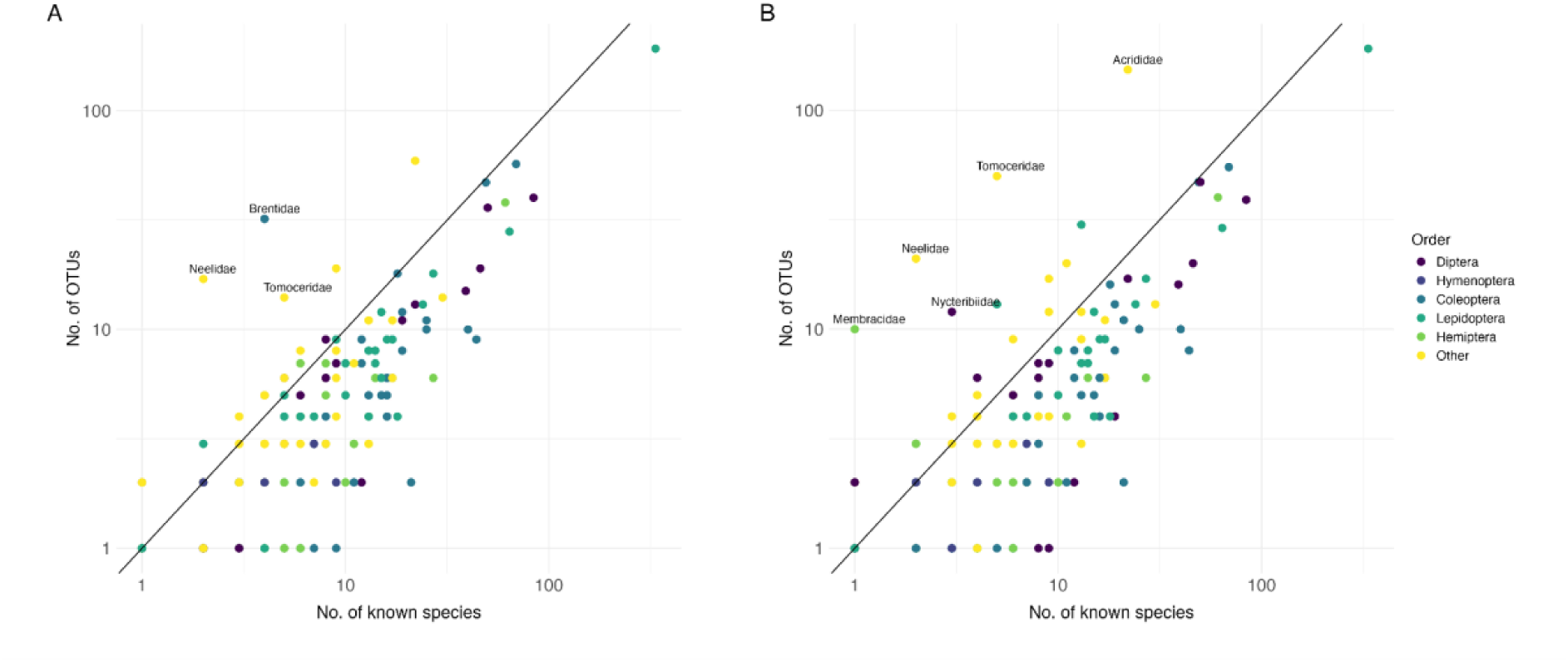
Validation of data quality for well-known families in the Swedish fauna. Comparison of estimated number of OTUs for well-known families (estimated proportion of known species ≥ 0.99) in the Swedish fauna against the number of known species (diagonal line). The colour indicates the order to which each family belongs. (A) Filtered data annotated to family by matching to the BOLD reference library using SINTAX. (B) Filtered data annotated to family by placing OTUs in the reference tree developed for this study using EPA-NG and GAPPA, without using the BOLD reference library. **Comment**: Despite the application of advanced noise filtering algorithms, a large number of spurious or ghost OTUs remain in a few families, such as Neelidae and Tomoceridae (Collembola). The problematic taxa tend to be relatively poor in species or belong to the minor orders. Matching to the BOLD reference library (A), which is generally fairly complete for the Swedish fauna analyzed here, helps in resolving some of these issues but is susceptible to others that do not cause problems in phylogenetic placement (B).

**Fig. S17.**
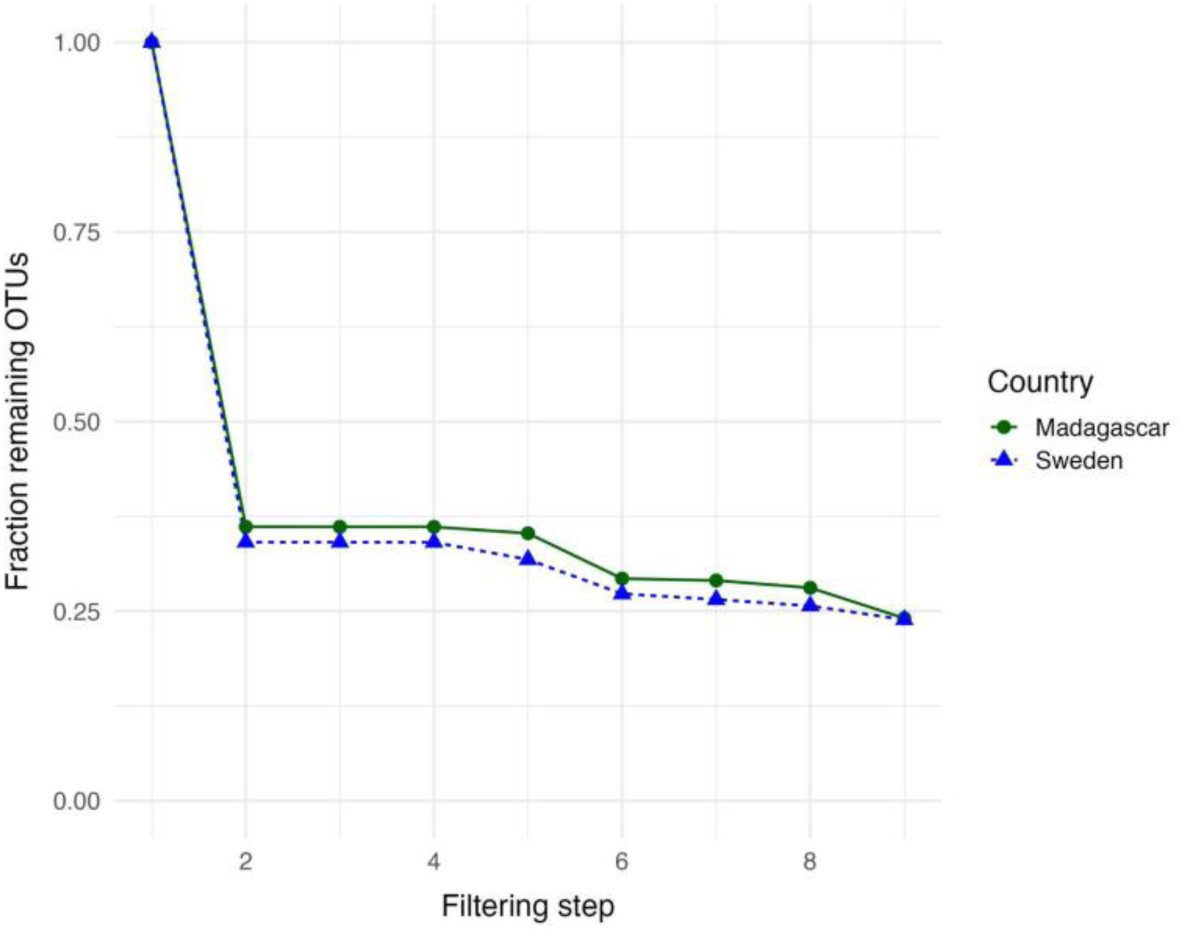
Performance of the filtering and annotation methodology on Swedish and Madagascan data. Filtering steps: (1) Unfiltered clusters from ampliseq and DADA2 pipeline. (2) NEEAT filtering. (3) Removal of contaminants based on control samples. (4) Removal of spike-ins. (5) Restriction to hexapods. (6) Removal of uncertain phylogenetic placements (aLWR ≤ 0.5). (7) Removal of OTUs with SINTAX BOLD placements conflicting at the order level. (8) Removal of OTUs with VSEARCH placements conflicting at the phylum level. (9) Restriction to OTUs assigned to family clade. **Comment**: The main filtering step, where most of the noise is removed, involves the removal of NUMTs, sequencing artefacts and other likely spurious OTUs by the NEEAT algorithm (step 2). Slightly fewer clusters are removed from the Madagascan data, presumably due to the shallower sampling per species, resulting in fewer artefacts being picked up. The shallower sampling also makes the recognition of artefacts by sample co-occurrence (the “echo” signal in NEEAT) less effective. Placing OTUs reliably in the reference tree (aLWR > 0.5; step 6) and to a family clade (step 9) is slightly less successful for Madagascar data than for Swedish data, presumably due to the reference tree not adequately covering the diversity of Madagascan insects

**Fig. S18.**
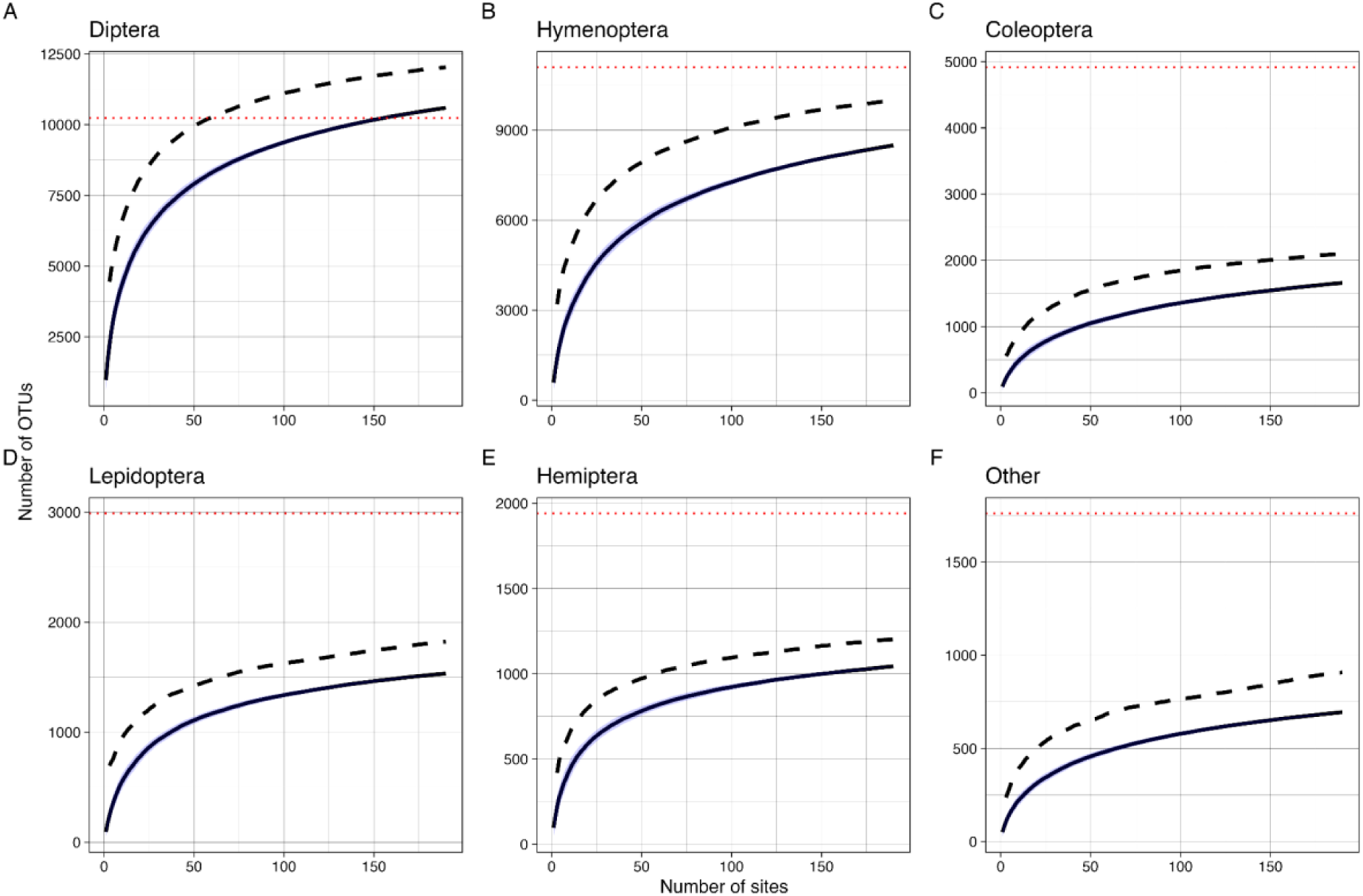
Species accumulation curves for the Swedish reference sample by order. The data from all habitat types (forest and non-forest) are included. Curves are based on the mean of 100 permutations; the blue region marks values within one standard deviation from the mean. The dashed line represents average Chao2 extrapolations from the randomly permuted data. The red dotted line represents the estimated number of species in each order (Ronquist et al. 2020). (A) Diptera. (B) Hymenoptera. (C) Coleoptera. (D) Lepidoptera. (E) Hemiptera. (F) Other orders.

**Fig. S19.**
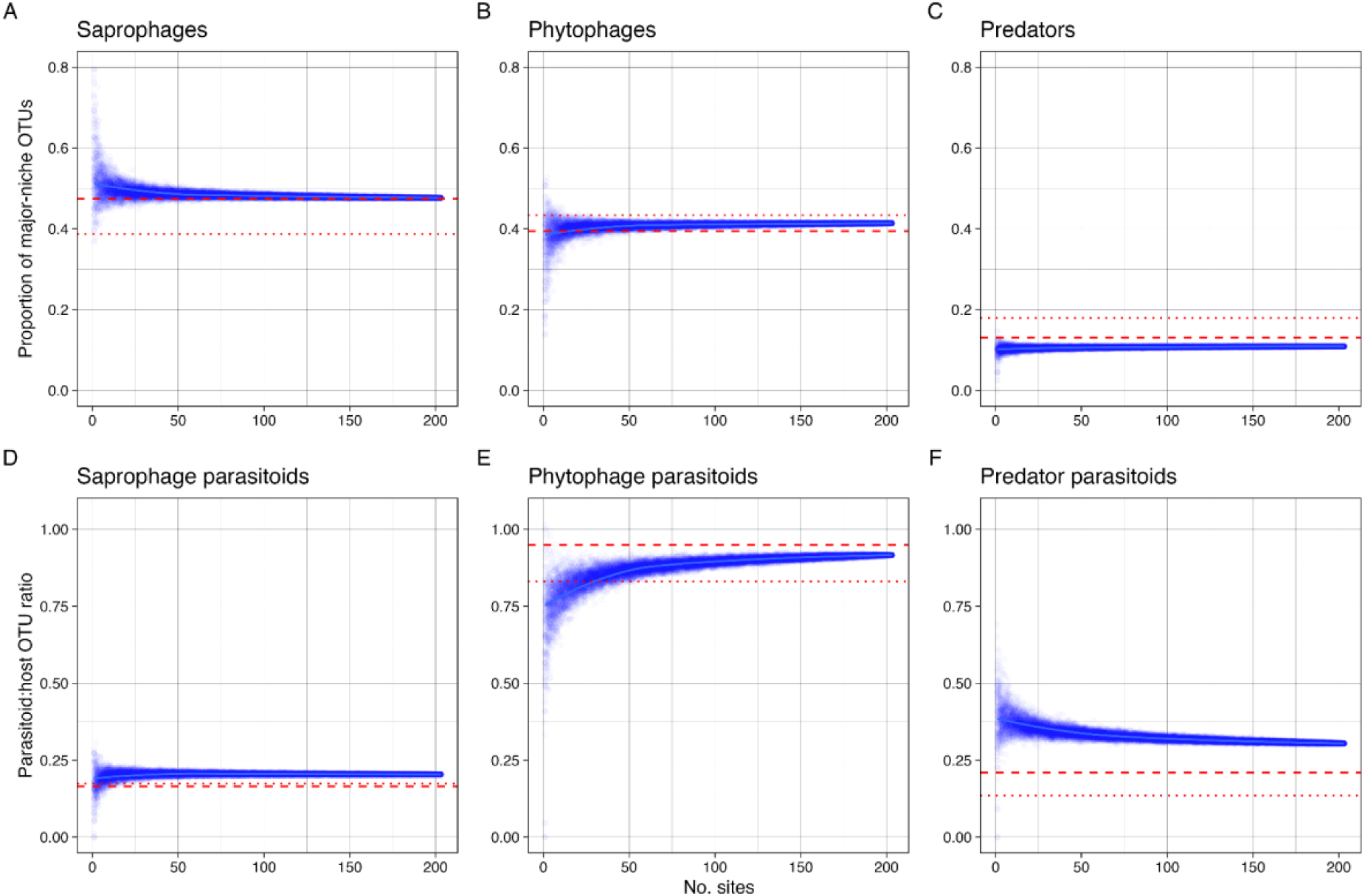
Spatial accumulation of niche composition across Swedish sites. The points represent 100 jackknife samples for each number of sites. The data include all sampled habitats (forest and non-forest). The red dotted lines indicate the taxonomic composition of the true fauna based on a previously published estimate (Ronquist et al. 2020); the red dashed lines indicate the revised estimate based on correcting the fraction of unknown species in Diptera and Hymenoptera. (A–C) Proportion of OTUs belonging to the three major feeding niches. (A) Saprophages. (B) Phytophages. (C) Predators. (D–F) Parasitoid:host OTU ratios for each of the three main communities. (D) Saprophage parasitoids. (E) Phytophage parasitoids. (F) Predator parasitoids.

**Fig. S20.**
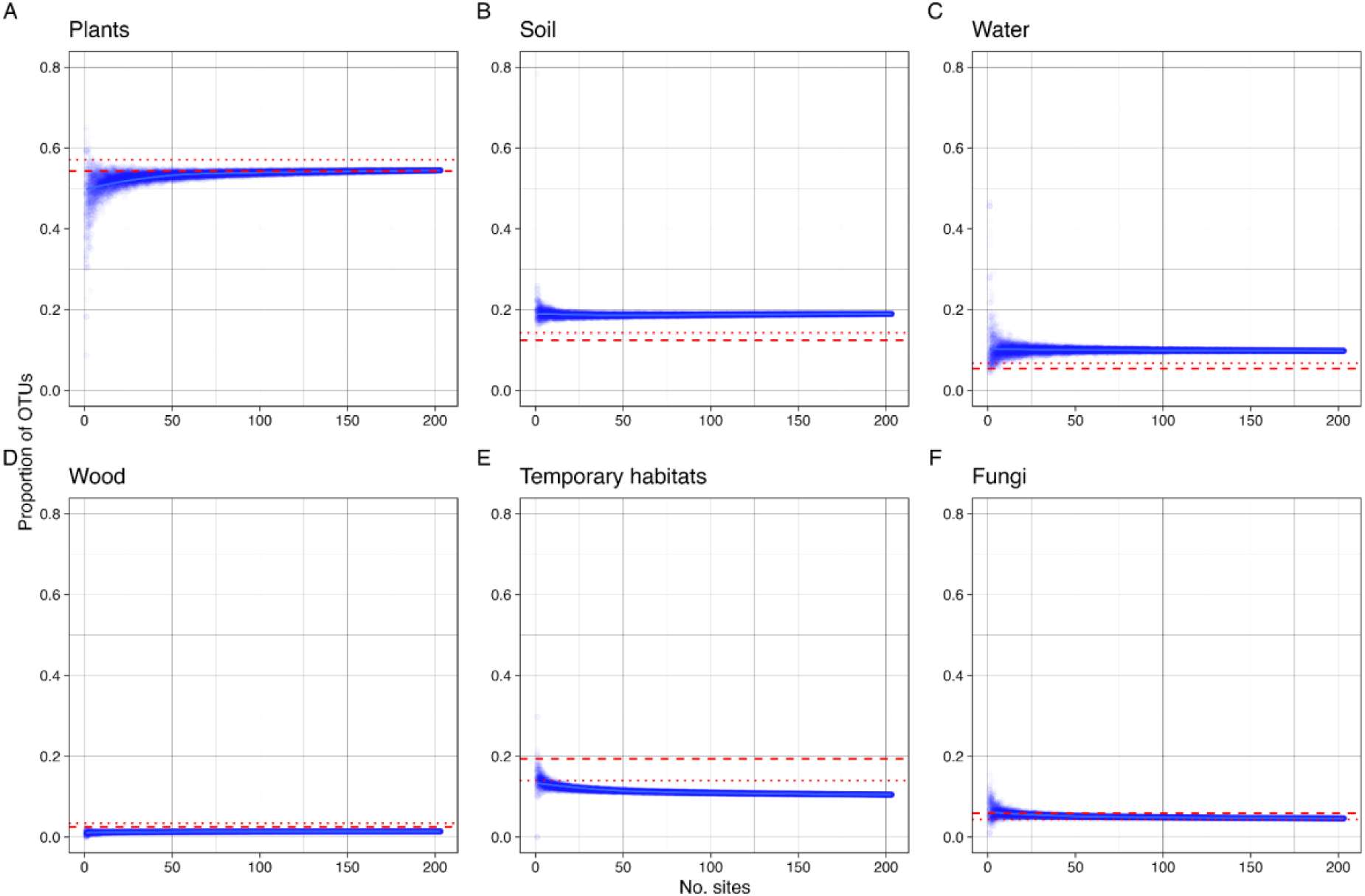
Spatial accumulation of microhabitat composition across sites. The points represent 100 jackknife samples for each number of sites. The data include all sampled habitats (forest and non-forest). The red dotted lines indicate the taxonomic composition of the true fauna based on a previously published estimate (Ronquist et al. 2020); the red dashed lines indicate the revised estimate based on correcting the fraction of unknown species in Diptera and Hymenoptera. (A) Species developing in or on plants. (B) Species developing in soil, litter or on the ground. (C) Species developing in water. (D) Species developing in wood. (E) Species developing in temporary microhabitats, such as carrion or dung. (F) Species developing in or on fungi.

**Fig. S21.**
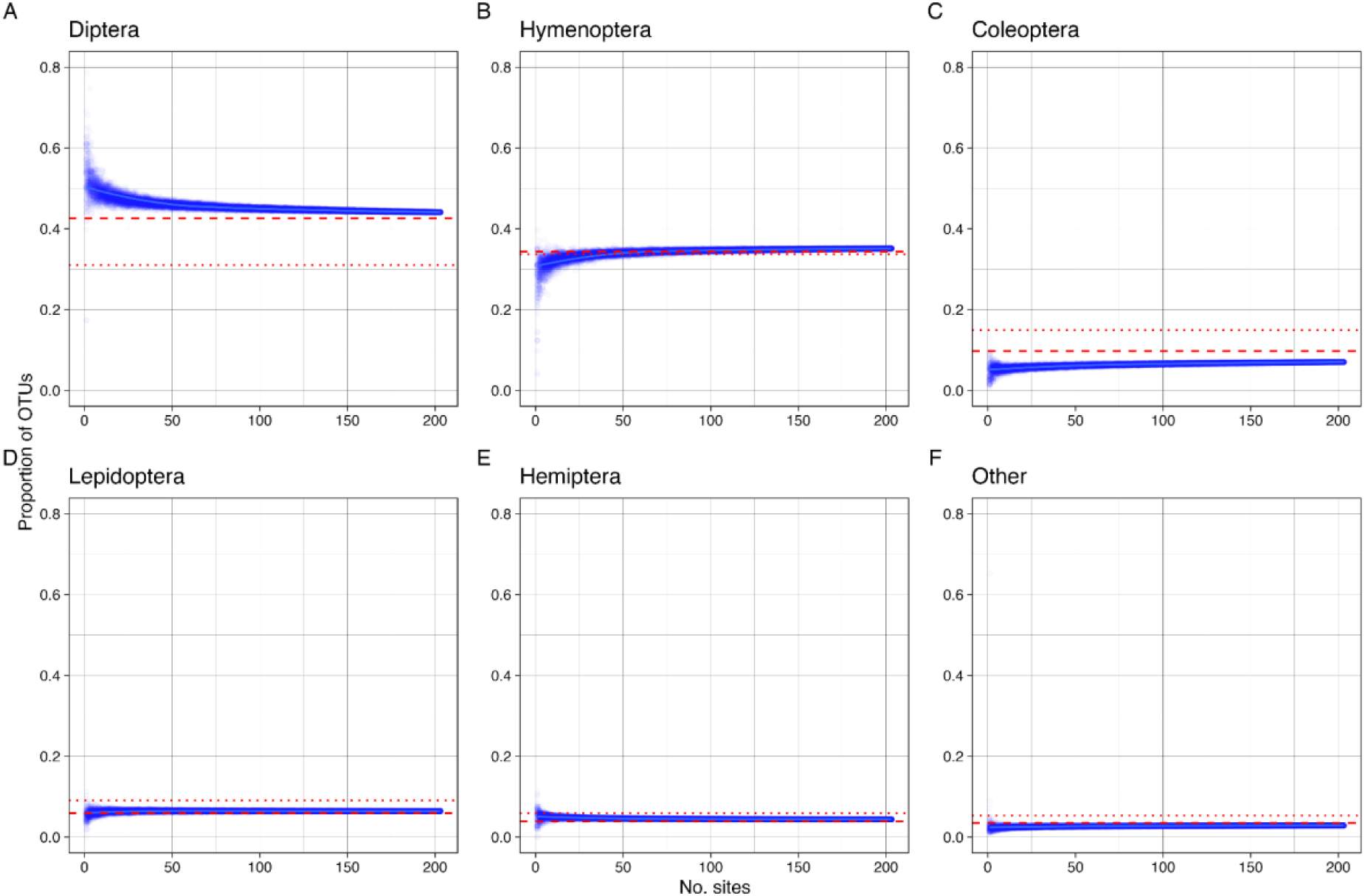
Spatial accumulation of taxonomic composition across Swedish sites. The points represent 100 jackknife samples for each number of sites. The data include all sampled habitats (forest and non-forest). The red dotted lines indicate the taxonomic composition of the true fauna based on a previously published estimate (Ronquist et al. 2020); the red dashed lines indicate the revised estimate based on correcting the fraction of unknown species in Diptera and Hymenoptera. (A) Diptera. (B) Hymenoptera. (C) Coleoptera. (D) Lepidoptera. (E) Hemiptera. (F) Remaining hexapod orders.

**Fig. S22.**
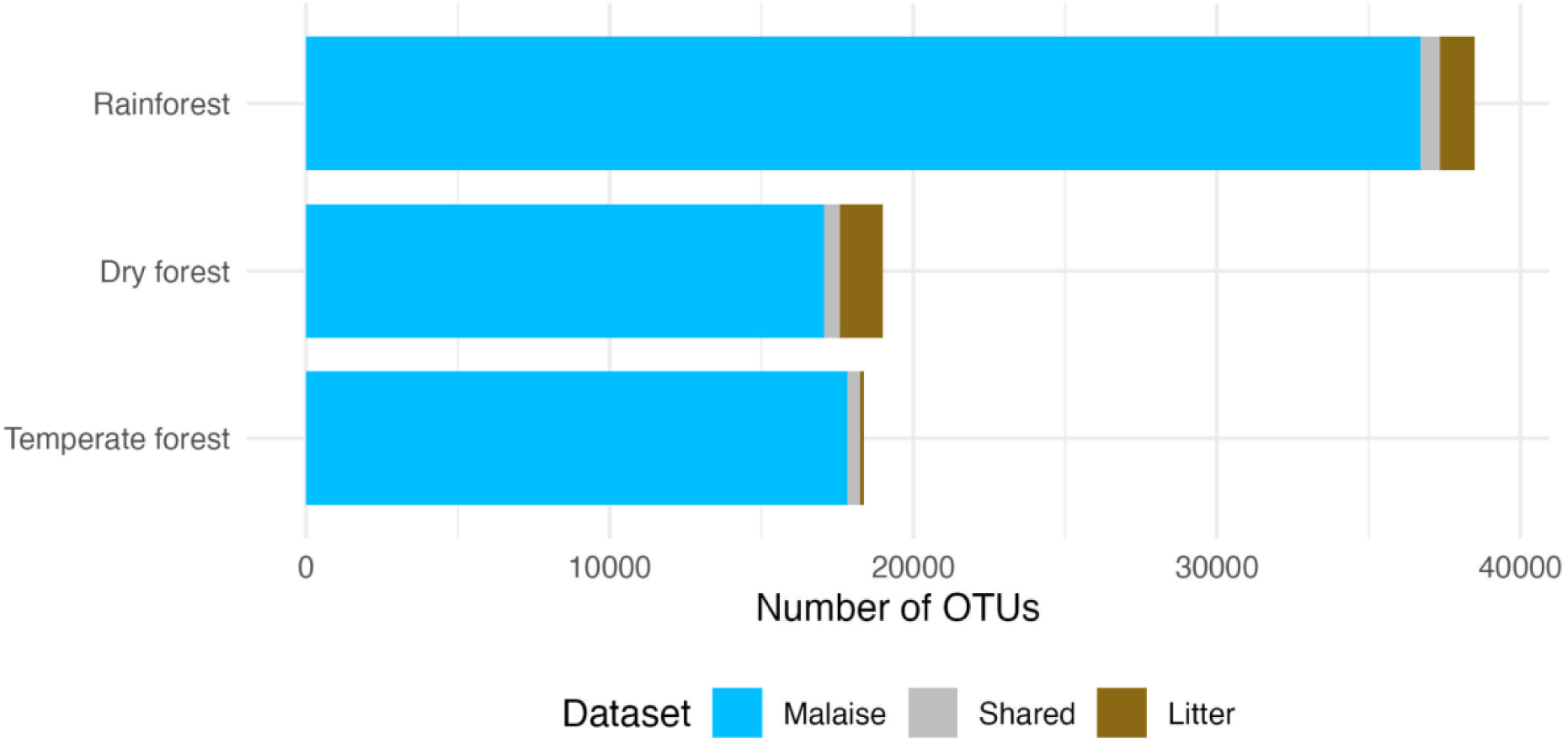
Contribution of different sampling methods (Malaise traps, litter samples, and shared) to the OTUs retrieved. The Madagascan and Swedish data are presented by forest type: Madagascan rainforest and dry forest, and Swedish temperate forest.

**Fig. S23.**
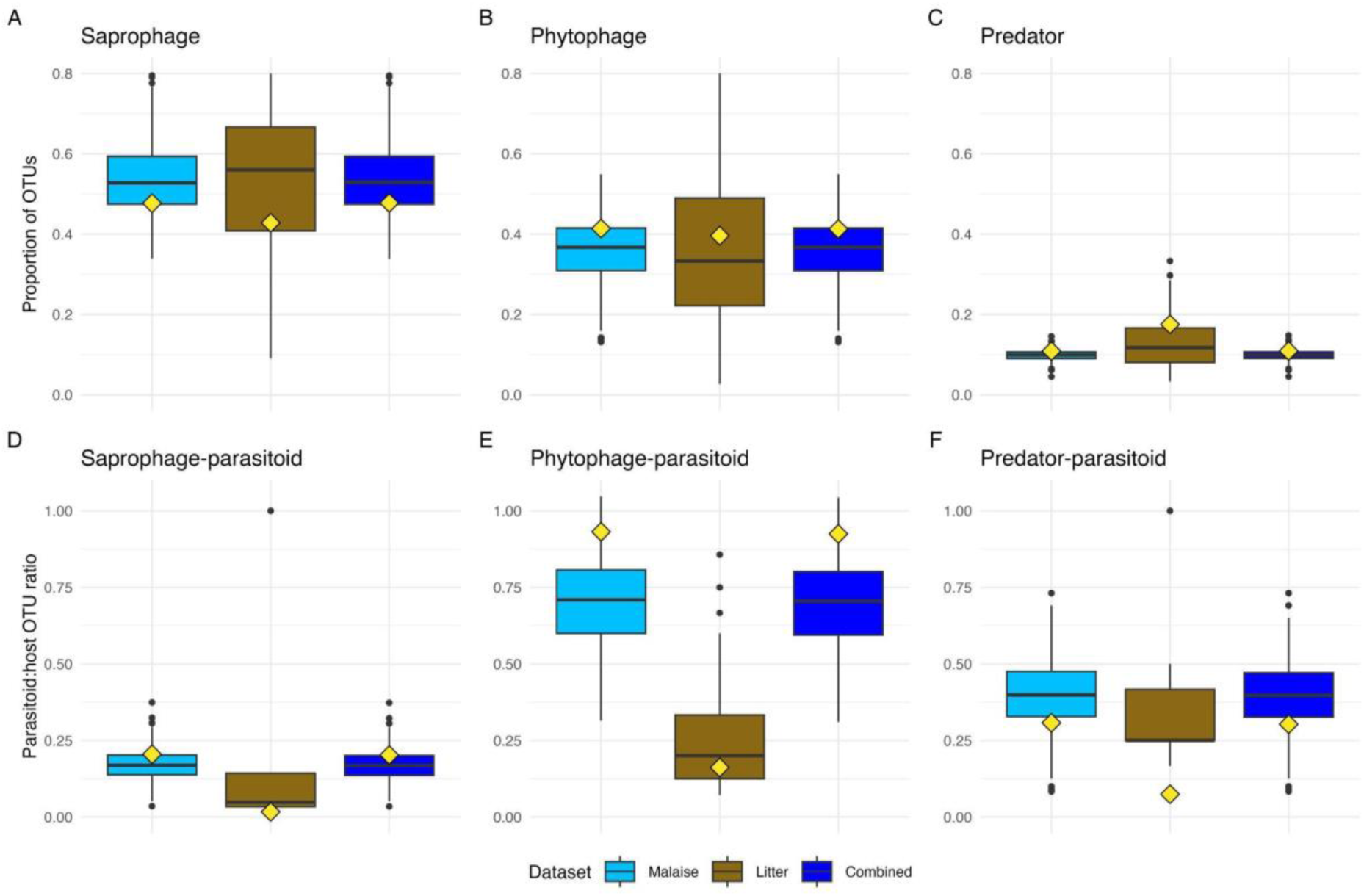
Contribution of different sampling methods (Malaise traps, litter samples, and combined) to the estimates of the niche composition of the Swedish fauna. Boxplots summarize the distribution of values across 12-month samples from each site. The red diamonds indicate the values to which the accumulation curves converge, and the distance between each diamond and the mean of the corresponding box plot indicates the rate at which the value changes with additional spatial sampling. (A–C) Proportion of OTUs belonging to the three major feeding niches for each trap type and combined. (D–F) Ditto for parasitoid to host OTU ratios.

**Fig. S24.**
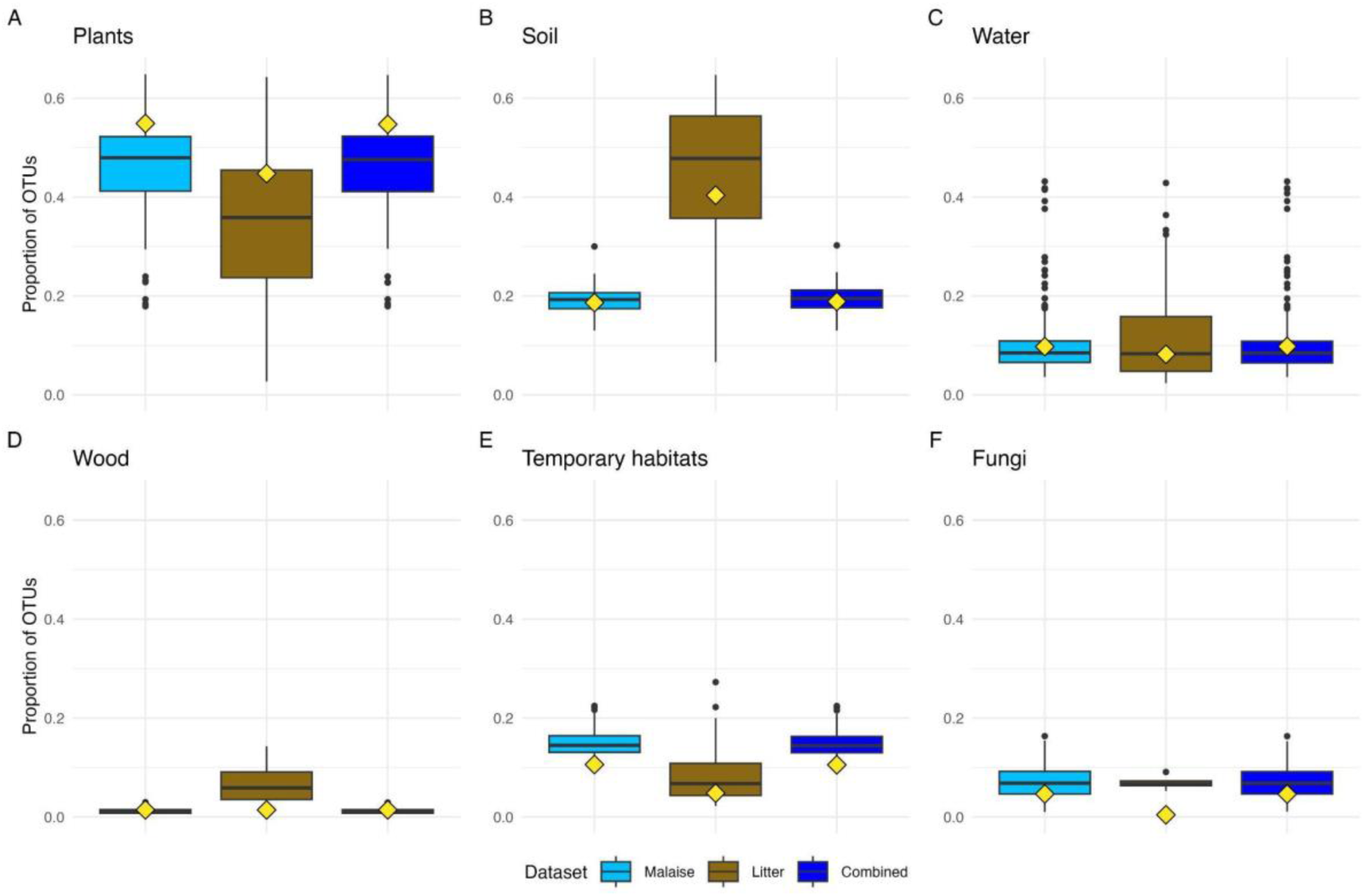
Contribution of different sampling methods (Malaise traps, litter samples, and combined) to the estimates of the microhabitat composition of the Swedish fauna. Boxplots summarize the distribution of values across 12-month samples from each site. The red diamonds indicate the values to which the accumulation curves converge, and the distance between each diamond and the mean of the corresponding box plot indicates the rate at which the value changes with additional spatial sampling.

**Fig. S25.**
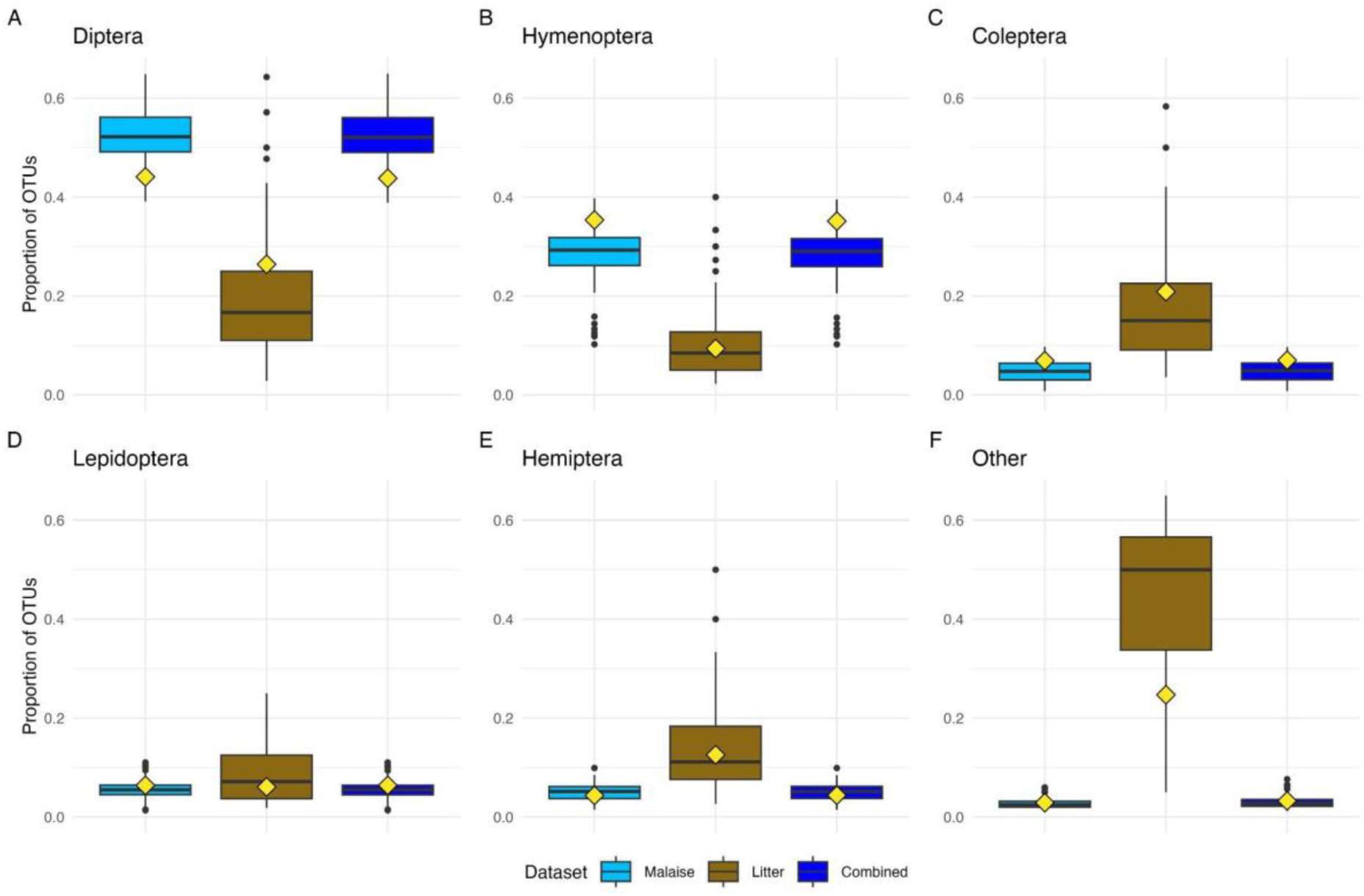
Contribution of different sampling methods (Malaise traps, litter samples, and combined) to the estimates of the taxonomic composition of the Swedish fauna. Boxplots summarize the distribution of values across 12-month samples from each site. The red diamonds indicate the values to which the accumulation curves converge, and the distance between each diamond and the mean of the corresponding box plot indicates the rate at which the value changes with additional spatial sampling.

**Fig. S26.**
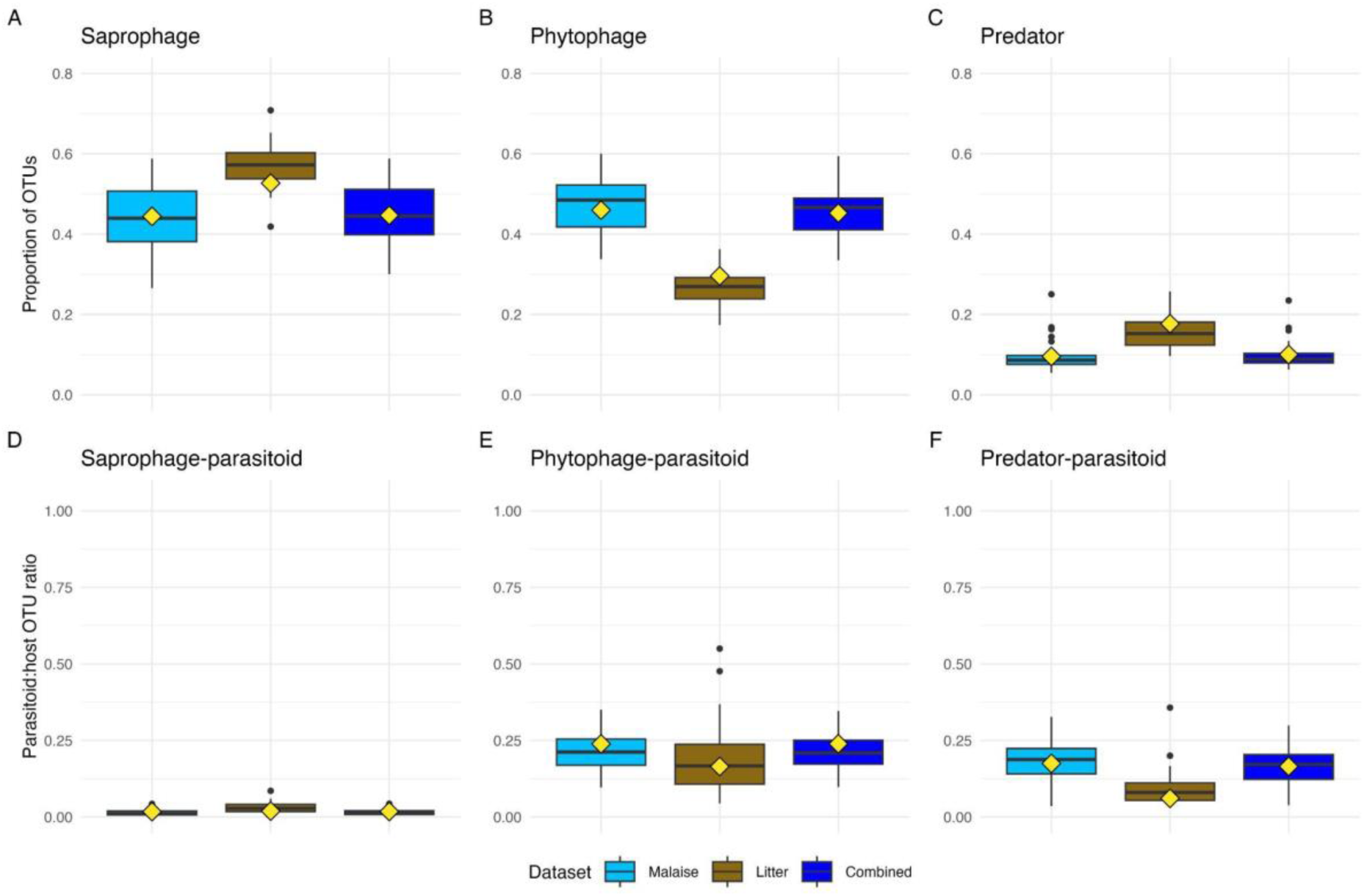
Contribution of different sampling methods (Malaise traps, litter samples, and combined) to the estimates of the niche composition of the Madagascan fauna. Boxplots summarize the distribution of values across 12-month samples from each site. The red diamonds indicate the values to which the accumulation curves converge, and the distance between each diamond and the mean of the corresponding box plot indicates the rate at which the value changes with additional spatial sampling. (A–C) Proportion of OTUs belonging to the three major feeding niches. (D–F) Parasitoid to host OTU ratios for each of the three main communities.

**Fig. S27.**
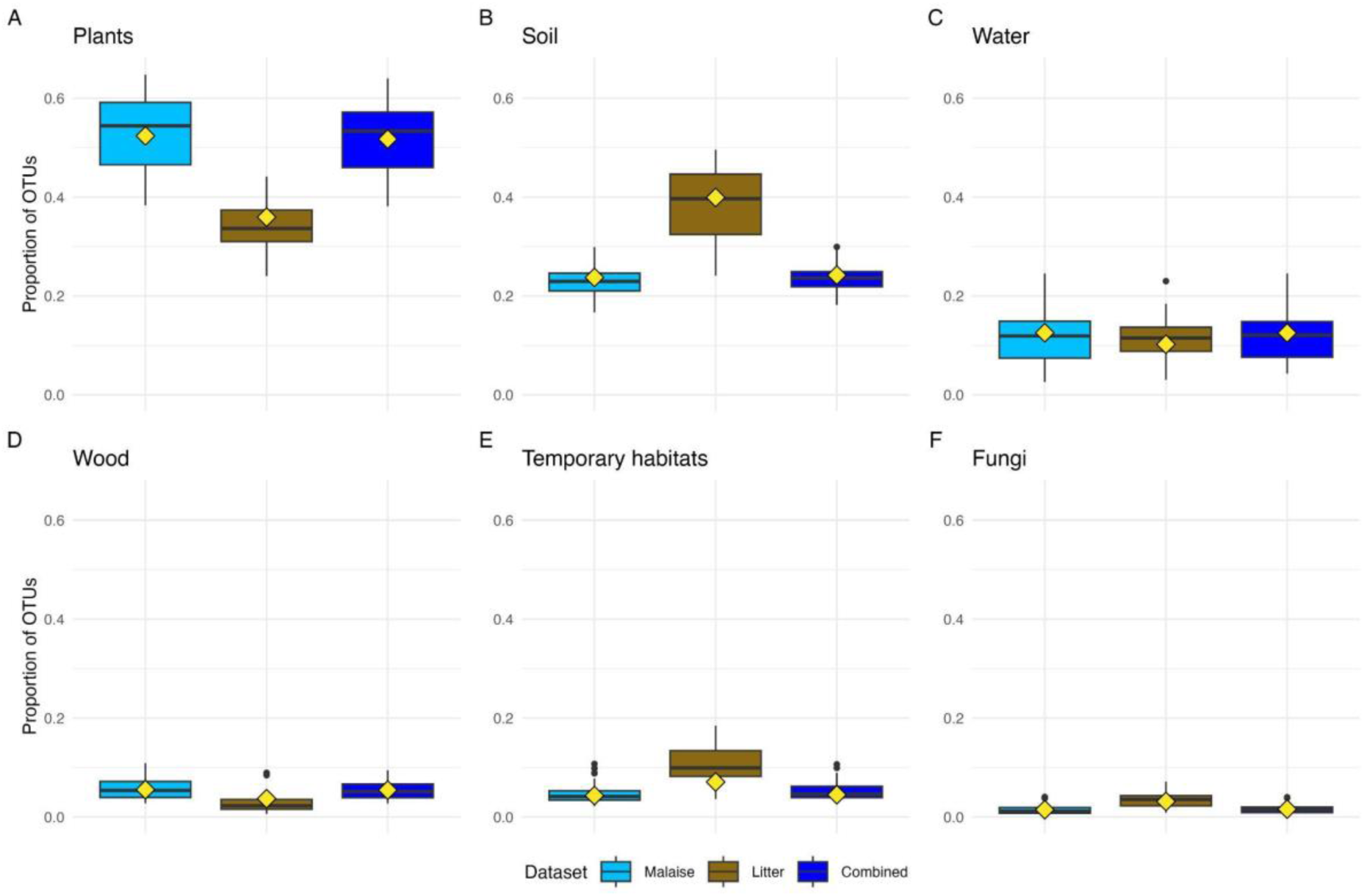
Contribution of different sampling methods (Malaise traps, litter samples, and combined) to the estimates of the microhabitat composition of the Madagascan fauna. Boxplots summarize the distribution of values across 12-month samples from each site. The red diamonds indicate the values to which the accumulation curves converge, and the distance between each diamond and the mean of the corresponding box plot indicates the rate at which the value changes with additional spatial sampling.

**Fig. S28.**
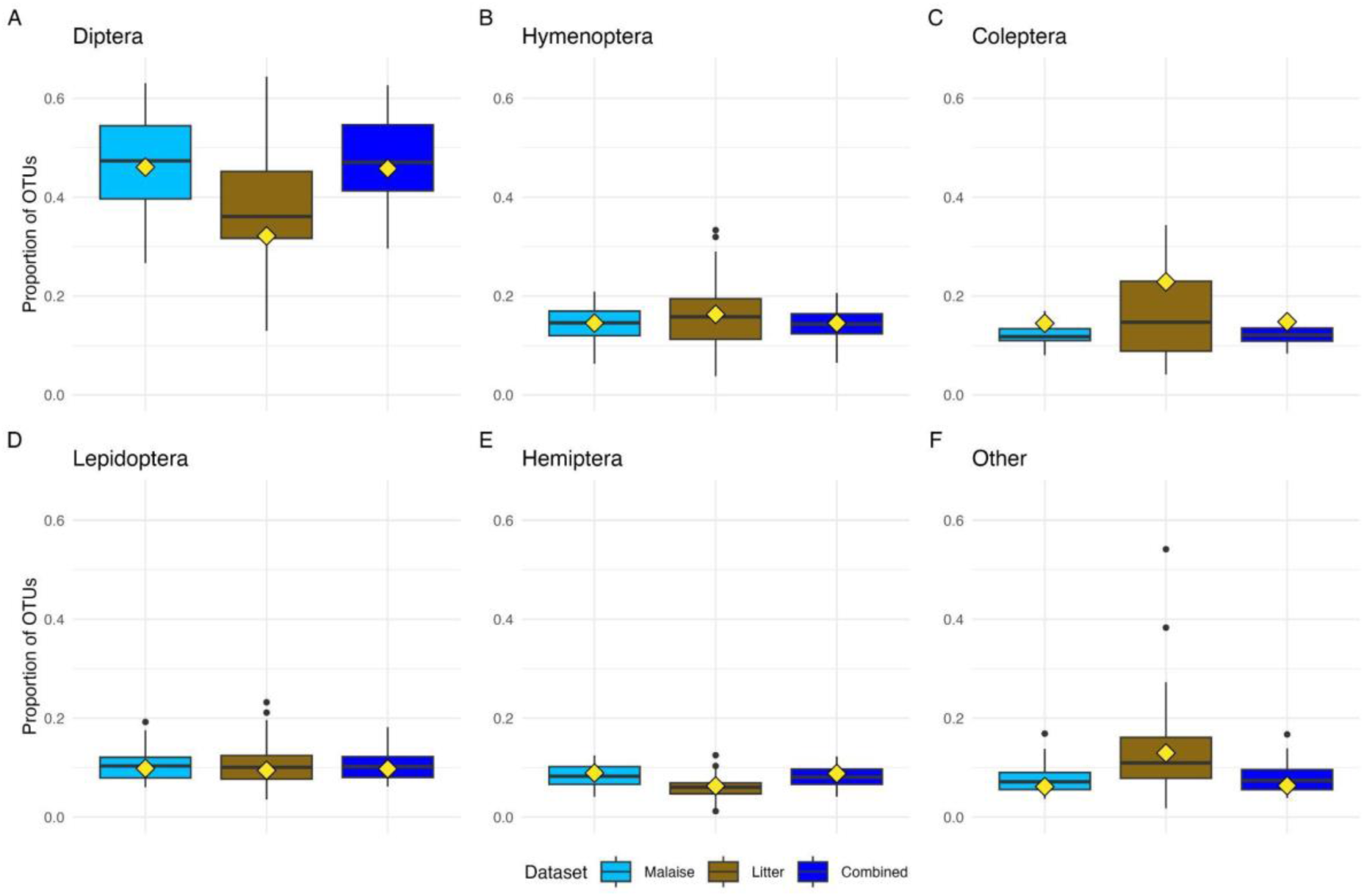
Contribution of different sampling methods (Malaise traps, litter samples, and combined) to the estimates of the taxonomic composition of the Madagascan fauna. Boxplots summarize the distribution of values across 12-month samples from each site. The red diamonds indicate the values to which the accumulation curves converge, and the distance between each diamond and the mean of the corresponding box plot indicates the rate at which the value changes with additional spatial sampling.

**Fig. S29.**
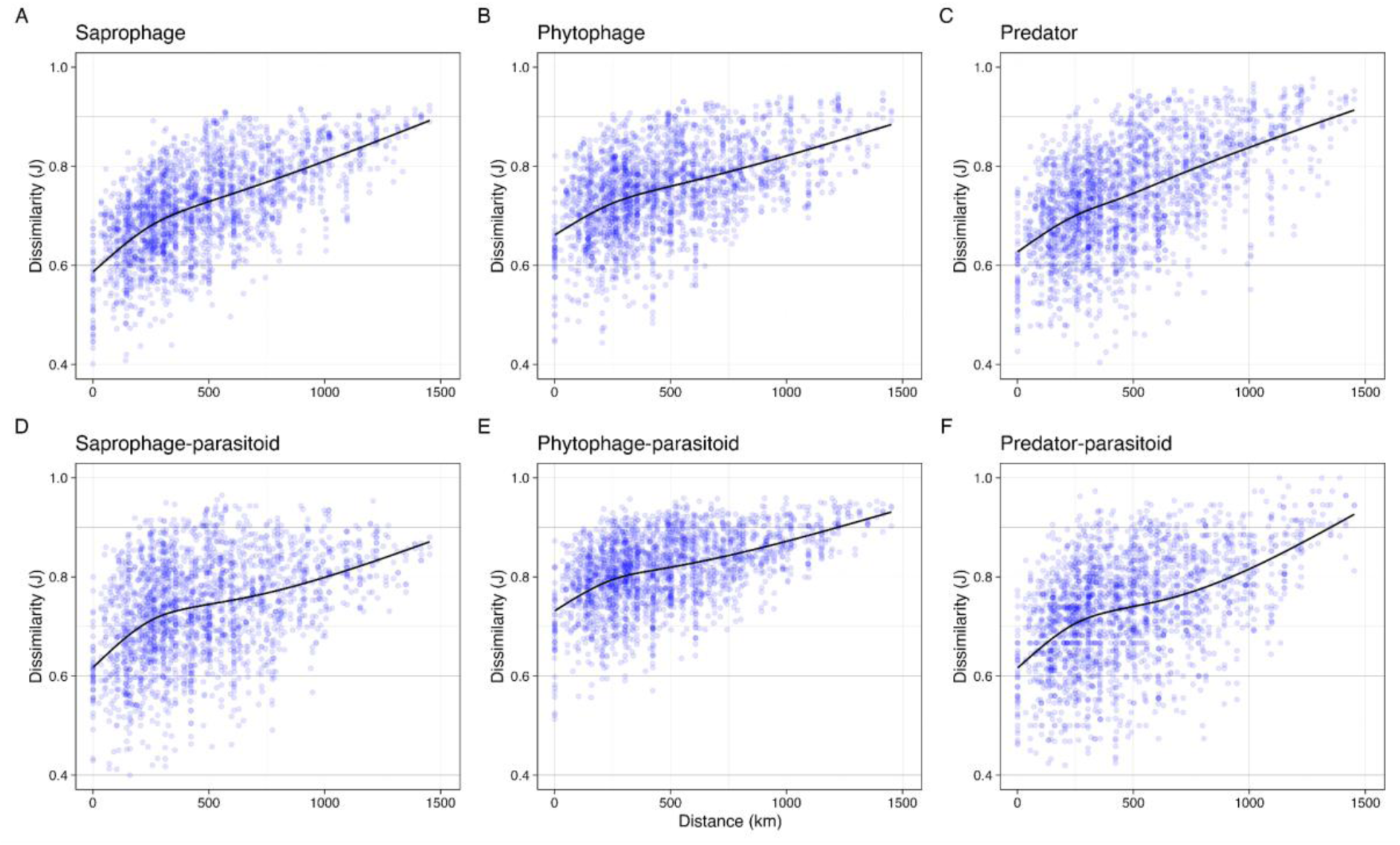
Spatial turnover of main feeding niches in the temperate forest sample. Turnover is expressed as Jaccard dissimilarity between pairs of trap sites. The fitted line is generated from a monotonic generalized additive model with five knots.

