## Supplementary Materials for "Global homeostatic forces shape the composition of a hyperdiverse tropical insect fauna"

### Supplementary Figures

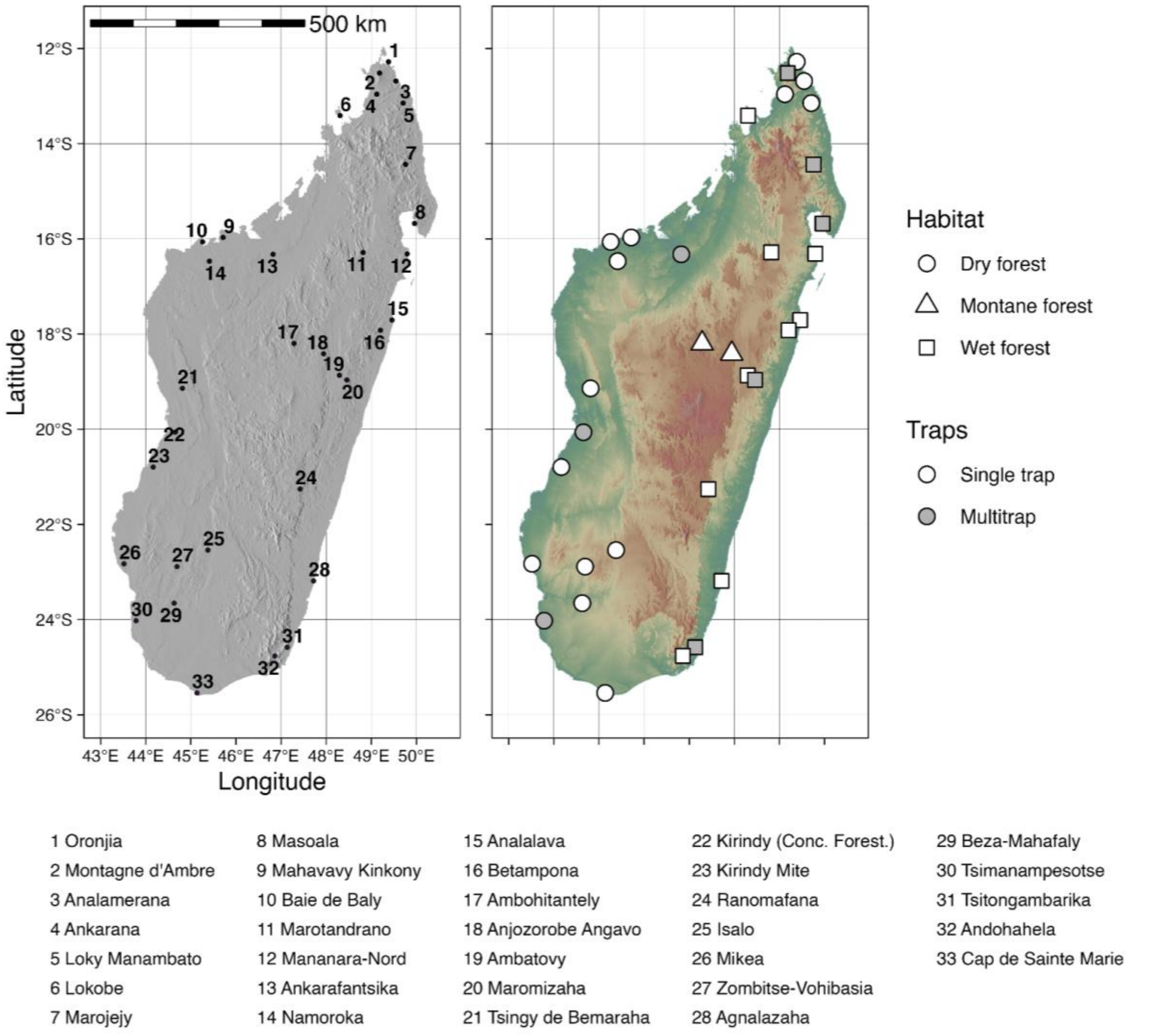


**Fig. S1. Sampling localities and forest types.** Malaise-trap sampling localities across Madagascar. Left: sampling localities. Right: the same localities, classified by forest habitat (dry forest, montane forest or wet forest) with point fill indicating trap type (single trap or multitrap, the latter sampled with 3–4 Malaise traps within a locality). All localities were situated in forested habitats within protected areas, managed by Madagascar National Parks or by other conservation bodies (e.g. Missouri Botanical Garden, Fanamby, Asity, GERP). In the forest-type analyses, montane forest ("montane rainforest") and wet forest ("rainforest") localities were grouped together as "rainforest" and contrasted with dry forest.

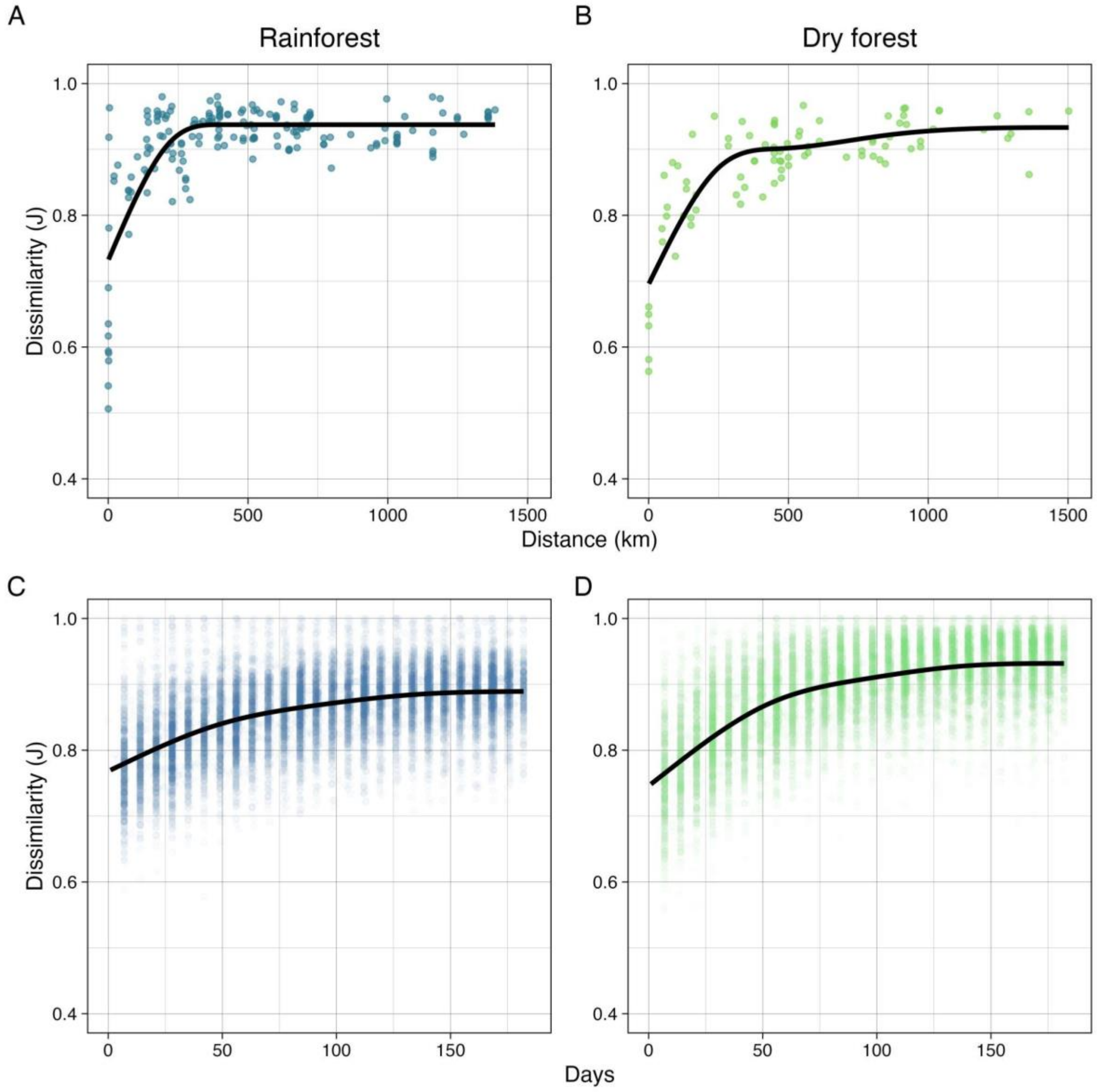


**Fig. S2. Spatial and temporal turnover by Madagascan forest type**. Turnover is expressed as Jaccard dissimilarity between sample pairs separated in space or time. The fitted line is generated from a monotonic generative additive model with five knots. (**A–B**) Spatial turnover across rainforest and dry forest sites, respectively. (**C–D**) Temporal turnover within sites for rainforest and dry forest sites, respectively.

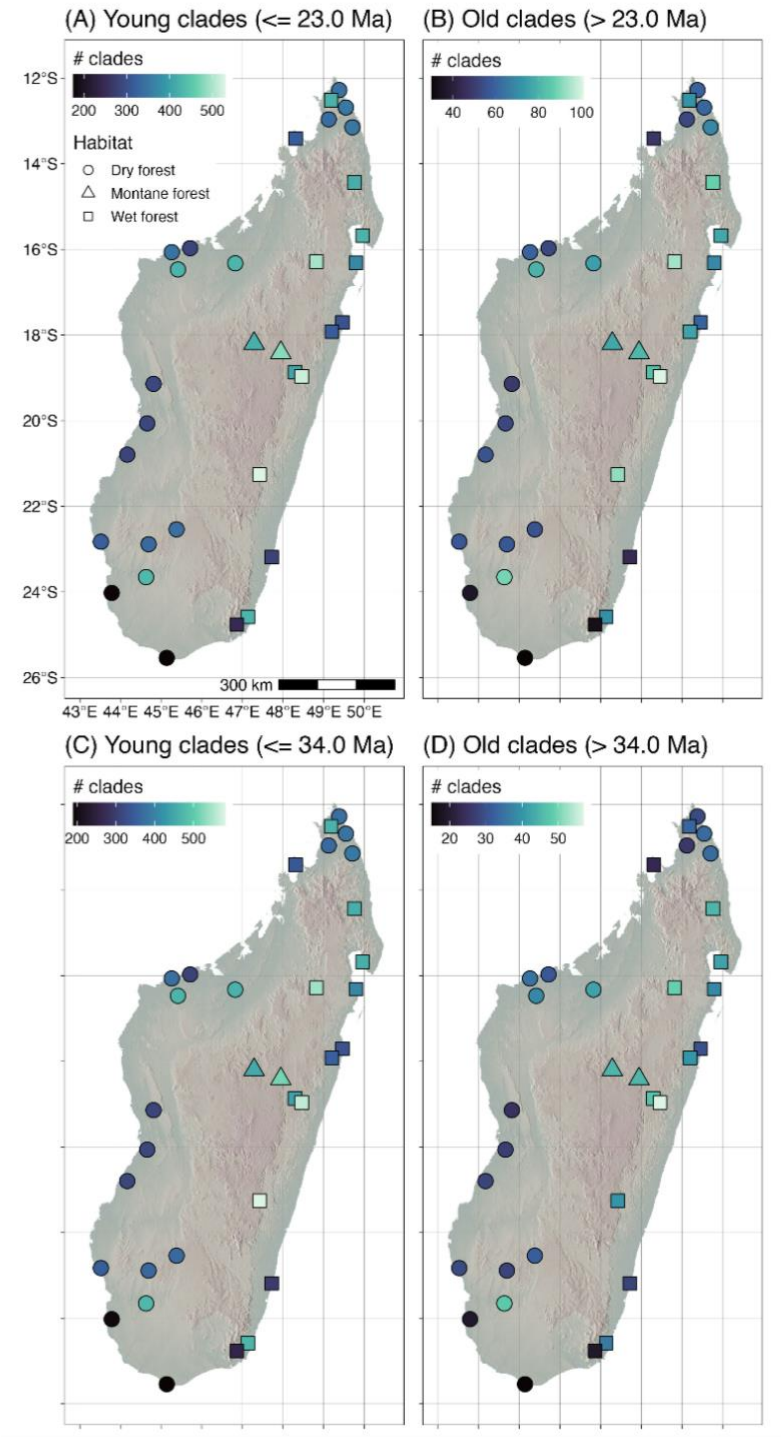


**Fig. S3. Distribution of old and young Madagascar lineages**. The number of Madagascar radiations (≥ 5 sampled species) represented at each site, subdivided by age. (A) Radiations dating from the Miocene or later (age ≤ 23 Ma). (B) Radiations predating the end of the Oligocene (age > 23 Ma). (C) Radiations dating from the Oligocene or later (age ≤ 34 Ma). (D) Radiations predating the end of the Eocene (age > 34 Ma). For multitrap locations, we show the median value across traps.

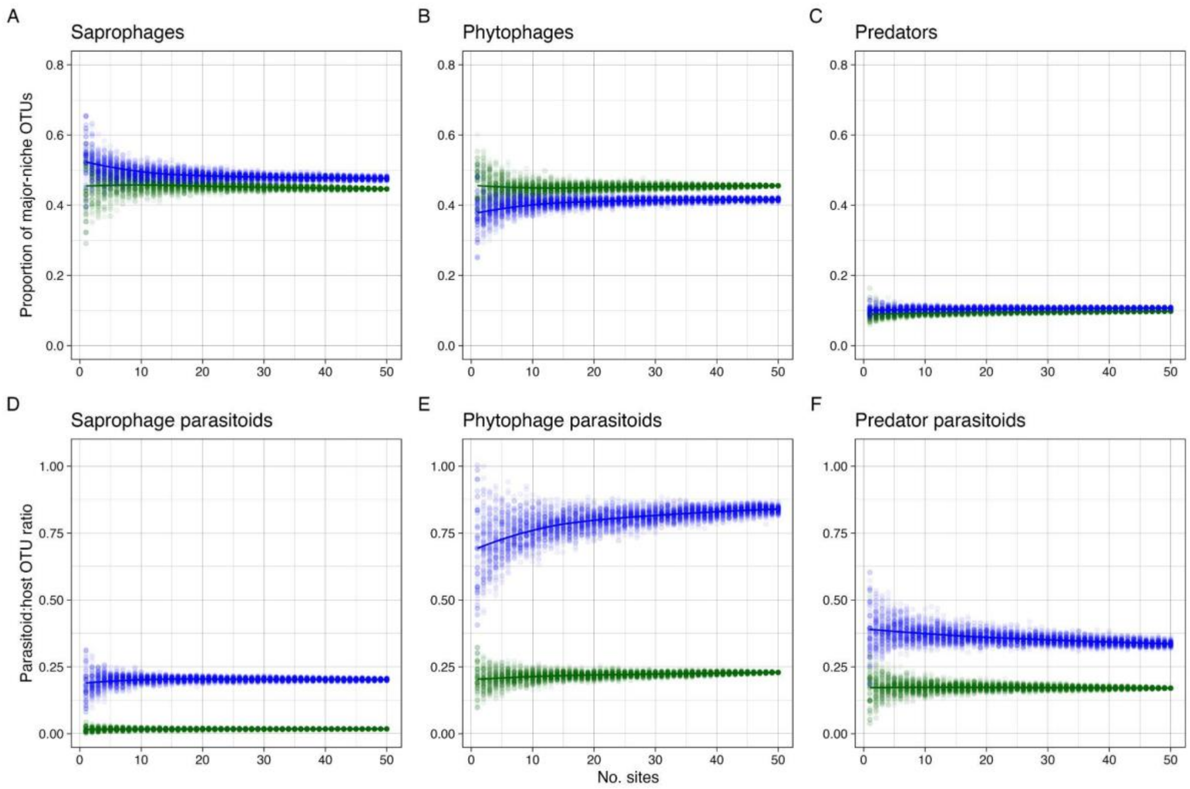


**Fig. S4. Spatial accumulation across sites of niche composition**. The points represent 100 random samples without replacement for each number of sites. Madagascan data in green, temperate forest reference data in blue. (A–C) Proportion of OTUs belonging to the three major feeding niches. (A) Saprophages. (B) Phytophages. (C) Predators. (D–F) Parasitoid:host OTU ratios for saprophage parasitoids (D), phytophage parasitoids (E), and predator parasitoids (F).

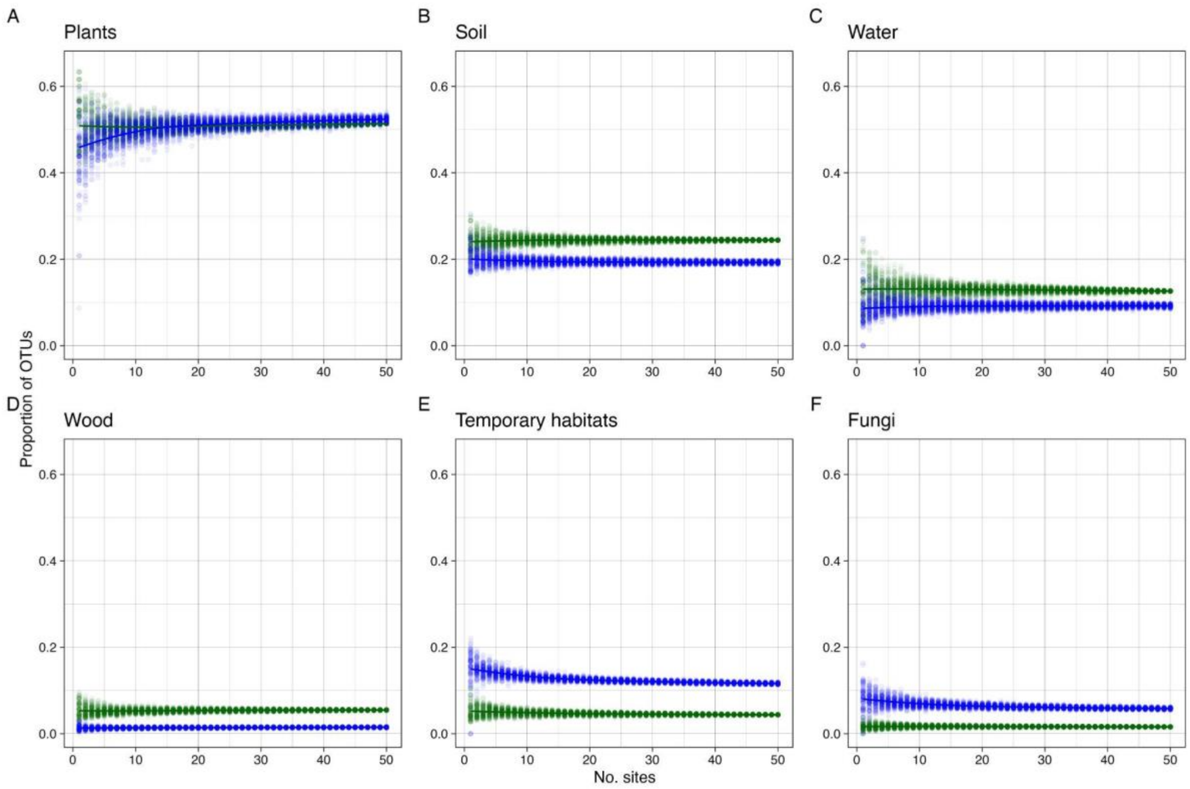


**Fig. S5. Spatial accumulation of microhabitat composition across sites**. The points represent 100 random samples without replacement for each number of sites. Madagascan data in green, temperate forest reference data in blue. (A) Species developing in or on plants. (B) Species developing in soil, litter or on the ground. (C) Species developing in water. (D) Species developing in wood. (E) Species developing in temporary microhabitats, such as carrion or dung. (F) Species developing in or on fungi.

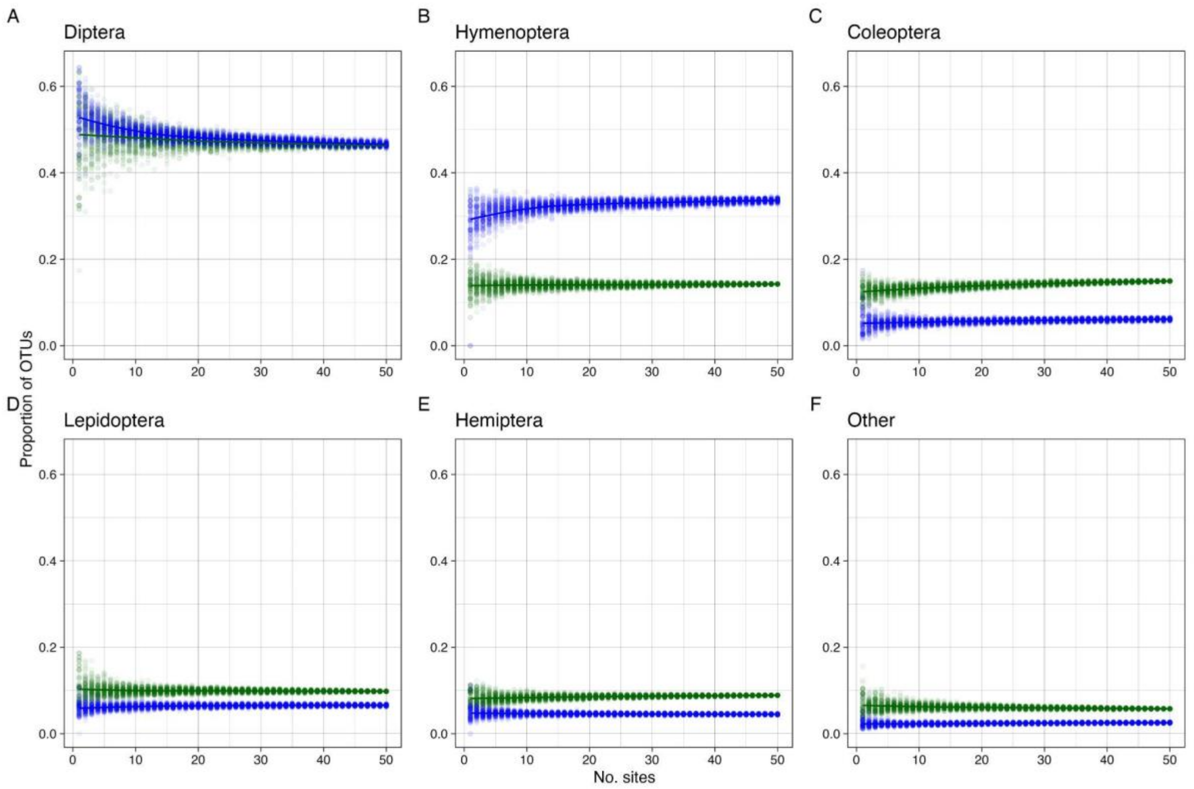


**Fig. S6. Spatial accumulation of taxonomic composition across sites**. The points represent 100 random samples without replacement for each number of sites. Madagascan data in green, temperate forest reference data in blue. (A) Diptera. (B) Hymenoptera. (C) Coleoptera. (D) Lepidoptera. (E) Hemiptera. (F) Remaining hexapod orders. Note that the y-scale is different in plots D–F to display the differences among the less diverse taxonomic groups more clearly.

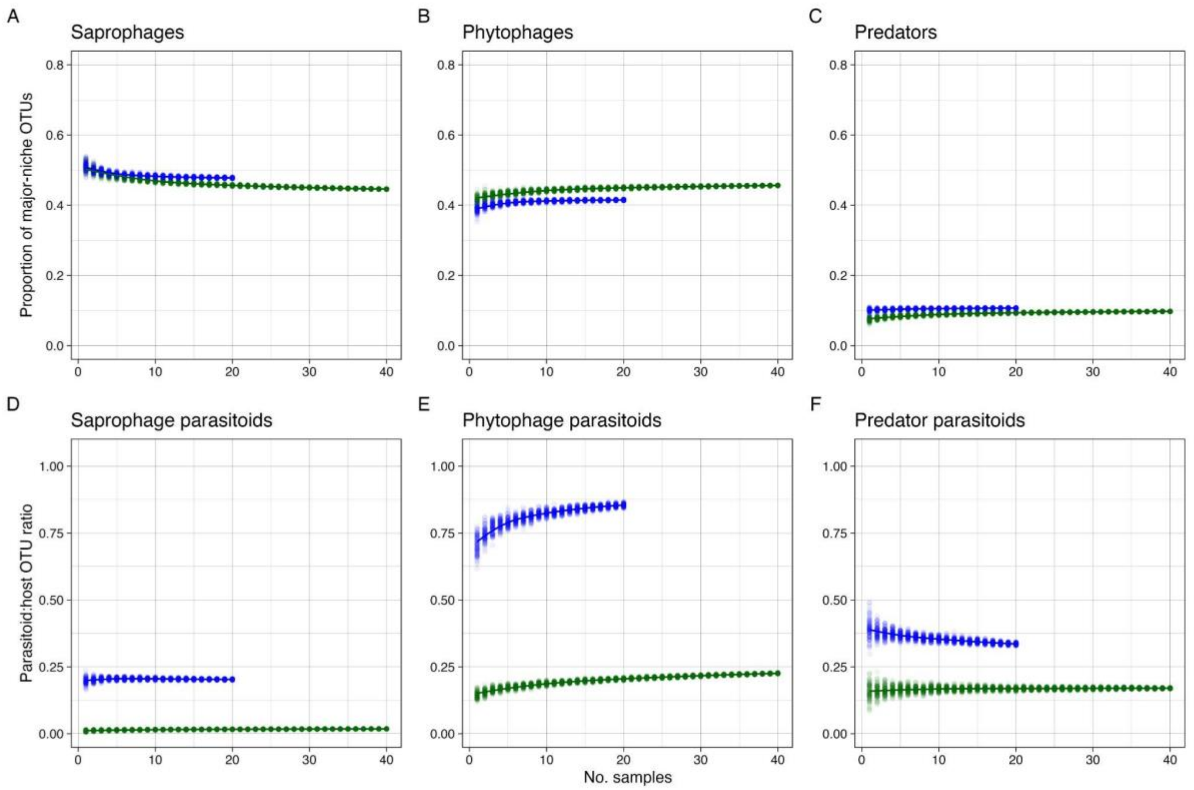


**Fig. S7. Temporal accumulation of niche composition across sites**. The points represent 100 random samples without replacement for each number of temporal samples (weekly samples in Madagascar, weekly to monthly samples in Sweden depending on season) from the studied trap sites. Madagascan data in green, temperate forest reference data in blue. (A–C) Proportion of OTUs belonging to the three major feeding niches. (A) Saprophages. (B) Phytophages. (C) Predators. (D–F) Parasitoid:host OTU ratios for each of the three main communities. (D) Saprophage parasitoids. (E) Phytophage parasitoids. (F) Predator parasitoids.

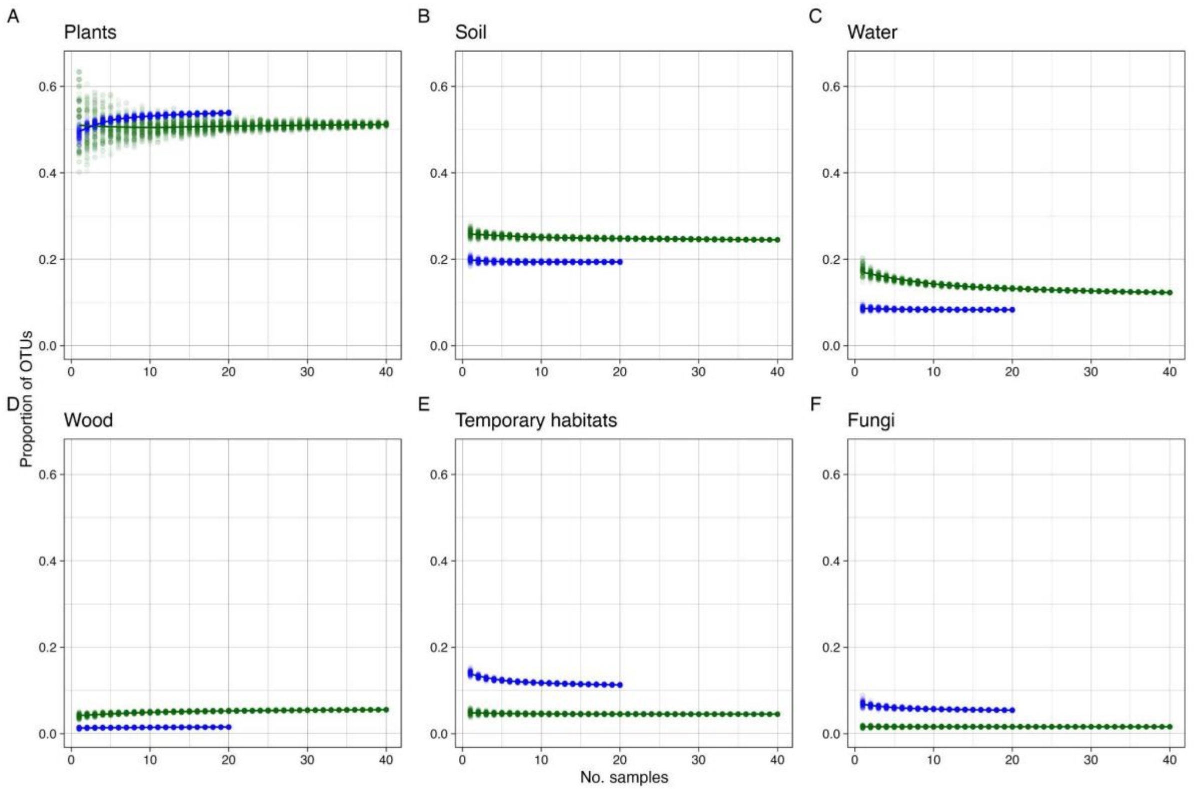


**Fig. S8. Temporal accumulation of microhabitat composition across sites**. The points represent 100 random samples without replacement for each number of temporal samples (weekly samples in Madagascar, weekly to monthly samples in Sweden depending on season) from the studied trap sites. Madagascan data in green, temperate forest reference data in blue. (A) Species developing in or on plants. (B) Species developing in soil, litter or on the ground. (C) Species developing in water. (D) Species developing in wood. (E) Species developing in temporary microhabitats, such as carrion or dung. (F) Species developing in or on fungi.

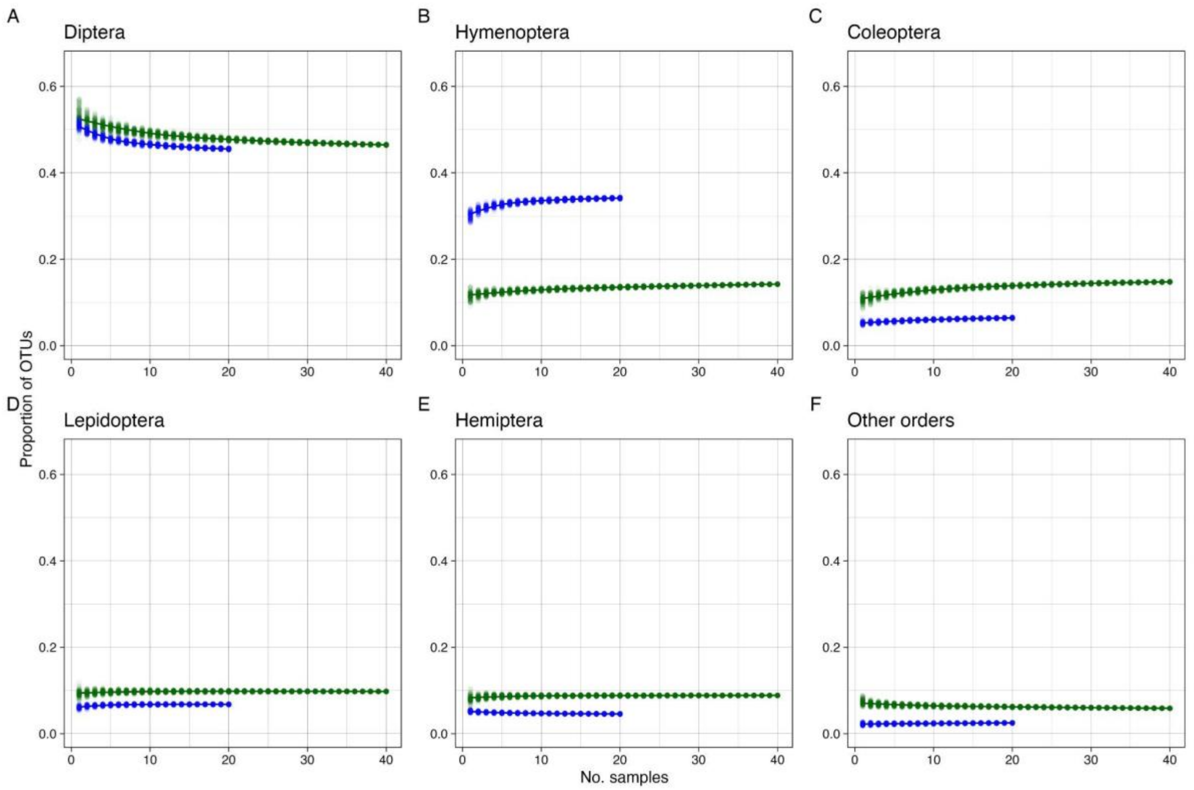


**Fig. S9. Temporal accumulation of taxonomic composition across sites**. The points represent 100 random samples without replacement for each number of temporal samples (weekly samples in Madagascar, weekly to monthly samples in Sweden depending on season) from the studied trap sites. Madagascan data in green, temperate forest reference data in blue. (A) Diptera. (B) Hymenoptera. (C) Coleoptera. (D) Lepidoptera. (E) Hemiptera. (F) Remaining hexapod orders.

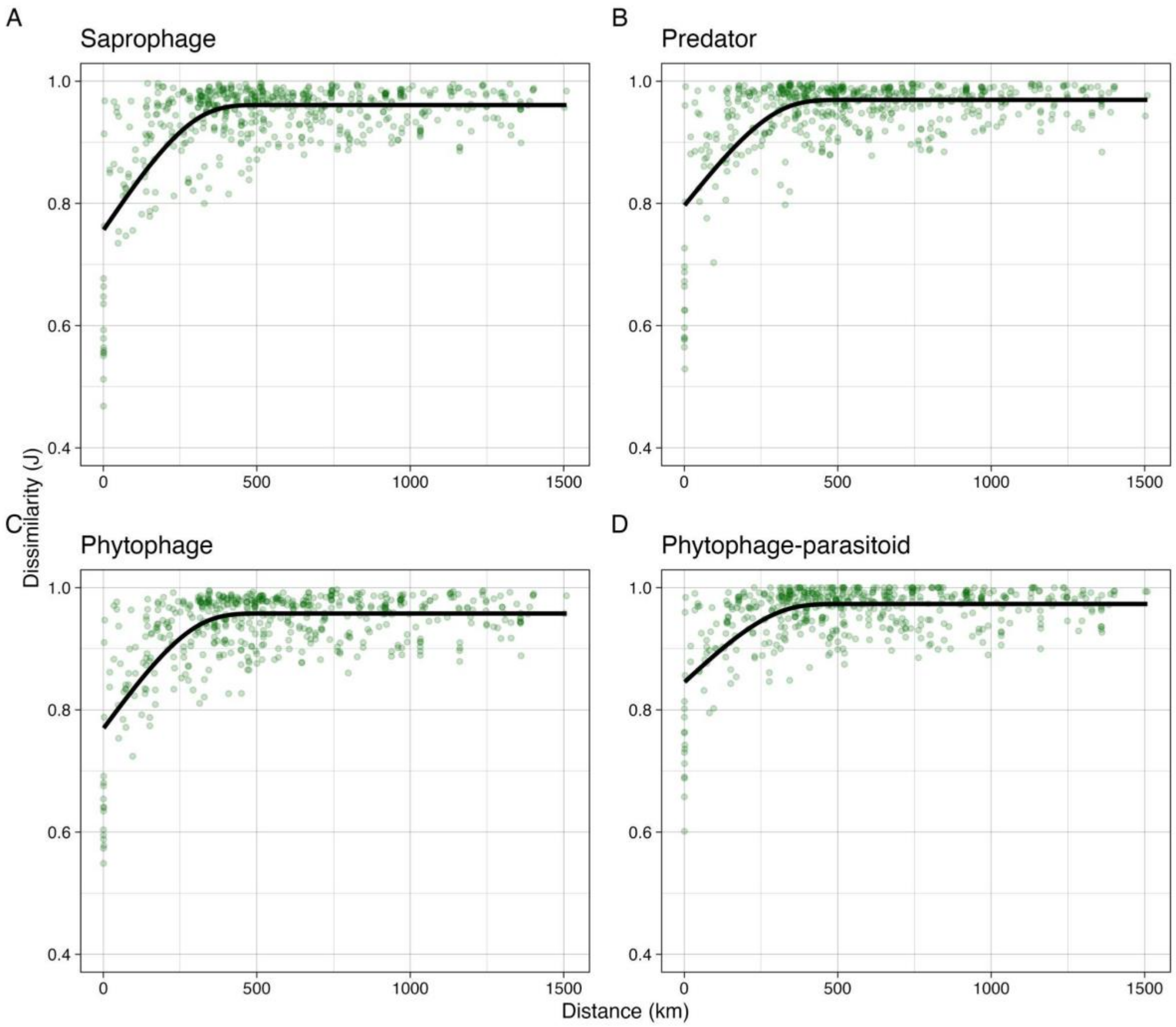


**Fig. S10. Spatial turnover of the three main feeding niches and of phytophage parasitoids in the Madagascan sample**. Turnover is expressed as Jaccard dissimilarity between pairs of trap sites. The fitted line is generated from a monotonic generalized additive model with five knots.

**Comment**. There are too few data points to meaningfully plot the spatial turnover for saprophage parasitoids and predator parasitoids. Note that the spatial turnover of phytophage parasitoids (**C**) is similar to that of their phytophage hosts (**D**). Compared to the temperate reference forest, the Madagascan forests are characterized by lower species diversity of parasitoids but similar parasitoid loads in the phytophage community. This suggests that Madagascan parasitoids have wider host ranges. An alternative explanation that we regard as less likely is that parasitoid loads are more unevenly distributed across host species in Madagascar. The wider host ranges in Madagascar could be achieved by parasitoids expanding their distribution ranges so that they can attack a larger set of closely related hosts; closely related host species are likely to vicariate with each other over a larger geographical range. It could also be achieved by parasitoids attacking a wider range of local hosts that are likely less closely related on average. The similarity in spatial turnover between hosts and parasitoids suggests that the latter is the more likely explanation.

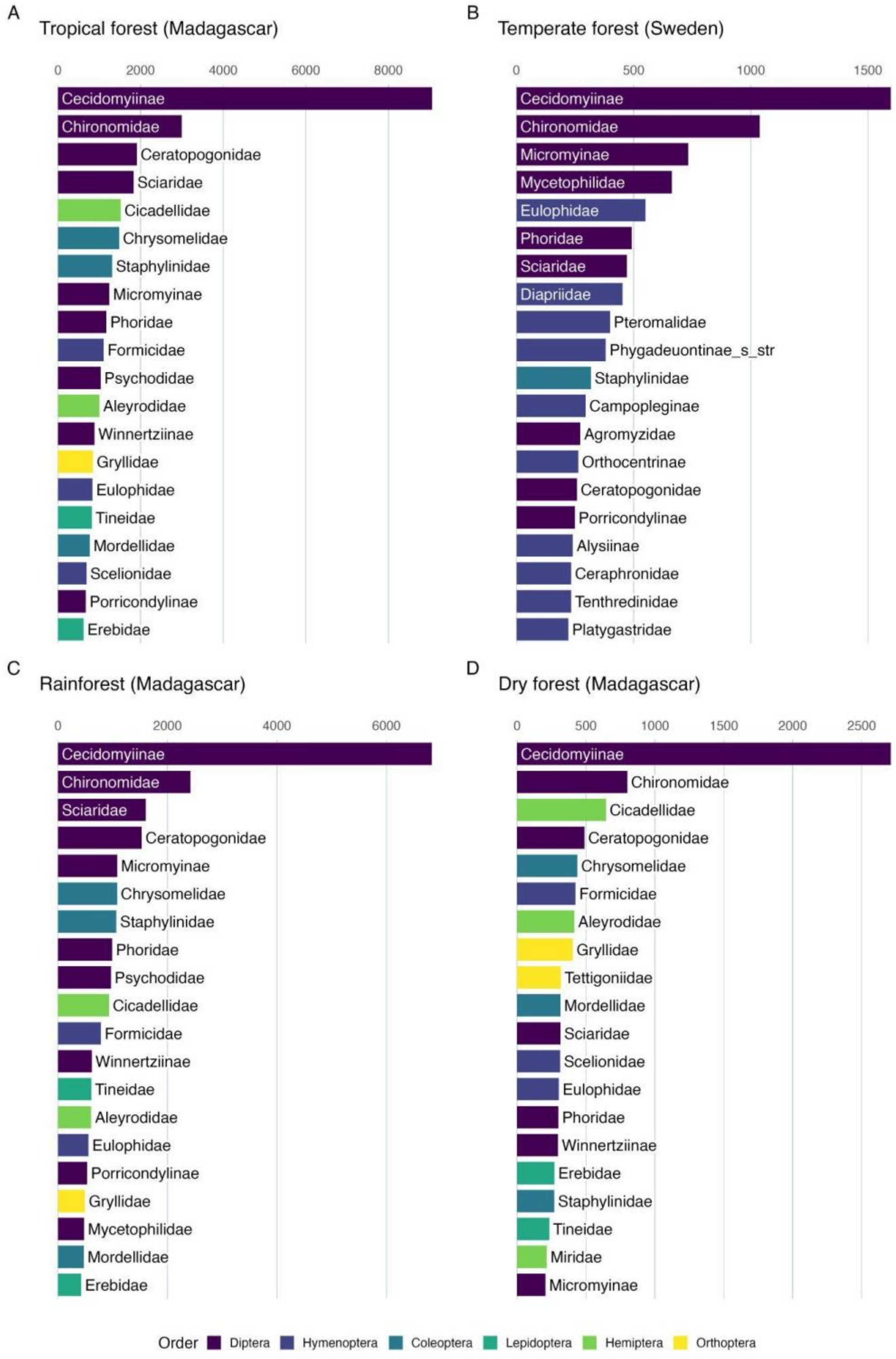


**Fig. S11. Most diverse family clades in Madagascan forests and the temperate reference forest**. Top 20 family clades (corresponding to subfamilies in Cecidomyiidae, Ichneumonidae and Braconidae, and to families in other taxa) are displayed for each forest type, ranked by the number of OTUs placed in them.

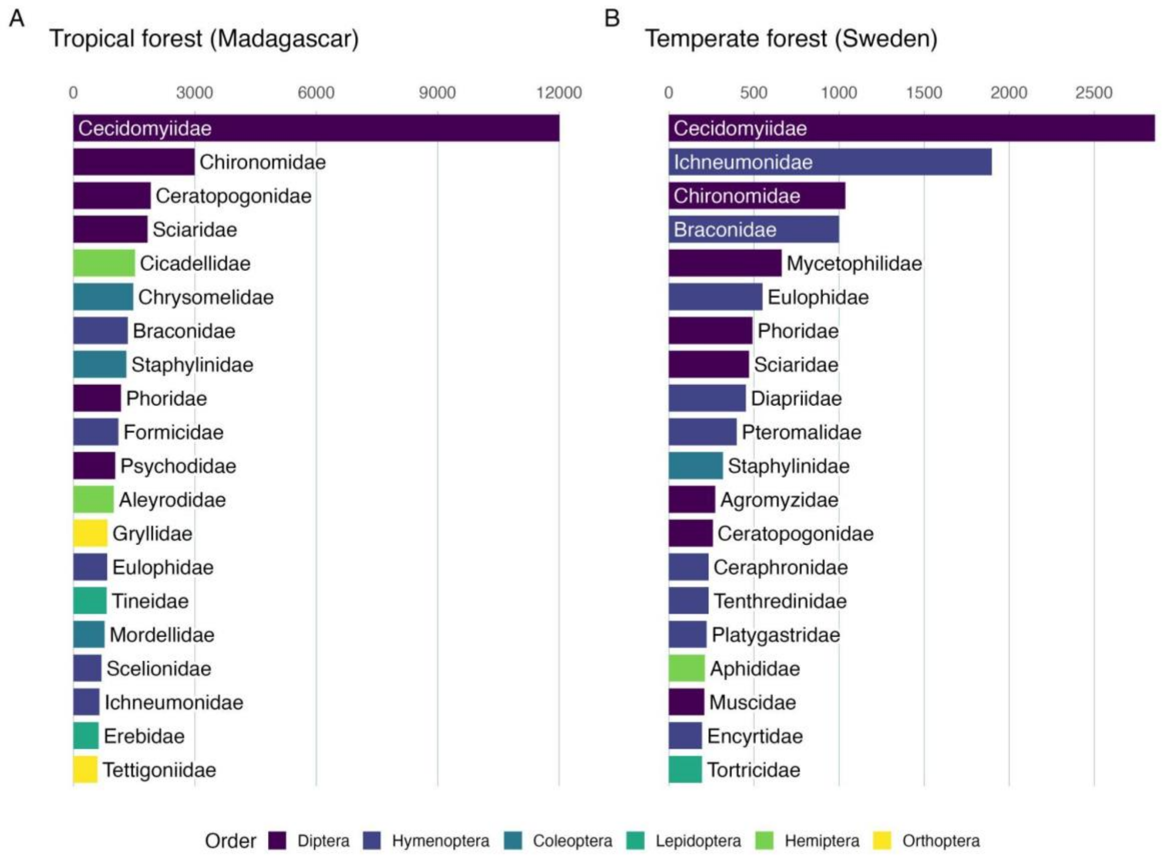


**Fig. S12. Most diverse families in the Madagascan forests and the temperate reference forest**. The top 20 families are displayed for each dataset, ranked by the number of OTUs placed in them.

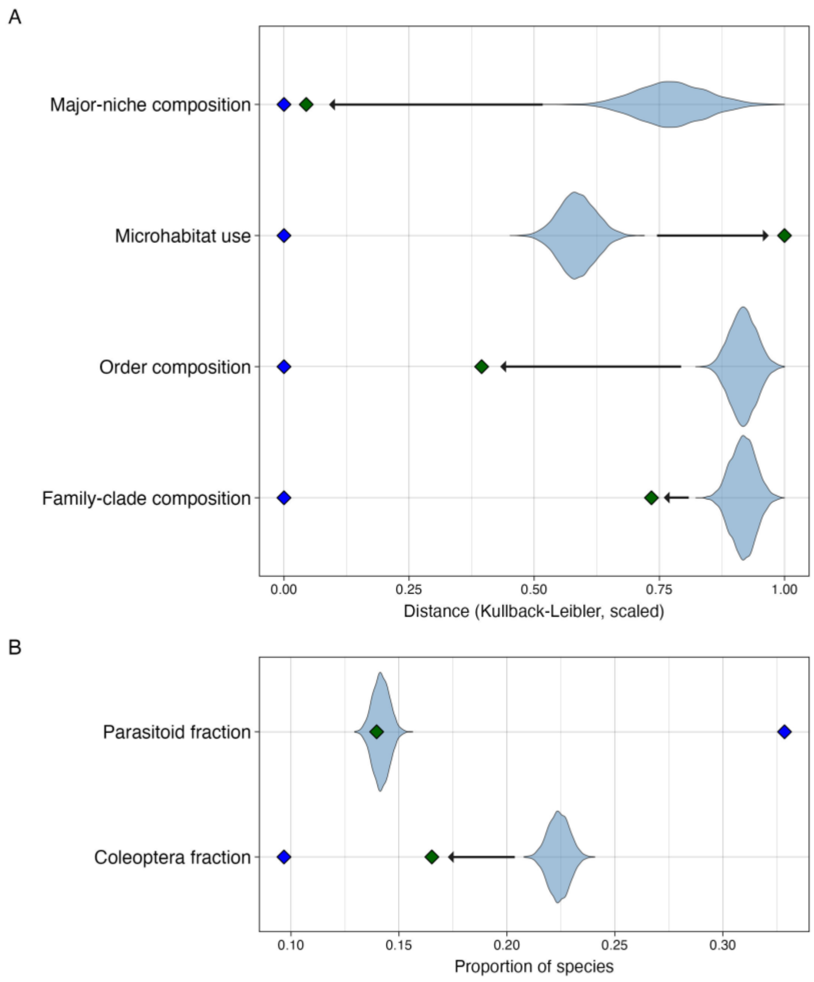


**Fig. S13. Diversification analysis of compositional features with no sampling bias correction**. We fitted a common time-dependent diversification model to all Madagascan radiations, conditioning on the age and the number of sampled OTUs of each radiation. Unlike Fig. 5 in the main text, we did not correct for any sampling bias among insect groups here. By simulating data from the fitted model, we tested whether observed compositional features (green diamonds) deviated from the expectations under neutral diversification of colonizing lineages and, if so, whether the composition evolved (arrows) to become more similar or more different to that of temperate forests (blue diamonds). The simulations (violin plots) are likely to represent compositional features of source faunas filtered through partly stochastic and partly deterministic selection of successful colonizers. (**A**) Compositional features measured in terms of relative Kullback-Leibler distance to temperate forests. (**B**) Fraction of parasitoids and beetles (Coleoptera).

**Comment**: Beetle lineages are diverse in terms of feeding niches and microhabitat use, explaining why the sampling bias against them has limited effect on most compositional features. However, the bias does affect the sampling of beetle species more than the sampling of beetle lineages colonizing Madagascar, changing the results of the analysis.

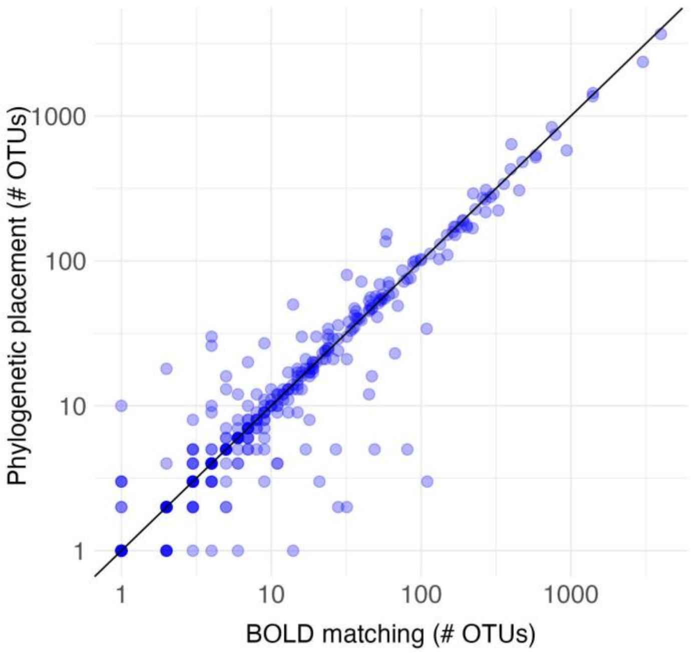


**Fig. S14. Validation of the phylogenetic placement methodology using Swedish data.** We compare the results of annotating OTUs to family by phylogenetic placement (with EPA-NG and GAPPA) on a reference tree with the results of annotation to family by matching to the BOLD reference library (with SINTAX).

**Comment**. For Swedish insects, the BOLD reference library has good coverage of the majority of families, resulting in good performance of annotation to higher taxa by matching sequences to the reference library. This is not the case for Madagascan insects, making phylogenetic placement in a reference tree more attractive. Here, we check that the phylogenetic placement methodology results in similar annotations to higher taxa for the Swedish data. Discrepancies include cases where there is disagreement in circumscription of families, or where families are not monophyletic, among other things. The discrepancies seem to primarily affect species-poor families.

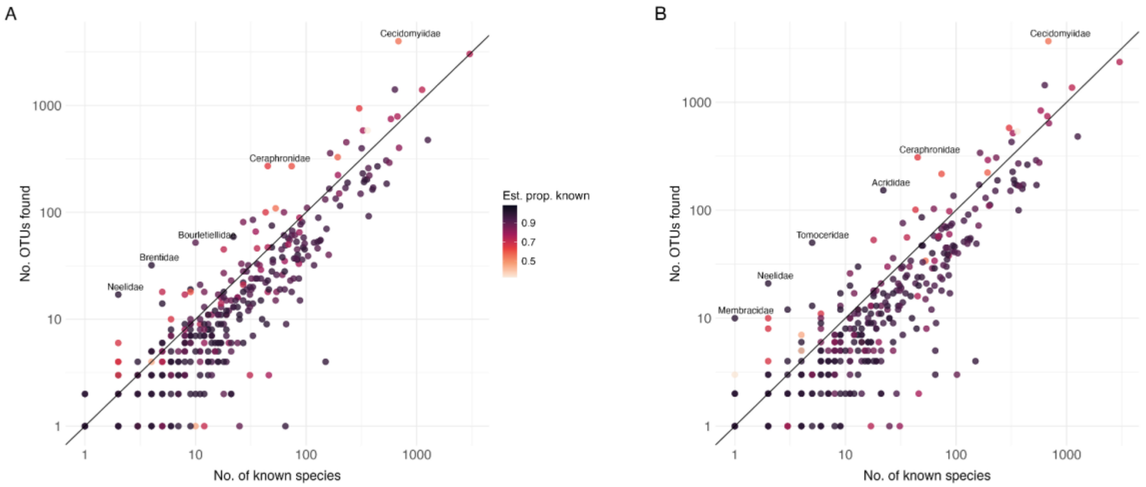


**Fig. S15. Validation of data quality using the Swedish insect fauna**. Correlation between the number of sampled OTUs (Malaise samples (lysates) and litter samples) per family and the number of known species in the Swedish insect fauna (Ronquist et al. 2020). (A) Data annotated to family by matching to the BOLD reference library using sintax. (B) Data annotated to family by placing OTUs in the reference tree developed for this study. The color reflects the estimated proportion of unknown species remaining to be discovered in each family (Ronquist et al. 2020). The line corresponds to the number of OTUs expected if all the known Swedish species were included in the sample; families are expected to fall below this line if the estimated proportion of known species is high.

**Comment**. Taxonomic annotation with the phylogenetic placement methodology used here (B), which is less sensitive to gaps in the reference data and thus more robust when analyzing poorly charted faunas, results in only a slight increase in false positives compared to matching to the BOLD reference library (A). Large numbers of ghost OTUs are still observed in some families that are well known (dark dots above the diagonal line), indicating that the noise filtering is ineffective for those taxa.

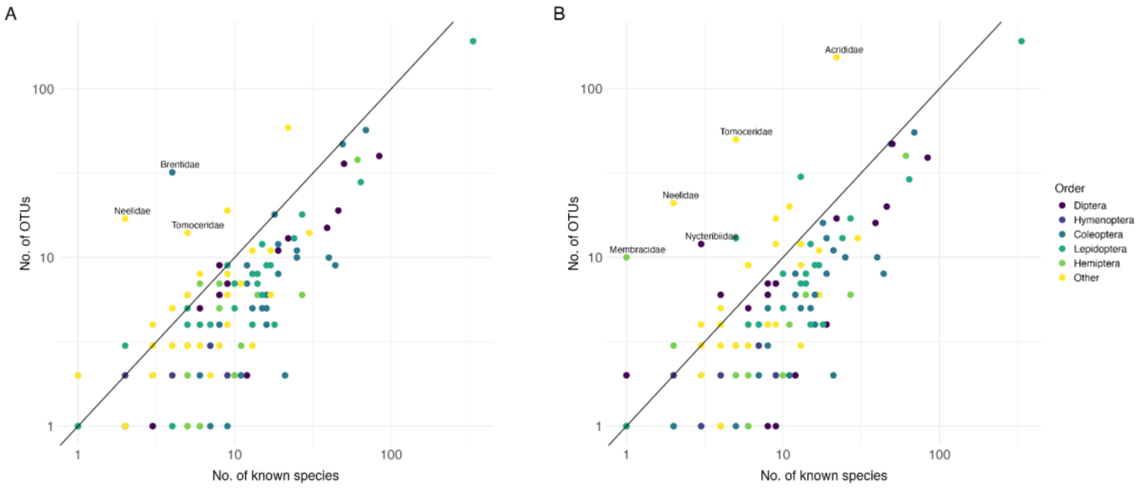


**Fig. S16. Validation of data quality for well-known families in the Swedish fauna**. Comparison of estimated number of OTUs for well-known families (estimated proportion of known species ≥ 0.99) in the Swedish fauna against the number of known species (diagonal line). The colour indicates the order to which each family belongs. (A) Filtered data annotated to family by matching to the BOLD reference library using SINTAX. (B) Filtered data annotated to family by placing OTUs in the reference tree developed for this study using EPA-NG and GAPPA, without using the BOLD reference library.

**Comment**: Despite the application of advanced noise filtering algorithms, a large number of spurious or ghost OTUs remain in a few families, such as Neelidae and Tomoceridae (Collembola). The problematic taxa tend to be relatively poor in species or belong to the minor orders. Matching to the BOLD reference library (A), which is generally fairly complete for the Swedish fauna analyzed here, helps in resolving some of these issues but is susceptible to others that do not cause problems in phylogenetic placement (B).

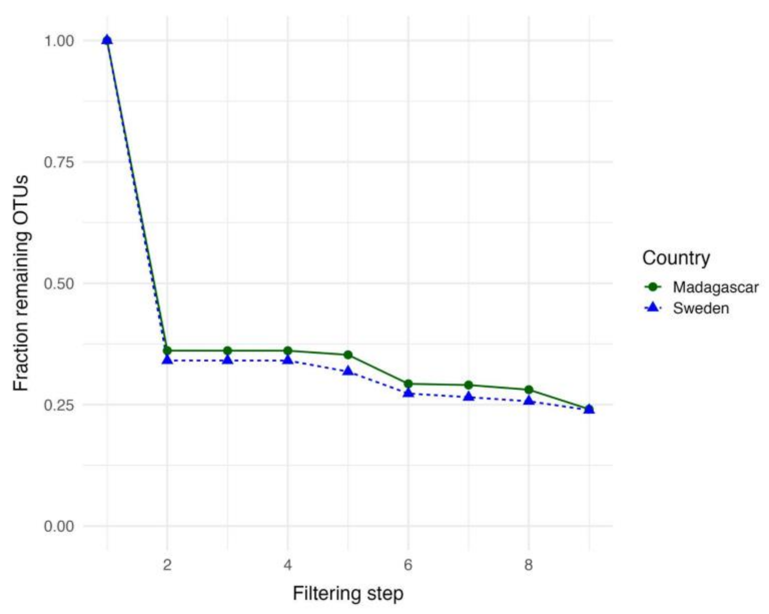


**Fig. S17. Performance of the filtering and annotation methodology on Swedish and Madagascan data**. Filtering steps: (1) Unfiltered clusters from ampliseq and DADA2 pipeline. (2) NEEAT filtering. (3) Removal of contaminants based on control samples. (4) Removal of spike-ins. (5) Restriction to hexapods. (6) Removal of uncertain phylogenetic placements (aLWR ≤ 0.5). (7) Removal of OTUs with SINTAX BOLD placements conflicting at the order level. (8) Removal of OTUs with VSEARCH placements conflicting at the phylum level. (9) Restriction to OTUs assigned to family clade.

**Comment**: The main filtering step, where most of the noise is removed, involves the removal of NUMTs, sequencing artefacts and other likely spurious OTUs by the NEEAT algorithm (step 2). Slightly fewer clusters are removed from the Madagascan data, presumably due to the shallower sampling per species, resulting in fewer artefacts being picked up. The shallower sampling also makes the recognition of artefacts by sample co-occurrence (the “echo” signal in NEEAT) less effective. Placing OTUs reliably in the reference tree (aLWR > 0.5; step 6) and to a family clade (step 9) is slightly less successful for Madagascar data than for Swedish data, presumably due to the reference tree not adequately covering the diversity of Madagascan insects

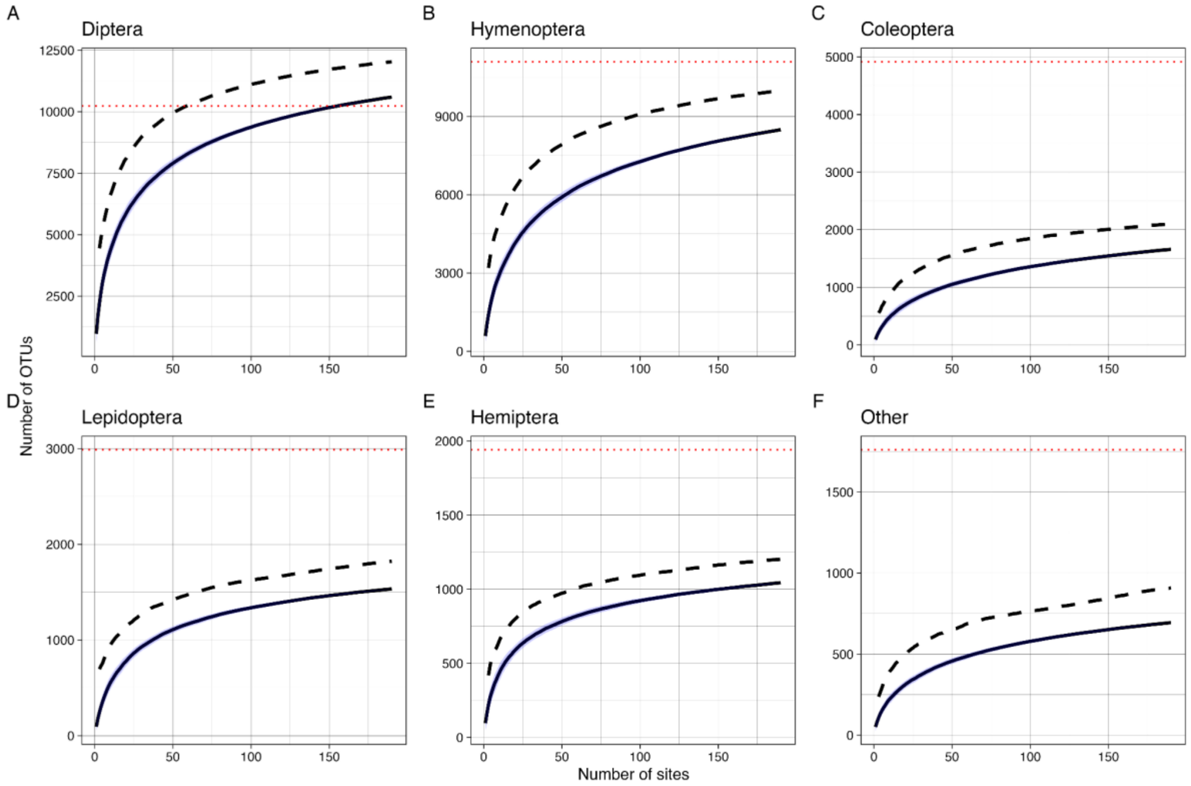


**Fig. S18. Species accumulation curves for the Swedish reference sample by order**. The data from all habitat types (forest and non-forest) are included. Curves are based on the mean of 100 permutations; the blue region marks values within one standard deviation from the mean. The dashed line represents average Chao2 extrapolations from the randomly permuted data. The red dotted line represents the estimated number of species in each order (Ronquist et al. 2020). (A) Diptera. (B) Hymenoptera. (C) Coleoptera. (D) Lepidoptera. (E) Hemiptera. (F) Other orders.

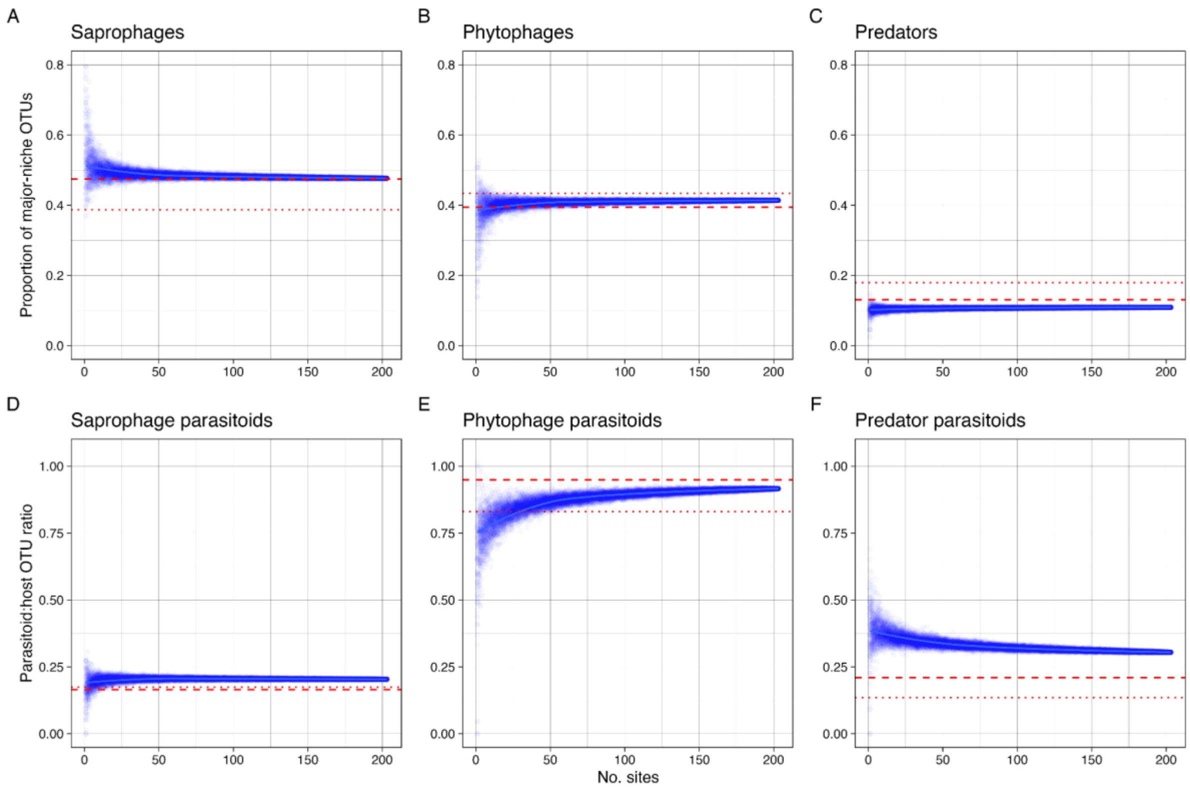


**Fig. S19. Spatial accumulation of niche composition across Swedish sites**. The points represent 100 jackknife samples for each number of sites. The data include all sampled habitats (forest and non-forest). The red dotted lines indicate the taxonomic composition of the true fauna based on a previously published estimate (Ronquist et al. 2020); the red dashed lines indicate the revised estimate based on correcting the fraction of unknown species in Diptera and Hymenoptera. (A–C) Proportion of OTUs belonging to the three major feeding niches. (A) Saprophages. (B) Phytophages. (C) Predators. (D–F) Parasitoid:host OTU ratios for each of the three main communities. (D) Saprophage parasitoids. (E) Phytophage parasitoids. (F) Predator parasitoids.

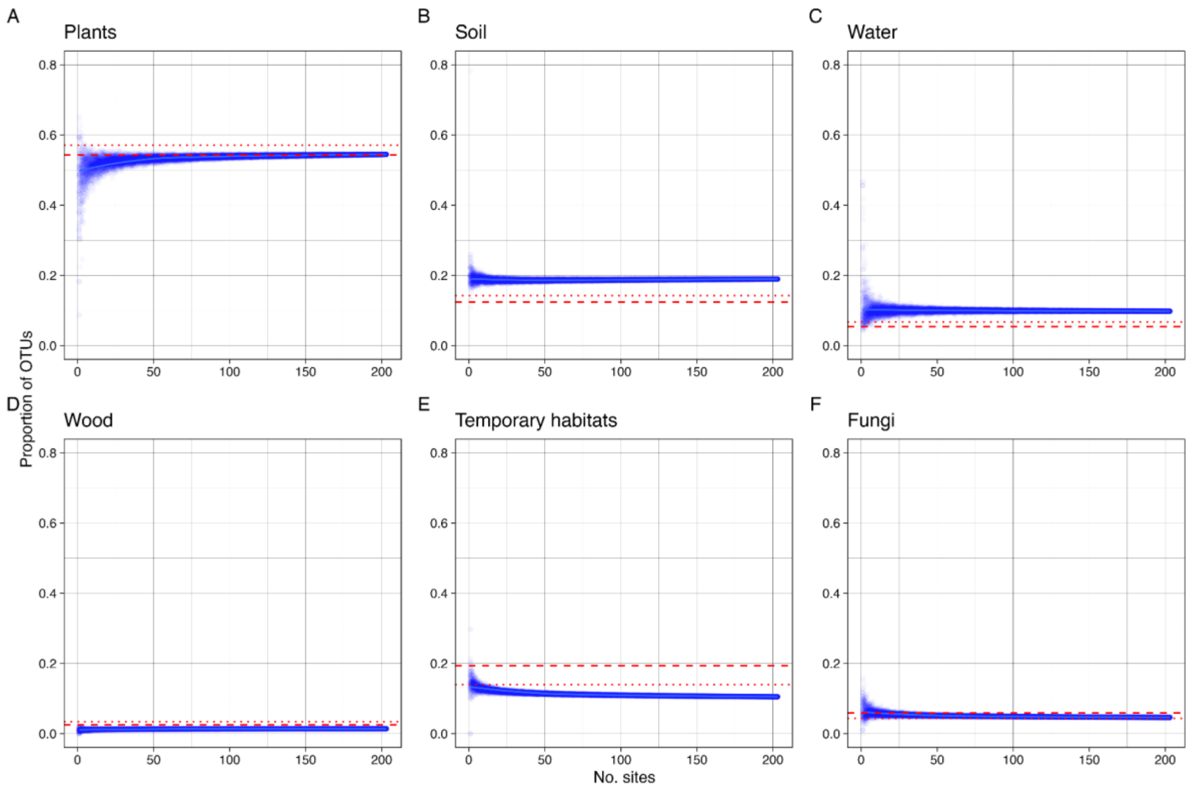


**Fig. S20. Spatial accumulation of microhabitat composition across sites**. The points represent 100 jackknife samples for each number of sites. The data include all sampled habitats (forest and non-forest). The red dotted lines indicate the taxonomic composition of the true fauna based on a previously published estimate (Ronquist et al. 2020); the red dashed lines indicate the revised estimate based on correcting the fraction of unknown species in Diptera and Hymenoptera. (A) Species developing in or on plants. (B) Species developing in soil, litter or on the ground. (C) Species developing in water. (D) Species developing in wood. (E) Species developing in temporary microhabitats, such as carrion or dung. (F) Species developing in or on fungi.

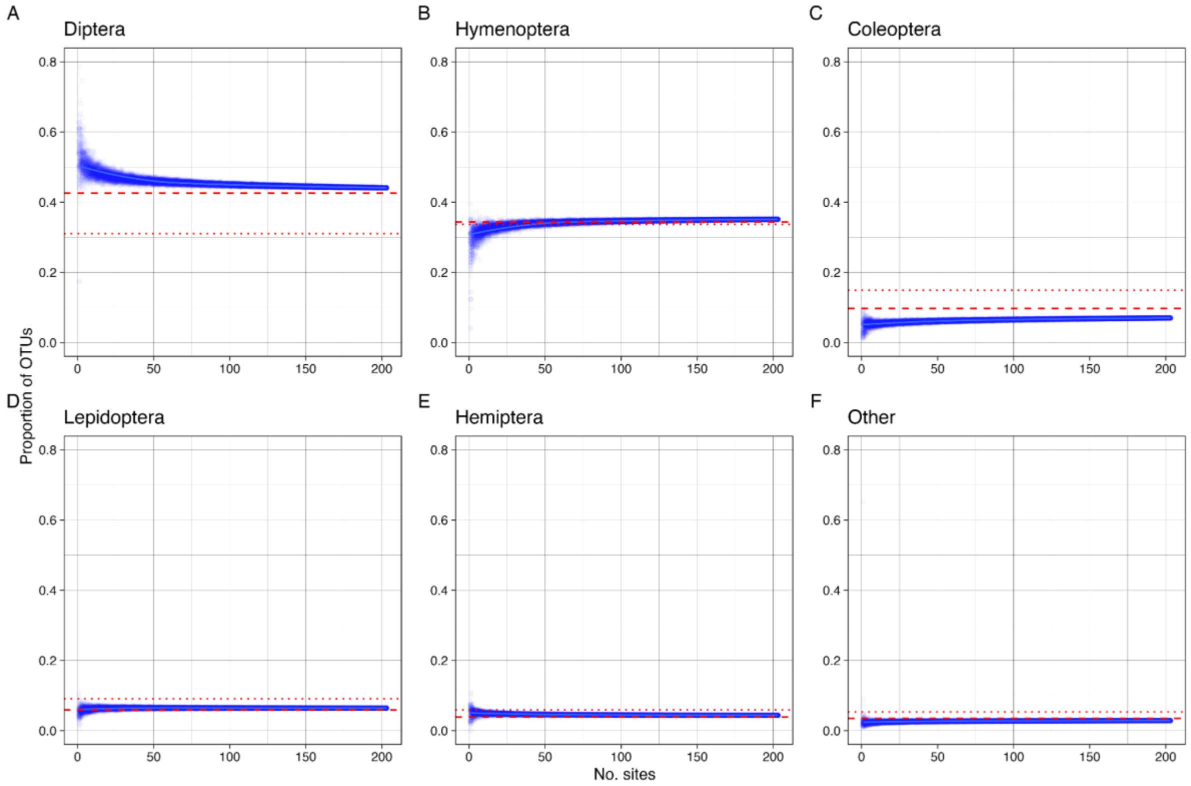


**Fig. S21. Spatial accumulation of taxonomic composition across Swedish sites**. The points represent 100 jackknife samples for each number of sites. The data include all sampled habitats (forest and non-forest). The red dotted lines indicate the taxonomic composition of the true fauna based on a previously published estimate (Ronquist et al. 2020); the red dashed lines indicate the revised estimate based on correcting the fraction of unknown species in Diptera and Hymenoptera. (A) Diptera. (B) Hymenoptera. (C) Coleoptera. (D) Lepidoptera. (E) Hemiptera. (F) Remaining hexapod orders.

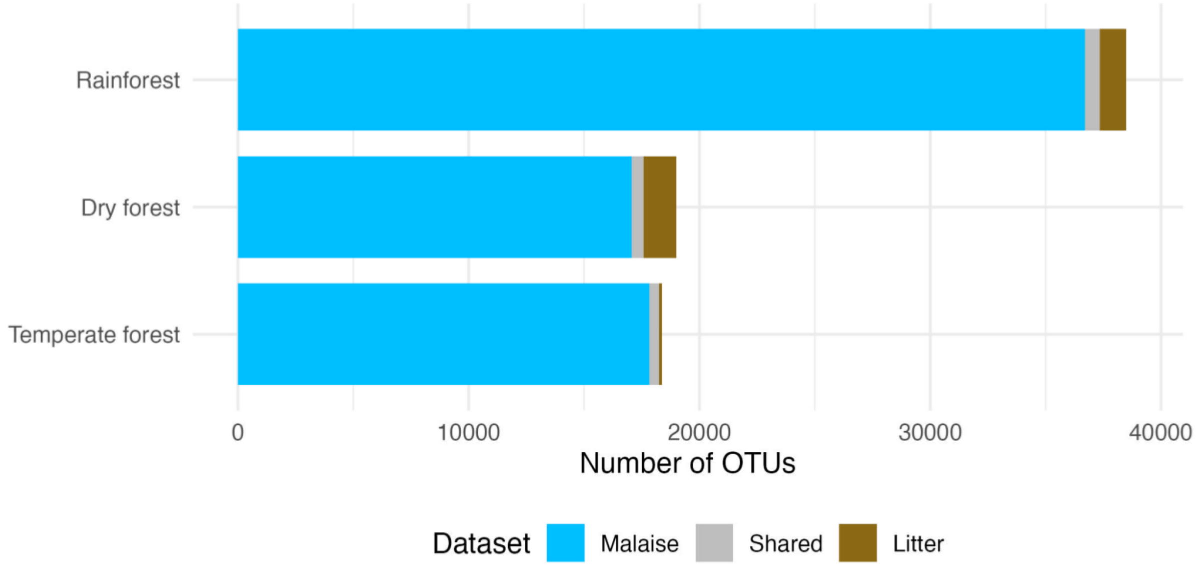


**Fig. S22. Contribution of different sampling methods (Malaise traps, litter samples, and shared) to the OTUs retrieved.** The Madagascan and Swedish data are presented by forest type: Madagascan rainforest and dry forest, and Swedish temperate forest.

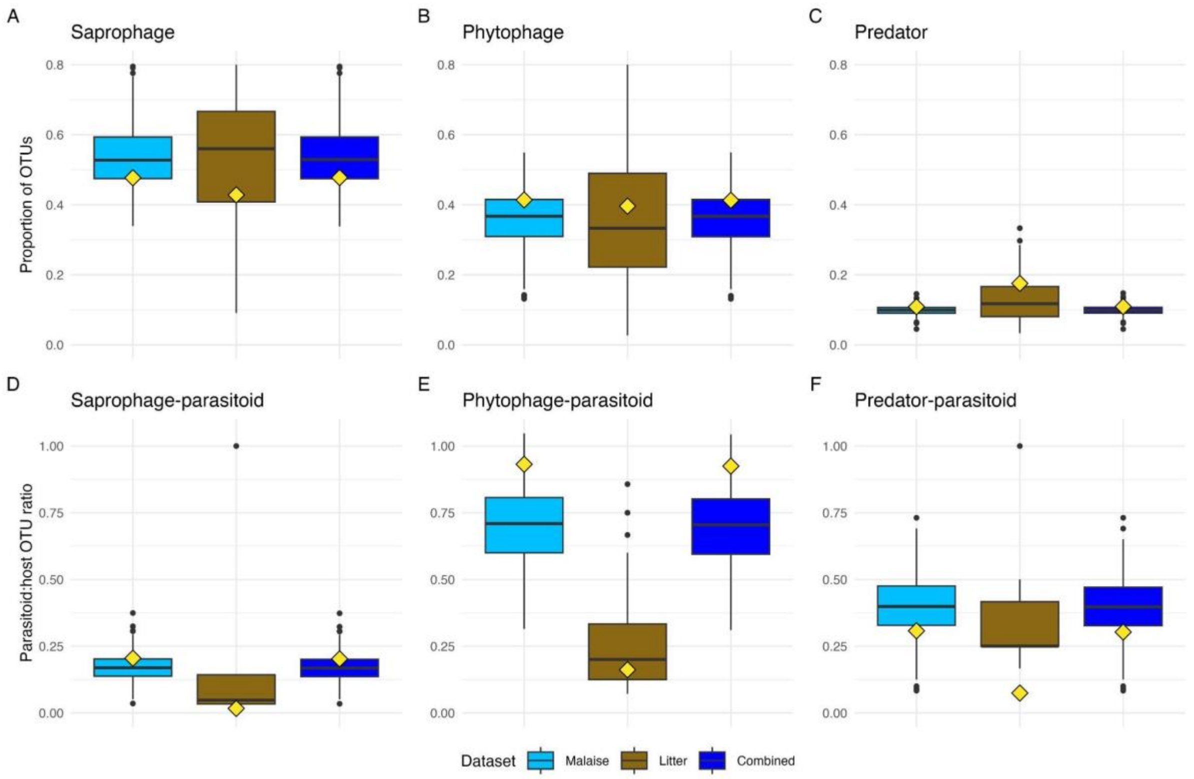


**Fig. S23. Contribution of different sampling methods (Malaise traps, litter samples, and combined) to the estimates of the niche composition of the Swedish fauna**. Boxplots summarize the distribution of values across 12-month samples from each site. The red diamonds indicate the values to which the accumulation curves converge, and the distance between each diamond and the mean of the corresponding box plot indicates the rate at which the value changes with additional spatial sampling. (**A–C**) Proportion of OTUs belonging to the three major feeding niches for each trap type and combined. (**D–F**) Ditto for parasitoid to host OTU ratios.

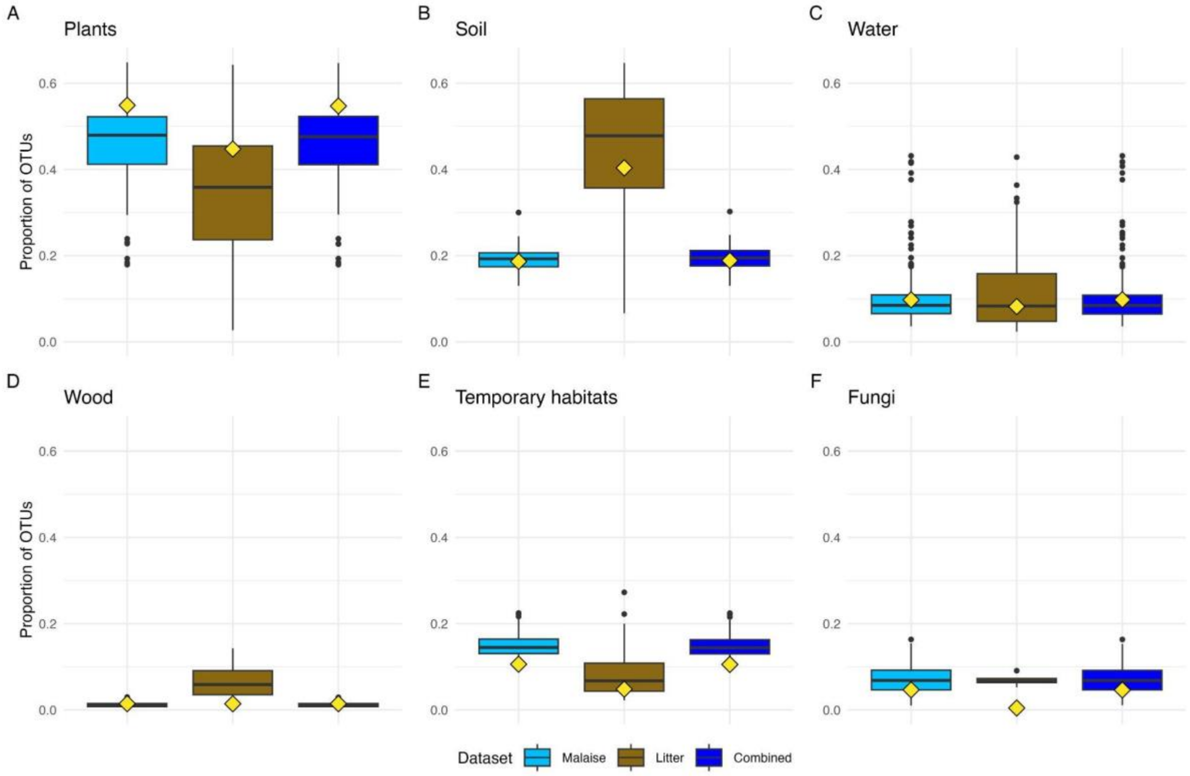


**Fig. S24. Contribution of different sampling methods (Malaise traps, litter samples, and combined) to the estimates of the microhabitat composition of the Swedish fauna**. Boxplots summarize the distribution of values across 12-month samples from each site. The red diamonds indicate the values to which the accumulation curves converge, and the distance between each diamond and the mean of the corresponding box plot indicates the rate at which the value changes with additional spatial sampling.

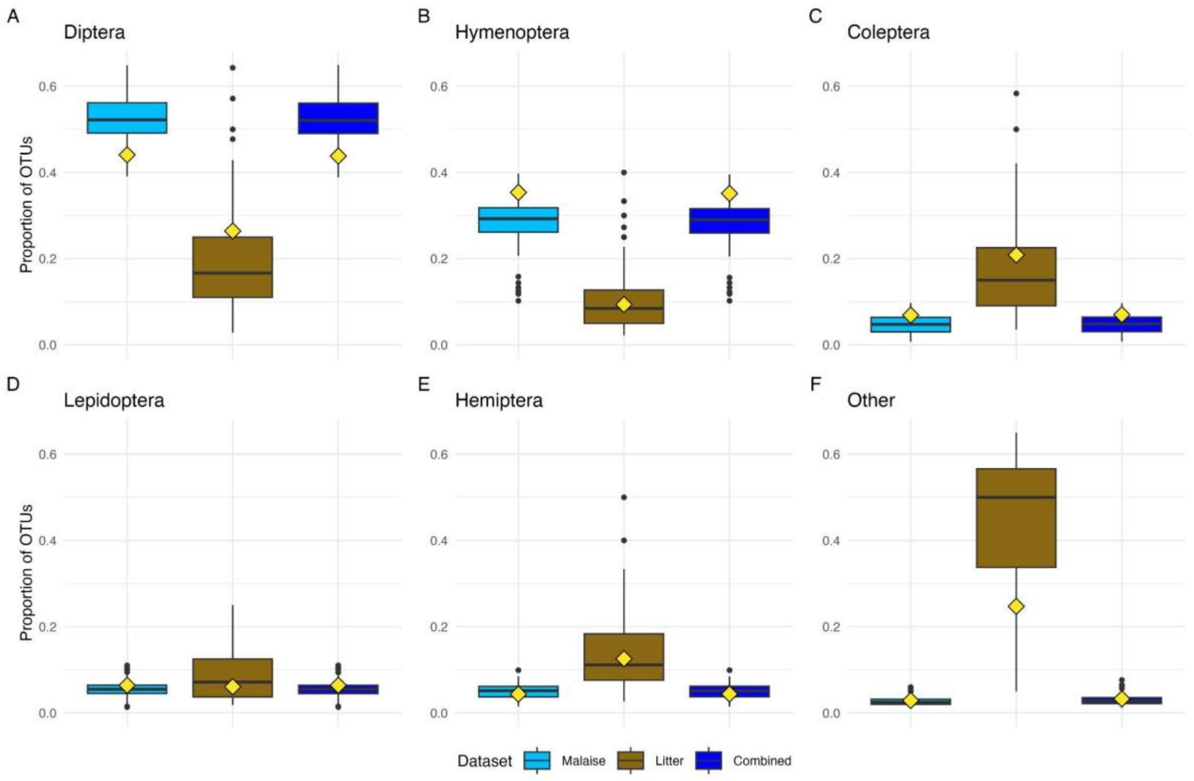


**Fig. S25. Contribution of different sampling methods (Malaise traps, litter samples, and combined) to the estimates of the taxonomic composition of the Swedish fauna**. Boxplots summarize the distribution of values across 12-month samples from each site. The red diamonds indicate the values to which the accumulation curves converge, and the distance between each diamond and the mean of the corresponding box plot indicates the rate at which the value changes with additional spatial sampling.

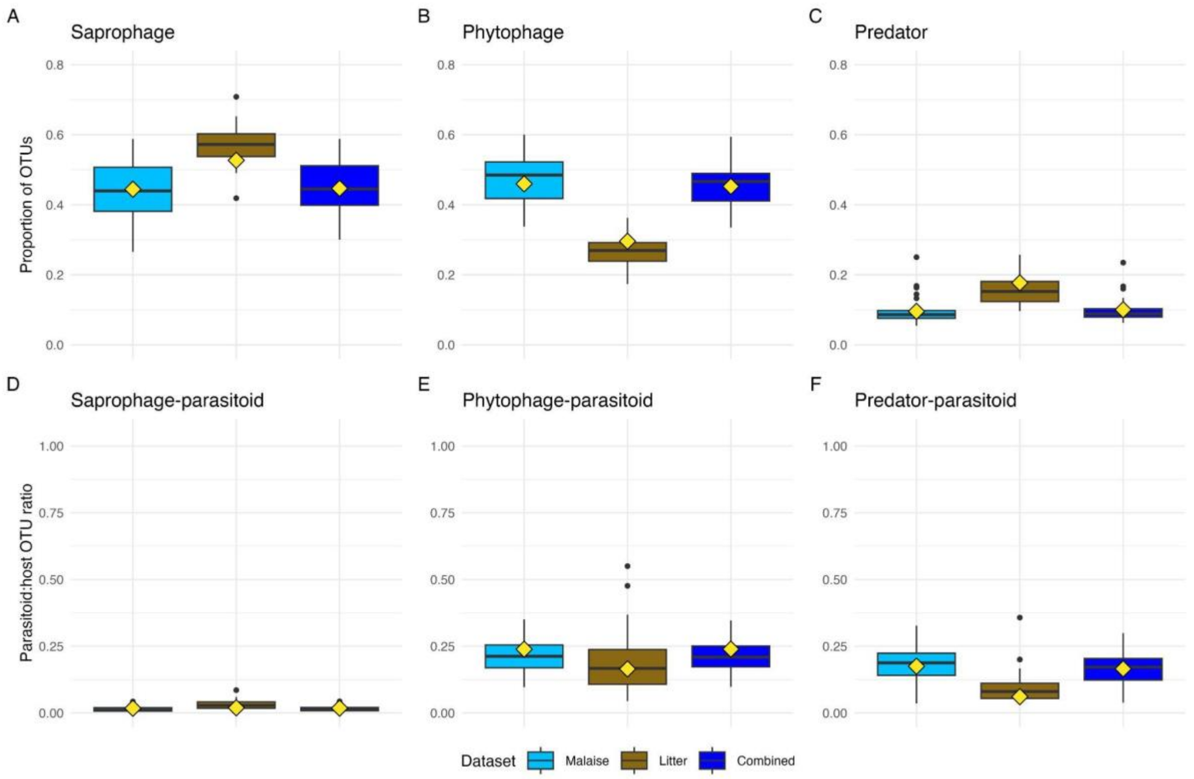


**Fig. S26. Contribution of different sampling methods (Malaise traps, litter samples, and combined) to the estimates of the niche composition of the Madagascan fauna**. Boxplots summarize the distribution of values across 12-month samples from each site. The red diamonds indicate the values to which the accumulation curves converge, and the distance between each diamond and the mean of the corresponding box plot indicates the rate at which the value changes with additional spatial sampling. (**A–C**) Proportion of OTUs belonging to the three major feeding niches. (**D–F**) Parasitoid to host OTU ratios for each of the three main communities.

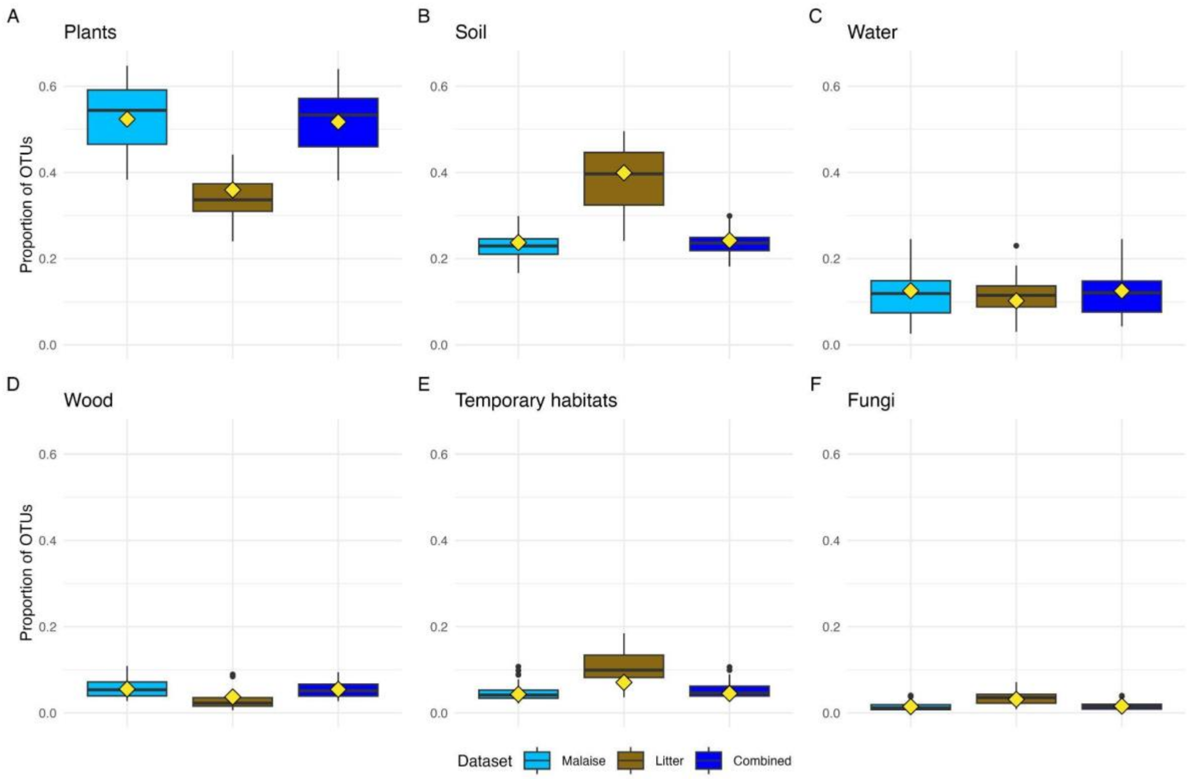


**Fig. S27. Contribution of different sampling methods (Malaise traps, litter samples, and combined) to the estimates of the microhabitat composition of the Madagascan fauna**. Boxplots summarize the distribution of values across 12-month samples from each site. The red diamonds indicate the values to which the accumulation curves converge, and the distance between each diamond and the mean of the corresponding box plot indicates the rate at which the value changes with additional spatial sampling.

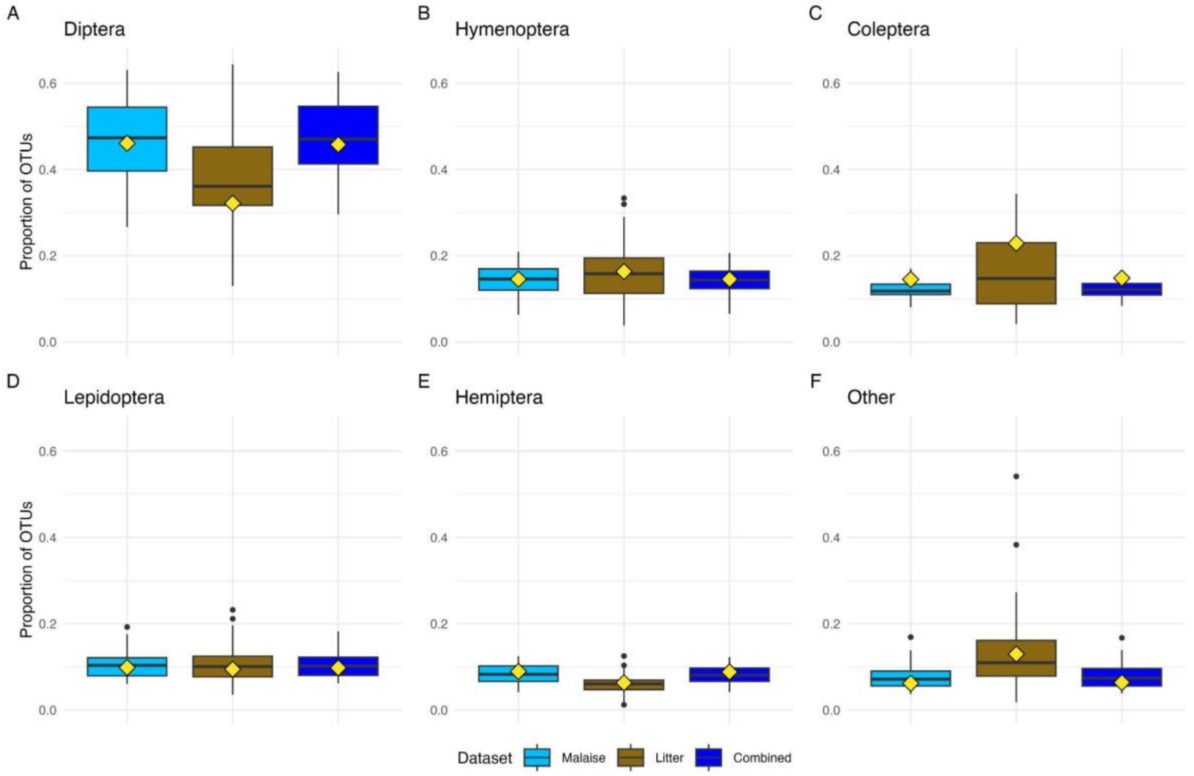


**Fig. S28. Contribution of different sampling methods (Malaise traps, litter samples, and combined) to the estimates of the taxonomic composition of the Madagascan fauna**. Boxplots summarize the distribution of values across 12-month samples from each site. The red diamonds indicate the values to which the accumulation curves converge, and the distance between each diamond and the mean of the corresponding box plot indicates the rate at which the value changes with additional spatial sampling.

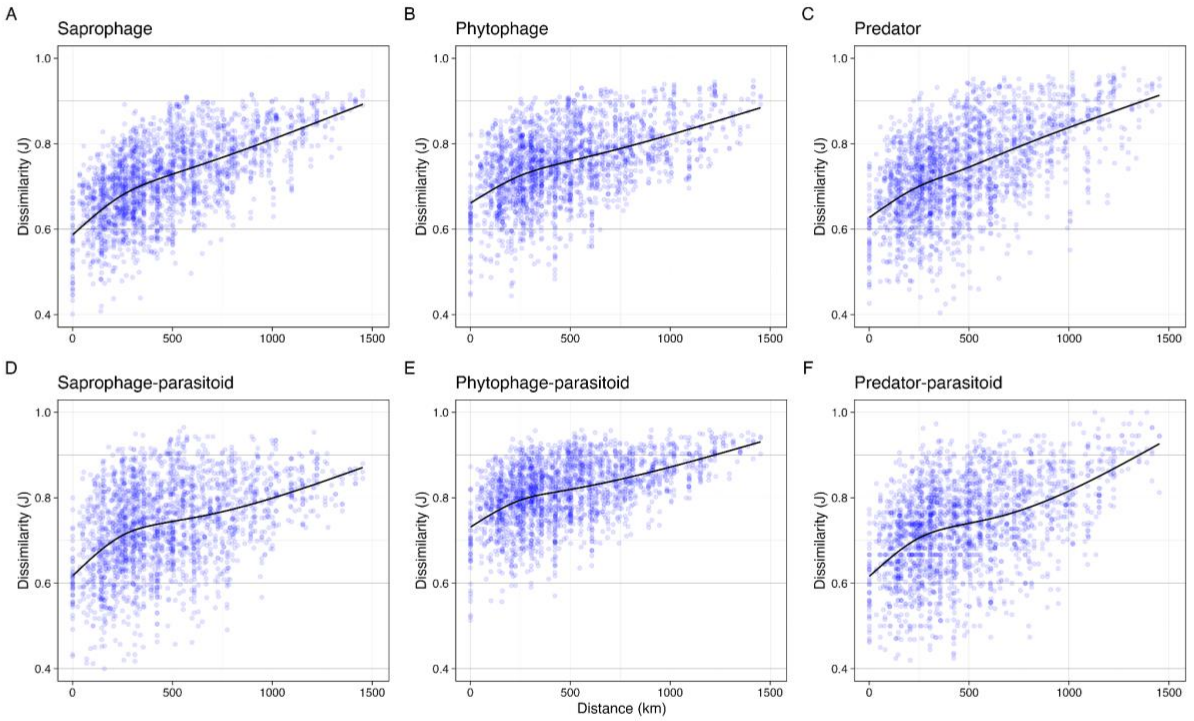


**Fig. S29. Spatial turnover of main feeding niches in the temperate forest sample**. Turnover is expressed as Jaccard dissimilarity between pairs of trap sites. The fitted line is generated from a monotonic generalized additive model with five knots.
